# Metabolic connectivity has greater predictive utility for age and cognition than functional connectivity

**DOI:** 10.1101/2024.08.15.608184

**Authors:** Hamish A. Deery, Emma Liang, Chris Moran, Gary F. Egan, Sharna D. Jamadar

## Abstract

Recently developed high temporal resolution functional [18F]-fluorodeoxyglucose positron emission tomography (fPET) offers promise as a method for indexing the dynamic metabolic state of the brain *in vivo* by directly measuring a timeseries of metabolism at the post-synaptic neuron. This is distinct from functional magnetic resonance imaging (fMRI) that reflects a combination of metabolic, haemodynamic and vascular components of neuronal activity. The value of using fPET to understand healthy brain ageing and cognition over fMRI is currently unclear. Here we use simultaneous fPET/fMRI to compare metabolic and functional connectivity and test their predictive ability for ageing and cognition. Whole-brain fPET connectomes showed moderate topological similarities to fMRI connectomes in 40 younger (mean age 27.9 years; range 20-42) and 46 older (mean 75.8; 60-89) adults. There were more age-related within- and between-network connectivity and graph metric differences in fPET than fMRI. fPET was also associated with performance in more cognitive domains than fMRI. These results suggest that ageing is associated with a reconfiguration of metabolic connectivity that differs from haemodynamic alterations. We conclude that metabolic connectivity has greater predictive utility for age and cognition than functional connectivity and that measuring glucodynamic changes has promise as a biomarker for age-related cognitive decline.

## Introduction

The history of neuroscientific inquiry into brain function that spans several centuries has vastly expanded our understanding of behaviour and cognition.^1^ One important area of study has been brain energetics because of the obligatory role that metabolism plays in brain function.^2^ A range of experimental methods and tools have been used to study metabolic pathways in the brain and have revealed a tight coupling between neuronal activity and metabolism at multiple temporal and spatial scales.^3-5^ Increases in the brain’s utilisation of glucose matches synaptic activity^4, 6-9^, and at the cellular level, neuronal glucose oxidation is directly related to the frequency of spike activity and neurotransmitter cycling rates.^10, 11^ Functional-metabolic associations are also evident at a regional level.^2 12^ Early studies using [18F]-fluorodeoxyglucose positron emission tomography (FDG-PET) compared images from rest and task conditions and showed an increase in glucose metabolism in task-relevant visual^13^ and motor^14^ regions.

The oxidation of glucose in the brain in response to neuronal activity is also accompanied by an increase in oxygen consumption and local cerebral blood flow, leading to the increase in oxygenation that produces the blood oxygenated level dependent (BOLD) signal that underpins fMRI.^15, 16^ fMRI has provided ground-breaking knowledge about the functional architecture of the brain, and shown that cognitive functions are dependent on coherent activity between brain regions in large-scale networks.^17, 18^ However, fMRI provides only one index of the coherent neurophysiological signals that underlie the brain’s functional network. Importantly, information transfer across the brain arises from multiple physiological processes, including haemodynamic, electrophysiological and molecular processes. Molecular imaging techniques, including high temporal resolution functional FDG-PET (fPET), allow for the dynamic within-subject measurement of molecular processes, such as glucose metabolism.^19-28^ The coherence of glucose signals across the brain indexed with FDG is termed ‘metabolic connectivity’.^29, 30^ We have reported that metabolic connectivity provides important and complementary insights into brain function relative to BOLD-fMRI derived networks.^20, 21, 29^

Given the large individual and societal costs of age-related cognitive decline, brain changes in normative ageing and neurodegenerative conditions are a crucial focus of neuroscientific enquiry.^31^ fMRI has revealed that large-scale functional brain networks in ageing are characterised by a reduction of within-network connectivity, an increase in between-network connectivity and a reduction in the efficiency of connections across the brain, particularly connections between associative networks.^32^ Mechanistically, neuroimaging studies utilising the BOLD signal and glucose metabolism have identified shared but important unique components that have implications for their interpretation.^30^ The BOLD signal reflects changes in deoxyhemoglobin driven by localised changes in blood flow and blood oxygenation. However, the interpretation of altered BOLD signals between groups can be challenging, as any alteration in the coupling in cerebral blood flow, volume or oxygen metabolism, unrelated to changes in neuronal activity, will result in an altered BOLD signal.^16, 33, 34^ For example, the coupling between blood flow, blood volume and metabolism are differentially altered in healthy ageing and hence age-related changes of the BOLD signal may reflect alterations in one or more neurovascular component in older adults.^9, 35-37^

In contrast, FDG-PET signals in the brain are proportional to glucose metabolism,^38^ and can therefore provide a complementary, more direct and quantifiable measure of neuronal activity than the BOLD signal. Whole-brain simultaneous fPET/fMRI in younger adults has revealed that metabolic connections are predominantly between frontal and parietal regions, whereas fMRI connections are more widespread across the brain.^29, 39, 40^ Moreover, fPET and fMRI connectomes in younger adults are differentially related to executive function.^24^ Using fPET, we recently showed differences in the metabolic connectomes of older and younger adults.^41^ We found lower metabolic connectivity in older than younger adults between the frontal and temporal regions, and more globally integrated posterior hub regions in older adults. Qualitatively, these results contrast with the fMRI literature which shows increased connectivity between the frontal and temporal regions and a posterior to anterior shift in ageing.^32^

Here we use simultaneous fPET/fMRI to compare metabolic and functional connectivity and test their predictive utility for healthy ageing and cognition. Given that the FDG and BOLD signals index some shared physiological processes, we hypothesise that there will be moderate similarity in the topology of fPET and fMRI connectomes.^29^ Secondly, following our previous qualitative observations of fPET and fMRI differences,^32, 41^ we hypothesise that differences between younger and older adults in network connectivity will be more widespread in fMRI than fPET, with lower within- and higher between-network connectivity in older compared to younger adults, particularly for associative networks. Thirdly, that fPET connectomes will show lower local and global efficiency for older than younger adults in line with large-scale functional network alterations reported in fMRI studies.^32^ Finally, we hypothesise that fPET and fMRI graph metrics will both predict cognitive performance, and that the metrics will differentially associate with cognition across domains, in line with our previous work in younger adults.^24^

## Materials & Methods

### Ethical Considerations

All methods were reviewed by the Monash University Human Research Ethics Committee, following the Australian National Statement of Ethical Conduct in Human Research (2023). Administration of ionising radiation was approved by the Monash Health Principal Medical Physicist, following the Australian Radiation Protection and Nuclear Safety Agency Code of Practice (2005). For participants aged over 18-years, the annual radiation exposure limit of 5mSv applies, and the effective dose in this study was 4.8mSv.

### Participants

Ninety participants were recruited from the local community. An initial screening interview ensured that participants did not have a history of diabetes, neurological or psychiatric illness and were not taking psychoactive medication. Participants were also screened for claustrophobia, non-MR compatible implants and a clinical or research PET scan in the previous 12 months. Women were screened for current or suspected pregnancy. Participants received a $100 voucher for participating in the study. Four participants were excluded for further analyses due to either excessive head motion (N=2) or incomplete PET data (N=2).

### Data Acquisition

Participants completed an online demographic and lifestyle questionnaire and a cognitive test battery. Briefly, the following cognitive measure were used (see Supplement for details): delayed recall and a recognition discrimination index from the Hopkins Verbal Learning Test; length of longest correct series of forward and backward recall from a digit span test to index working memory; the proportion of correct trials and mean reaction time in a task switching test to index cognitive control and flexibility; the probability of responding and reaction time in a stop signal task to measure response inhibition; and number of correct responses and seconds per correct response in a digit substitution task to measure visuospatial performance and processing speed.

Participants underwent a 90-minute simultaneous MR-PET scan in a Siemens (Erlangen) Biograph 3-Tesla molecular MR scanner. At the beginning of the scan, half of the 260 MBq FDG tracer was administered via the left forearm as a bolus, the remaining 130 MBq of the FDG tracer dose was infused at a rate of 36ml/hour over 50 minutes.^42^ Non-functional MRI scans were acquired during the first 12 minutes, including a T1 3D MPRAGE and T2 FLAIR. Thirteen minutes into the scan, list-mode PET and T2* EPI BOLD-EPI sequences were initiated. A 40-minute resting-state scan was undertaken in naturalistic viewing conditions. Full details of the acquisition parameters are provided in Supplementary Information.

### MRI and PET Pre-Processing

For the T1 images, the brain was extracted in Freesurfer; quality of the pial/white matter surface was manually checked, corrected and registered to MNI152 space using Advanced Normalization Tools (ANTs). For the BOLD-fMRI data, T2* images were brain extracted (FSL BET), unwarped and motion corrected with six rotation and translation parameters (FSL MCFLIRT), temporally detrended, normalised to MNI space and smoothed at 8mm FWHM.^43^ Framewise displacement was calculated for each participant to check for excessive head motion (mean and % of frames > .3mm).

The list-mode PET data for each subject were binned into 344 3D sinogram frames of 16s intervals. Attenuation was corrected via the pseudo-CT method for hybrid PET-MR scanners.^44^ Ordinary Poisson-Ordered Subset Expectation Maximization algorithm (3 iterations, 21 subsets) with point spread function correction was used to reconstruct 3D volumes from the sinogram frames. The reconstructed DICOM slices were converted to NIFTI format with size 344 × 344 × 127 (size: 1.39 × 1.39 × 2.03 mm^3^) for each volume. All 3D volumes were temporally concatenated to form a single 4D NIFTI volume. After concatenation, the PET volumes were motion corrected using FSL MCFLIRT,^43^ with the mean PET image used to mask the 4D data. PET images were corrected for partial volume effects using the modified Müller-Gartner method^45^ implemented in Petsurf.

Participants’ T1 images and pre-processed fMRI and fPET timeseries data were loaded to the CONN toolbox in Matlab.^46^ For the fPET data, the first 10 minutes of the timeseries was excluded to remove the initial radiotracer uptake and to reflect the time corresponding to the onset of a stable signal.^21^ The timeseries were denoised using the default CONN pipeline, regressing out the potential confounding from white matter and CSF. The fMRI timeseries data was bandpass frequency filtered between 0.01 Hz and 0.1 Hz. A low pass filter (.0625 Hz) was applied to the fPET data to filter out high frequency noise in the PET signal.^25^ Regions of interest were generated for the Schaefer atlas.^47^ As it is currently unclear if fMRI-derived ‘functional’ atlases are transferable to metabolic connectomes,^19, 28, 29, 40^, we also applied an anatomical parcellation using the 106 cortical and subcortical regions of the Harvard Oxford atlas, excluding the cerebellum. The analyses from the main manuscript are repeated with the Harvard Oxford atlas and are reported in the Supplementary Information Section 3.

The following graph theory metrics were calculated in CONN at the top 30% of network edges: Global efficiency, local efficiency, betweenness centrality and degree for each node and at the whole brain level (see Supplement for definitions). The top 30% of edges was chosen as it is able to reliably identify age group differences in fPET.^41^ The graph metrics were also calculated at the Schaefer network level for the fPET and fMRI data. Because the regions within the networks differ in cortical volume, the network values could not simply be averaged. Rather, each regional value was weighted by the percentage its volume represented within the network.

### Similarity of fPET and fMRI Connectomes in the Whole Sample

To test the first hypothesis that there will be moderate similarity in the topology of fPET and fMRI connectomes, we qualitatively assessed the fPET and fMRI connectomes in the whole sample. We also calculated the DICE coefficient between the connectomes at costs from 10% and 90% of connections to quantify their spatial similarity. To further explore the degree of similarity between the fPET and fMRI connectomes, we calculated the covariance between them, by calculating the Pearson’s correlation of each connection in the matrices across participants.

### Analyses of Modality and Age Group Differences

#### Metabolic Connectomes

To test the second hypothesis that differences between younger and older adults in network connectivity will be more widespread in fMRI than fPET, we qualitatively assessed the connectomes of older and younger adults and compared them across modalities. The connectomes were generated by reverse transforming the individual participant matrices of Fisher transformed z-values back to r-values, and calculating the average for each node separately for the younger and older adult groups for both the fPET and fMRI data.

#### Within- and Between-Network Connectivity

To quantify the connectivity in the younger and older adult connectomes, within- and between-network connectivity was calculated. *Within-network connectivity* was calculated as the average Fisher transformed z-value of all regions within the network. *Between-network connectivity* was calculated as the average of the regional connections within that network and the regions in all other networks. We used independent sample *t*-tests (p-FDR < .05, one tailed) to test the hypothesis that older adults will have lower within- and higher between-network connectivity than younger adults, particularly in the frontal and temporal regions.

#### Age Differences in fPET and fMRI Graph Metrics

A series of *t*-tests was used to test the third hypothesis that the graph metrics derived from the fPET and fMRI connectomes will show lower global and local efficiency and centrality and degree in older than younger adults. First we compared younger and older adults on the graph metrics at the region level. Second, we compared younger and older adults on the graph metrics at the network level. Each series of *t*-tests for a graph metric was tested at p-FDR < .05, one tailed.

### Relative Predictive Strength of fPET and fMRI Graph Metrics for Age Group

Discriminant function analysis was used to classify participants into their predefined age groups based on the fPET and fMRI whole brain graph metrics separately (degree was not included, as the graphs were thresholded at the top 30% of edges for all participants, resulting in the same degree at the whole brain level). Wilks’ Lambda was used to test the extent to which the graph metrics contribute significantly to the discriminant function. A Chi-square test is used to evaluate the significance of the discriminant function, tested at p = .05.

### Association Between fPET and fMRI Graph Metrics and Cognition

To test the associations between the graph metrics and cognition, we first normalised the behavioural data so larger numbers indicate improved performance; this was applied to stop signal reaction time, seconds per correct response in the digit substitution and category switch reaction time. The cognitive scores were then converted to Z-scores.

To test the hypothesis that the graph metrics will differentially associate with cognitive performance across domains, we ran a series of canonical correlational analyses using the 10 cognitive measures. The graph theory metrics at the whole brain level from the fPET and fMRI were included in separate canonical correlation analysis to return a cognitive profile associated with the graph measures. The number of potential canonical variates in each analysis reflects the smallest number of variables in either set, in this case the three graph metrics. The F-statistic from the Wilk’s test was used to test the null hypothesis that the canonical correlation and all smaller ones are equal to zero.

To further test the hypothesis that the fPET and fMRI graph metrics will both predict cognitive performance, we used a series of multiple regression analyses. Whole brain global and local efficiency and betweenness centrality were used as the independent variables with each of the 10 cognitive measures as the dependent variable in separate analyses (degree was again excluded). A second series of regression analyses was run with age group also included to assess whether age mediated the relationships between the graph metrics and cognition.

## Results

### Sample characteristics

The characteristics of the younger (N = 40) and older (N = 46) cohorts of participants are shown in Table 1. The mean age of the younger group was 28 years and the older group 76 years. The younger group had a higher proportion of women (53%) and lower mean BMI (24.1 kg/m2) than those in the older group (46% women, BMI 25.7Kg/m2), but these differences were not statistically significant. Those in the older group had greater mean fasting glucose (5.2mmol/L) than the younger group (4.8mmol/L, p < .001).

**Table 1.** Mean and standard deviation (SD) of demographic and cognitive measures for older and younger adults, and p-value of statistical tests of age group differences^1^.

|  | Whole Sample<br>(N = 86) |  | Younger<br>(N = 40) |  | Older<br>(N = 46) |  | Younger<br>vs Older<br>p-value |
| --- | --- | --- | --- | --- | --- | --- | --- |
|  | Mean | SD | Mean | SD | Mean | SD |  |
| Age (years) | 53.5 | 24.8 | 27.9 | 6.2 | 75.8 | 6.0 | 0.000 |
| Sex (number and % female) | 42 (49%) |  | 21 (53%) |  | 21 (46%) |  | 0.662 |
| Education (years) | 17.0 | 3.5 | 18.0 | 2.7 | 16.3 | 4.0 | 0.028 |
| Body Mass Index (kg/m<sup>2</sup>) | 25.0 | 4.1 | 24.1 | 4.6 | 25.7 | 3.6 | 0.075 |
| Fasting blood glucose (mmol/L) | 5.0 | 0.5 | 4.8 | 0.4 | 5.2 | 0.6 | 0.000 |
| HVLT: Delayed recall (total) | 8.2 | 2.7 | 9.3 | 2.5 | 7.3 | 2.6 | 0.000 |
| HVLT: Recognition discrimination index | 9.8 | 2.0 | 10.7 | 1.6 | 9.1 | 2.1 | 0.000 |
| Digit span: Forward (longest) | 6.7 | 1.3 | 6.7 | 1.5 | 6.8 | 1.1 | 0.587 |
| Digit span: Backwards (longest) | 5.4 | 1.3 | 5.7 | 1.4 | 5.1 | 1.2 | 0.066 |
| Category switch: RT (seconds) | 0.38 | 0.37 | 0.28 | 0.23 | 0.48 | -0.43 | 0.011 |
| Category switch: % correct trials | 0.9 | 0.1 | 0.94 | 0.08 | 0.89 | 0.13 | 0.028 |
| Digit substitution: Correct count | 44.2 | 23.4 | 63.6 | 16.5 | 27.8 | 13.9 | 0.000 |
| Digit substitution: Se per correct | 5.9 | 14.4 | 3.40 | 9.31 | 8.01 | 17.45 | 0.144 |
| Stop signal: Prob of reacting | 48.9 | 11.2 | 51.0 | 12.6 | 47.2 | 9.7 | 0.115 |
| Stop signal: RT (seconds) | 27.5 | 0.1 | 0.27 | 0.05 | 0.28 | 0.06 | 0.170 |

HVLT = Hopkins Verbal Learning Test.
<sup>1</sup>All p-values are for *t*-tests (two-tailed) of age group differences, except for sex, which is tested with Chi-square (df = 1,86). An alpha of .05 was used to test for age group differences on each measure. *t*-test degrees of freedom were 84 for all measures except education and fasting blood glucose (81) and HVLT discrimination index (83).

### Cognitive function

Overall, the older adult group showed poorer memory, processing speed and cognitive flexibility compared to the younger adult group (Table 1). In particular, the older group showed lower delayed recall and recognition discrimination index (both p < .001) in the verbal learning test than did younger adults. Older adults also had a lower percentage of correct switch trials than younger adults (p = .028). Compared to younger adults, older adults had lower correct count in the digit substitution task (p < .001) and a slower reaction time in the category switch task (p = .011). Digit span and stop signal task performance was similar in both age groups.

### Similarity of fPET and fMRI Connectomes in the Whole Sample

Given that glucose metabolism contributes to the BOLD signal,^48^ we predicted that fPET and fMRI connectomes would have moderate similarity. While both whole sample averaged connectomes showed a clustering of connections in line with the functional network structure (Figure 1), a notable characteristic of the connectomes was the differences and heterogeneity of the strength of the connections between modalities. The within- and between-network connections using fPET were in the order of *r* = .20 to .35, and were relatively homogenous in strength (SD = .05) (Figure 1A). Although the connectivity strength was more heterogenous using fMRI (SD = .19) than fPET, these differences were not statistically significant (p = .515). The maximum correlations were larger in the fMRI (r = .88) than the fPET (r = .36) (p < .001).

**Figure 1.**
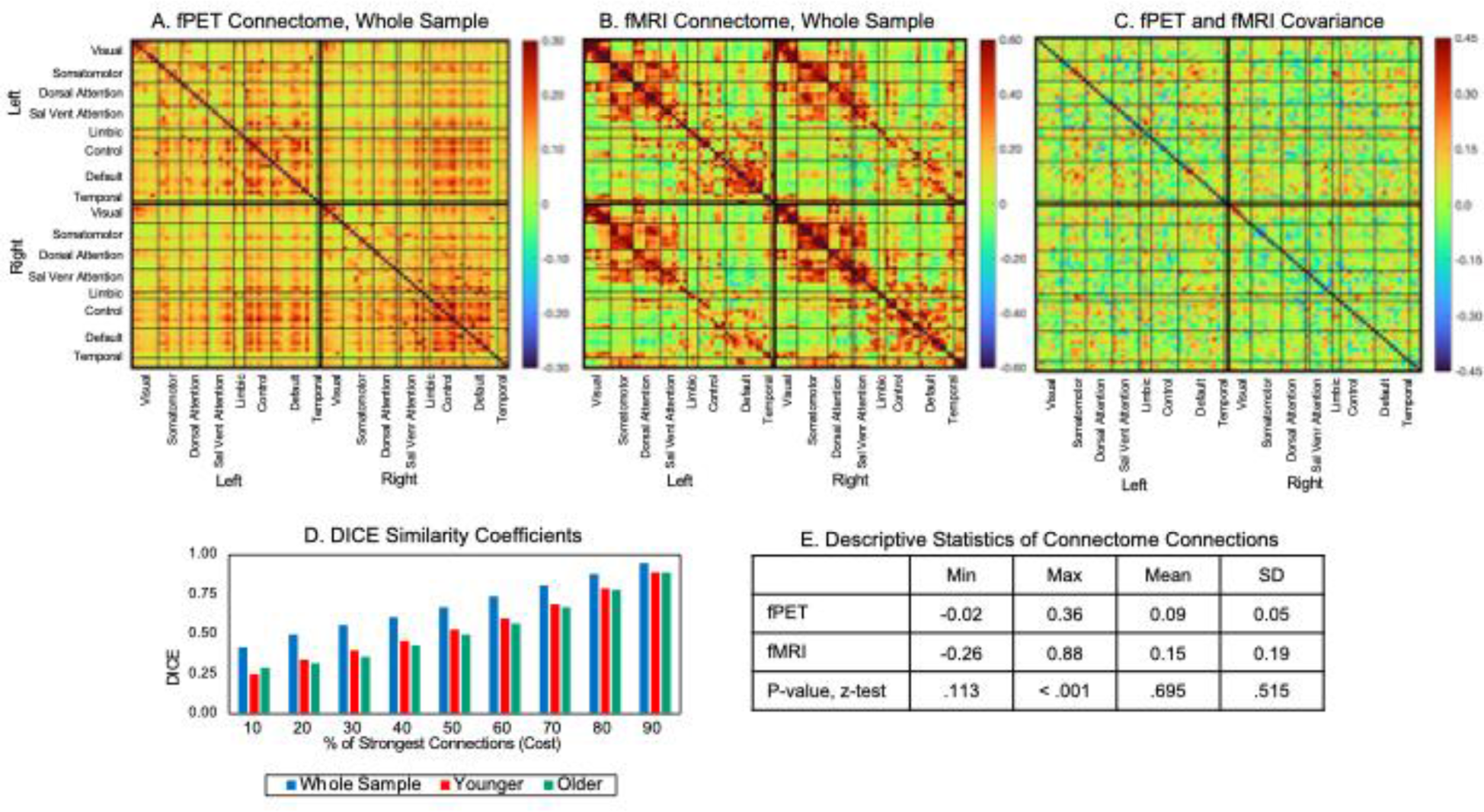
fPET and fMRI connectomes and covariance among the whole sample. (A) fPET connectome, (B) fMRI connectome,(C) across-subject covariance of fPET and fMRI connectomes, (D) DICE coefficients at connection costs, and (E) descriptive statistics and z-test of modality differences.

The DICE coefficient between the fPET and fMRI connectome was .67 for the top 50% of connections and .42 at top 10% of connections (Figure 1D). It is notable that the fPET and fMRI connectomes showed low-to-moderate covariance, with values ranging from r = -.36 to .40 (Figure 1C). However, the fPET-fMRI covariance patterns did not tend to cluster together: regions within- and between-networks differed in covariance between the two modalities.

### fPET and fMRI Connectome Characteristics and Age Group Differences

A salient qualitative characteristic of the group-averaged connectomes is the apparent age group differences in the fPET relative to the fMRI (Figure 2). Using fPET, younger adults had stronger connections than older adults (indicated by the darker red-to-orange colours), particularly between the dorsal attention, salience ventral attention, control and default networks. In contrast, using fMRI, the correlations were visually similar in both age groups with both younger and older adults showing relatively high connectivity between the somatomotor and dorsal attention and salience ventral attention networks, as well as the control and default networks.

**Figure 2.**
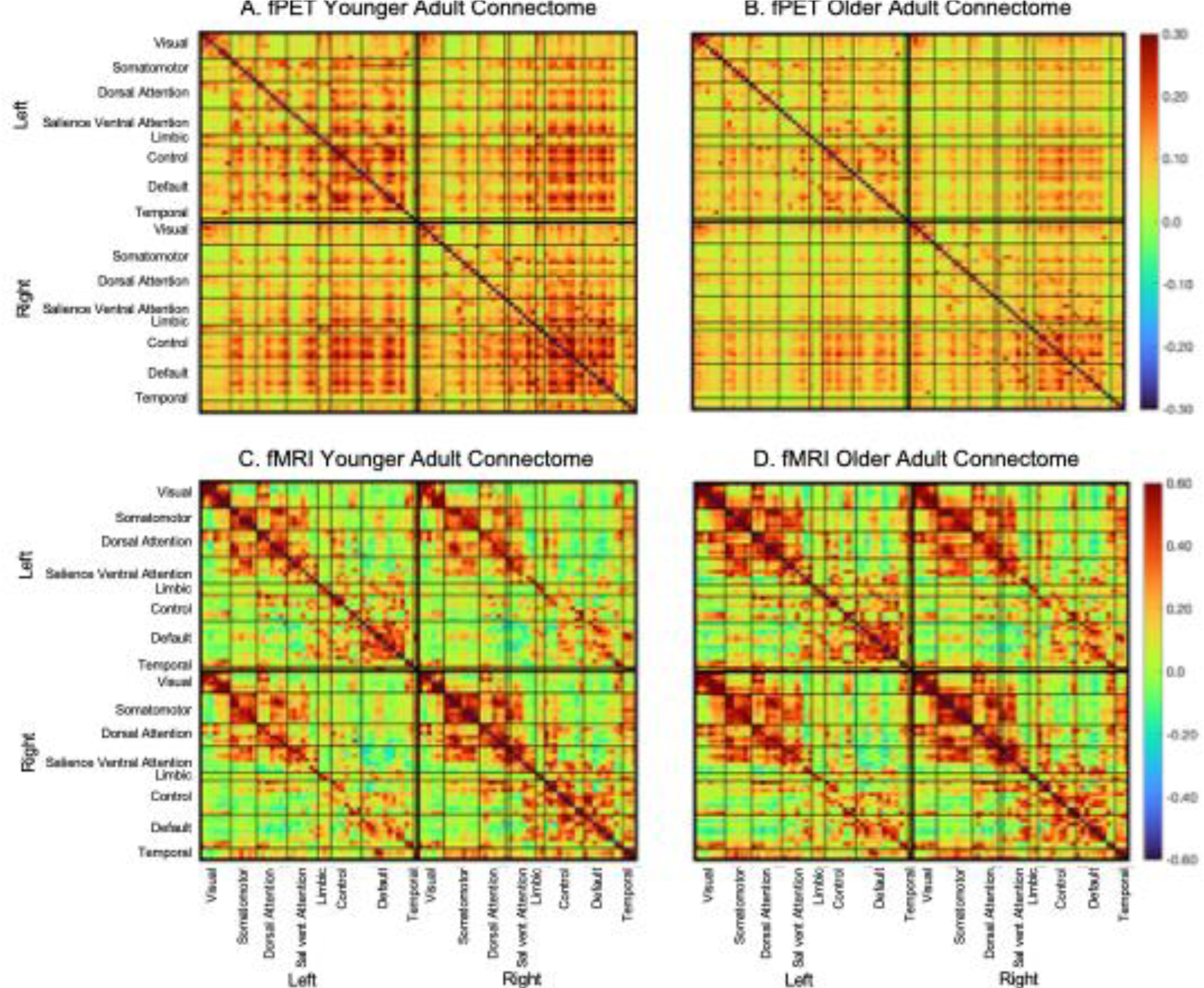
fPET and fMRI connectomes for younger and older adults using the Schaefer parcellation. (A) Younger and (B) older adult fPET metabolic connectomes, and (C) younger and (D) older adult fMRI connectomes. Connectomes are across the 100 nodes in 8 functional networks.

#### Within and Between-Network Connectivity

The strength, heterogeneity and topology of connections in the connectomes noted above for younger and older adults were evident when within- and between-network connectivity was calculated. There were also differences between fPET and fMRI in the connection strength for within-relative to between-network connectivity. The fPET connectomes had lower and more homogenous connection strength within-networks (Figures 3A and 3B) than the fMRI connectomes (Figure 3D and 3E). In contrast, the average fPET strength across all between-network connections was only slightly lower and showed less variability compared to the fMRI between-network connectivity (see Supplementary Table 1 for minimum, maximum, average and standard deviations of the connections).

**Figure 3.**
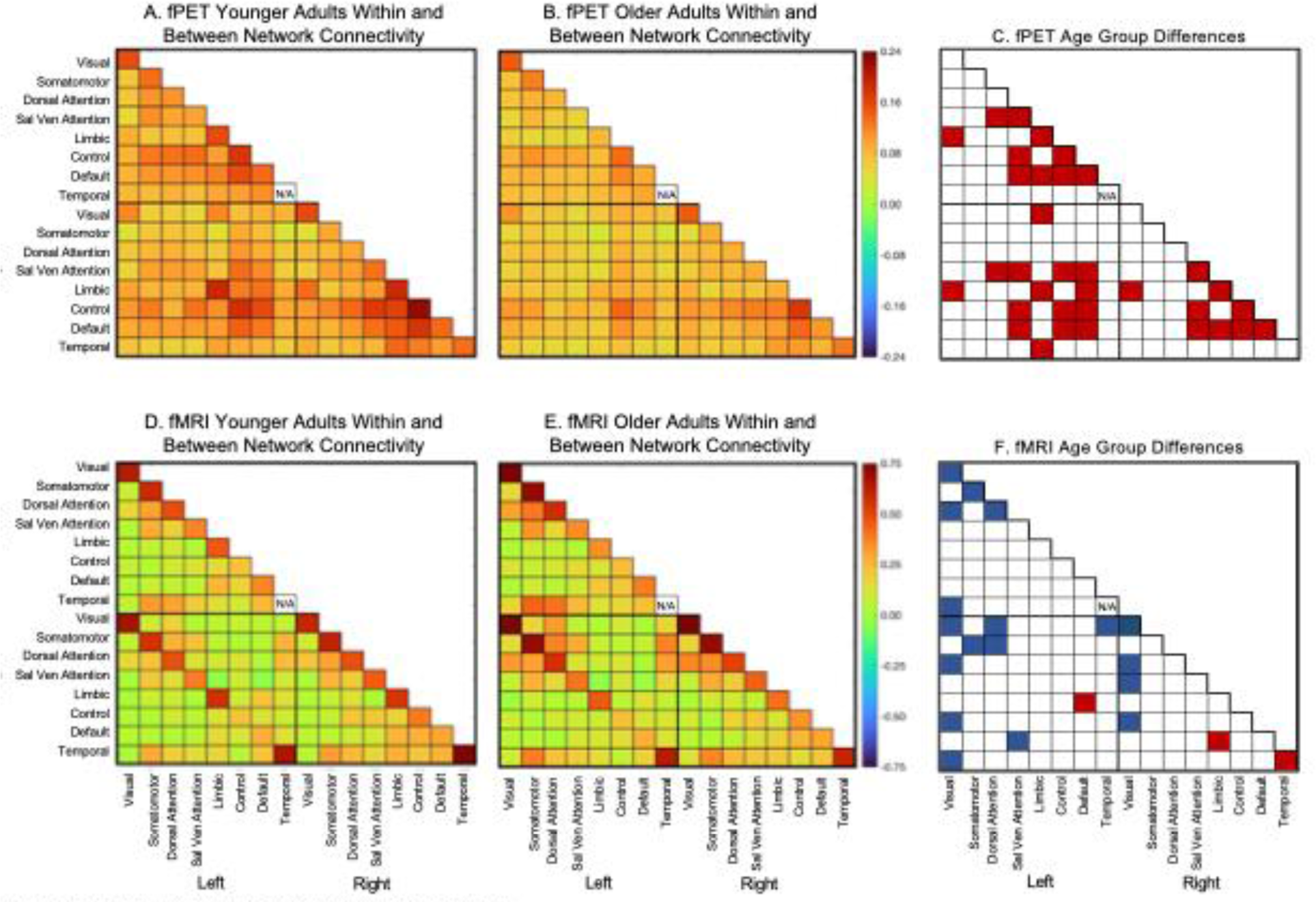
Within and between-network connectivity averaging the nodes within the networks for (A) younger and (B) older adults using fPET, and (D) Younger and (E) older adults in fMRI. Within connectivity is shown in the diagonal cells and between connectivity on the off-diagonal cells of each matrix. Significance test (*t*-test, df = 84) of younger vs older adults, with shaded cells indicating a statistically significant age group differences at p-FDR<.05 ((C) and (F)). Red shading younger > older; blue shading older > younger.

Although fPET showed a lower within-network connectivity maxima than fMRI, age group differences were more widespread in fPET than fMRI. Age-differences in within-network connectivity were found in eight fPET networks compared to five fMRI networks (Figure 3C & F). With fPET, younger adults had higher within-network connectivity strength than older adults bilaterally within the salience ventral attention, limbic, control and default networks. In contrast, with fMRI, the within-network connections were relatively high for both younger and older adults. Older adults also had higher fMRI within-network connections than younger adults in the left and right visual network and the left somatomotor and dorsal attention networks but low within network-connectivity in the right temporal network.

The topology and direction of age group differences in between-network connectivity were notably different for the fPET and fMRI. Using fPET, younger adults had higher between-network connectivity strength than older adults in 26 network pairs (Figure 3C). However, when using fMRI, younger adults had lower connectivity than older adults in most, but not all network pairs (16 of 19) (Figure 3F). For fPET, younger adults had higher network connectivity strength between the limbic, dorsal attention, salience ventral attention, control and default networks in the left and right hemispheres and across hemispheres. In the fMRI, young adults had higher connectivity strength between the left and right limbic and default networks.

In contrast, older adults had higher fMRI network connectivity between the visual and dorsal attention, temporal and control networks. Younger adults also had lower fMRI connectivity between the dorsal attention and somatomotor networks.

#### Regional Efficiency, Centrality and Degree

The strength, direction and topology of age group differences in the graph metrics was notably different between the fPET and fMRI. Significant age group differences were found in global efficiency for 20 fPET regions and 30 fMRI regions (Figure 4A and Supplementary Table 2). Younger adults had higher fPET global efficiency than older adults in regions in the prefrontal cortex in the control, salience ventral attention and default networks. In contrast, in the fMRI, young adults had higher global efficiency in the insula, temporal regions of the limbic network and left inferior parietal lobule and right parahippocampal cortex in the default network. Older adults had higher fPET and fMRI global efficiency than younger adults in a number of visual and somatomotor regions. For fMRI, older adults also had higher global efficiency in regions of the superior parietal lobule and postcentral cortex in the dorsal attention network and the precuneus in the control network.

**Figure 4.**
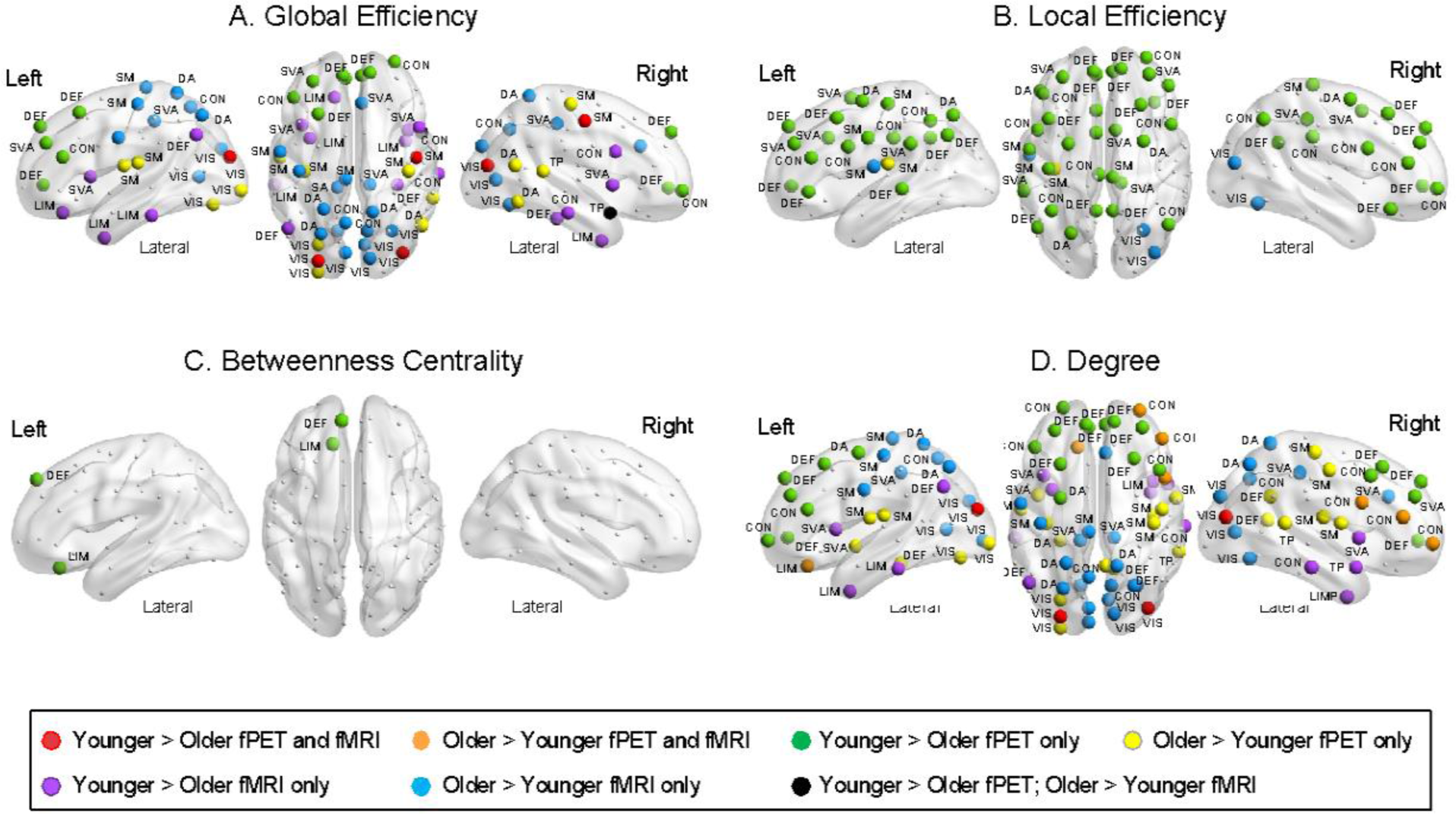
fPET and fMRI age group differences in regions of the Schaefer networks for (A) global efficiency, (B) local efficiency, (C) betweenness centrality, and (D) degree. Regions are shown as coloured dots where there is a statistically significant age group difference at p-FDR < .05 from one-sided *t*-tests, and the label indicates the network in which the region belongs. VIS = visual; SM = somatomotor; DA = dorsal attention, SVA = salience ventral attention; LIM = limbic; CON = control; DEF = default; TP = Temporal Parietal. For each region, the mean, standard deviations, age group effect sizes and t-tests are shown in Supplementary Tables 1 to 4.

The pattern of results was relatively similar for degree, with significant age group differences in 31 fPET regions and 32 fMRI regions (Figure 4D and supplementary Table 5). Younger adults had higher fPET degree than older adults in regions in the prefrontal cortex but lower degree in regions in the visual and sensorimotor network, and posterior regions of the control and default networks. For fMRI, younger adults had lower degree in regions in the visual and somatomotor networks but higher degree in temporal regions in the limbic and temporal parietal networks, the temporal poles and the insula in the salience ventral attention network.

Significant age group differences were found in local efficiency for 36 fPET regions and four fMRI regions (Figure 4B and supplementary Table 3). Younger adults had higher fPET local efficiency than older adults across the brain, including regions in the prefrontal, parietal and motor cortices and the cingulate in the control, dorsal attention, salience ventral attention, default and somatomotor networks. In contrast, older adults had higher fMRI local efficiency than younger adults in regions of the visual and somatomotor networks and lower local efficiency in the precuneus in the control network.

Significant age group differences were found in betweenness centrality for two fPET regions but no fMRI regions (Figure 4C and supplementary Table 4). Younger adults had higher fPET betweenness centrality than older adults in the orbital prefrontal cortex in the limbic network and dorsal prefrontal cortex in the default network.

#### Network Efficiency, Centrality and Degree

Age group differences at the network level largely followed the patterns at the region level. Older adults had significantly higher fPET and fMRI global efficiency than younger adults in the left and right visual network. Older adults also had higher fPET global efficiency than younger adults in the right somatomotor network and higher fMRI global efficiency in the left somatomotor network (Figure 5A). Older adults had higher fPET and fMRI betweenness centrality and degree than younger adults in the visual and somatomotor networks (Figure 5C and 5D). In contrast, younger adults had higher fPET local efficiency in the bilateral salience ventral attention, control and default networks (Figure 5B) and higher fMRI local efficiency in the left limbic and right dorsal attention networks. Younger adults also had higher fPET betweenness centrality in the left limbic, default, temporal parietal and right control networks (Figure 5C).

**Figure 5.**
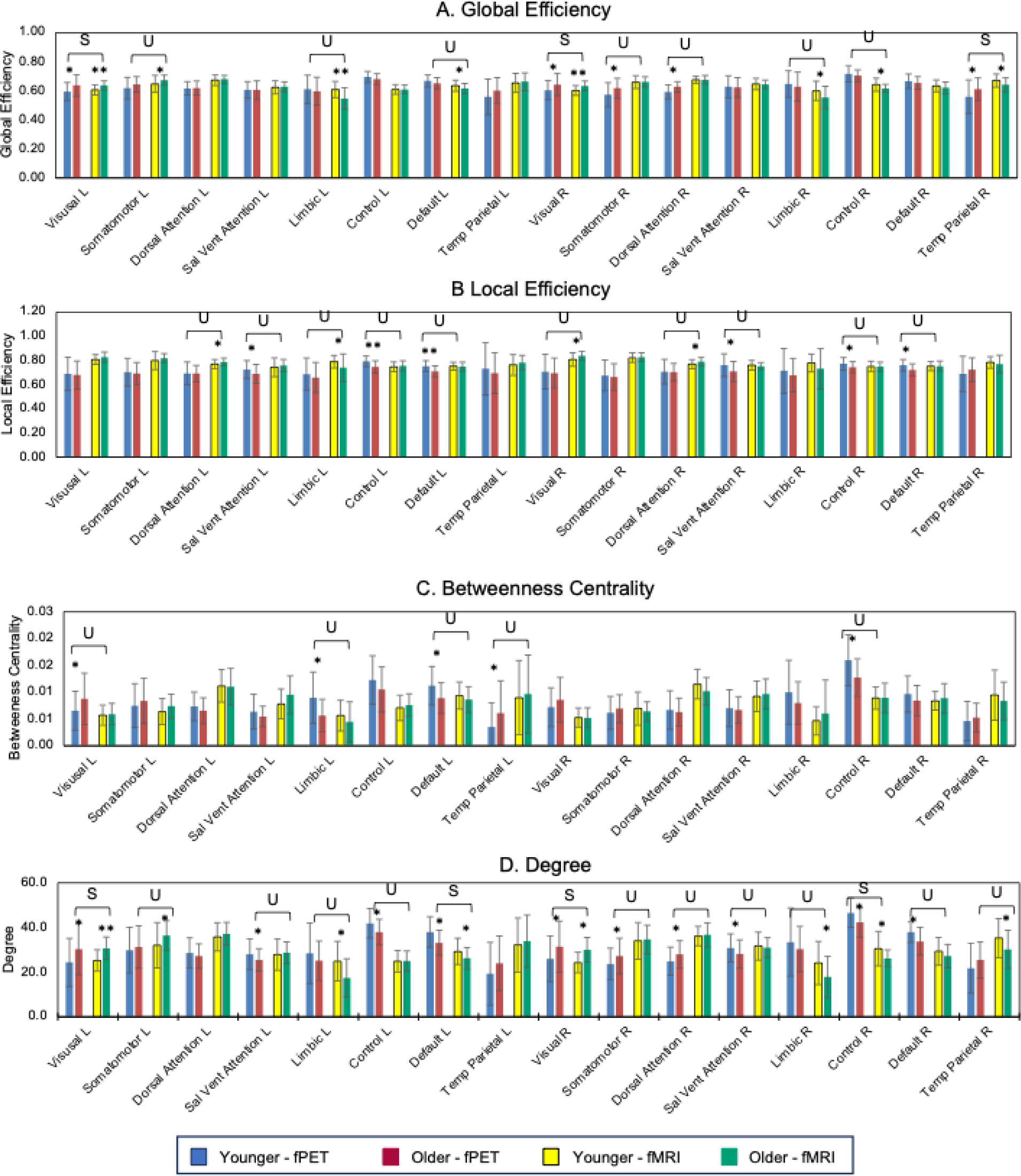
Mean and standard deviation (error bars) of fPET and fMRI graph metrics for younger and older adults in the Schaefer networks. Age group differences at *P-FDR < .05 and **p-FDR < .001 from one-sided *t*-tests. U = unique age group differences in one modality; S = Age group differences, same direction in both modalities.

### Relative Predictive Strength of fPET and fMRI Graph Metrics for Age Group

The discriminant function analysis was significant for the fPET whole brain graph metrics (Wilk’s Lambda = .795, Chi-square = 18.9, p < .001) but not the fMRI metrics (Wilk’s Lambda = .958, Chi-square = 3.5, p = .318). The probability of a correct age group classification based on the graph metrics was 60% and 61% for younger and older adults in the fPET, respectively; and 52% and 52% for younger and older adults in the fMRI.

### Multivariate Association Between fPET and fMRI and Cognition

The canonical correlation analyses revealed a closer multivariate relationship between the fPET whole brain graph metrics and the cognitive task performance than the fMRI whole brain graph metrics and cognition. The canonical correlation analysis of the fPET graph metrics identified one significant canonical mode (Figure 6). The linear combinations of the fPET graph metrics and the cognition measures were significantly correlated with each other, with a moderately high correlation coefficient of 0.60 (p = .041). However, for the fMRI analysis, the linear combinations of the graph metrics and the cognition measures were lower and not significant (r = .42, p = 1.1). For the fPET, higher local efficiency in particular was associated with better delayed recall and discrimination index score in the HVLT, a faster category switch reaction time and higher score in the digit substitution task.

**Figure 6.**
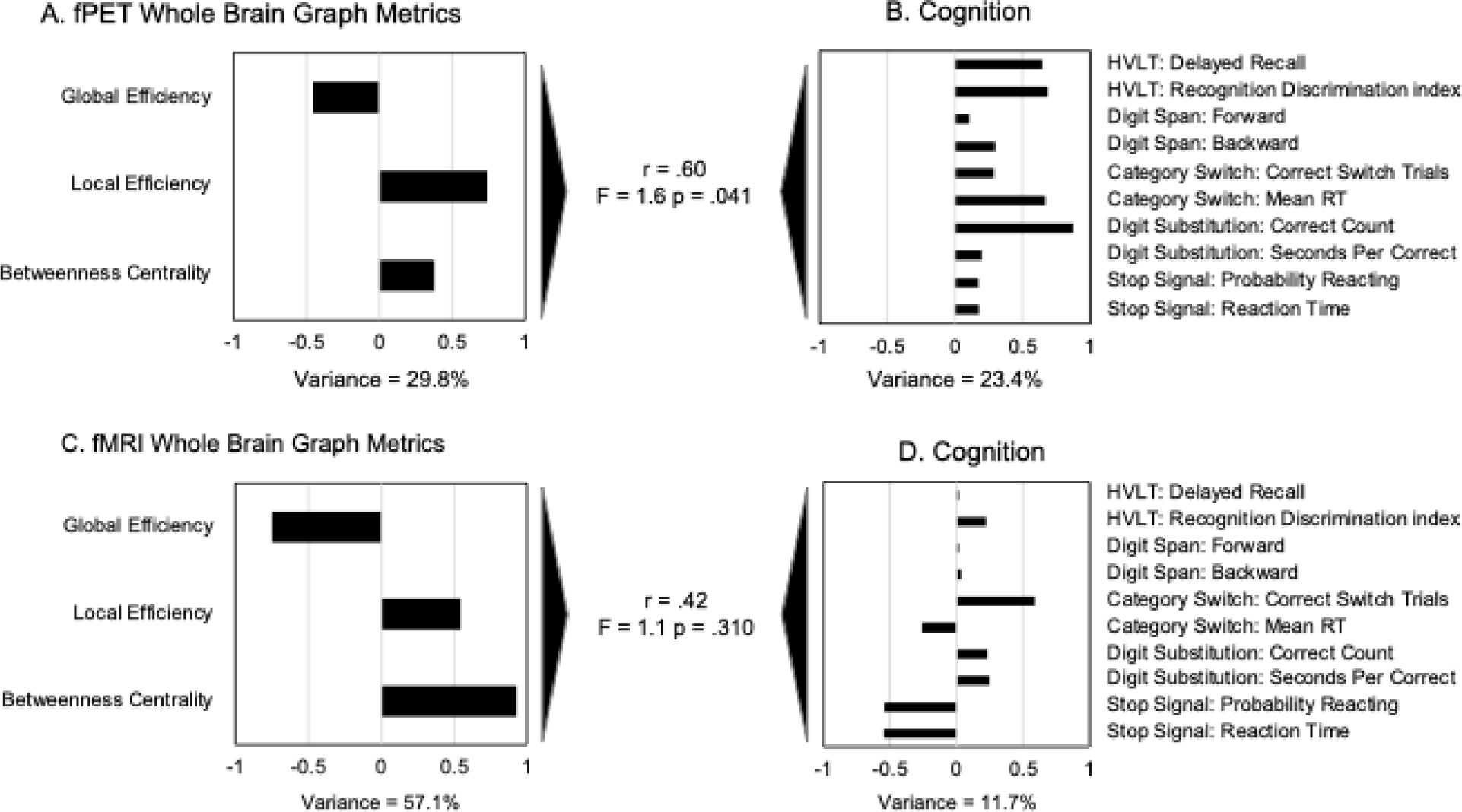
Canonical correlations between the whole brain graph metrics from fPET and fMRI ((A) and (C)) and cognition ((B) and (D)). r-value is the canonical correlation between the linear combinations of the graph metrics and cognition variables that maximally covary across subjects. F-statistic is Wilk’s test of the null hypothesis that the canonical correlation and all smaller ones are equal to zero and was significant for one canonical variate for fPET but not fMRI. The correlations on each variable set represent the strength of the association between the variable and the canonical variate. Variance explained is the percentage of variance explained by the variables in their variate. Stop signal reaction time, seconds per correct response in the digit substitution and category switch reaction time were multiplied by -1 so that higher scores reflect better performance. The test of difference between two correlations = 1.6; p = 0.114 (two tailed).

### Relative Predictive Strength of fPET and fMRI Graph Metrics for Cognition

The multiple regression analyses revealed that the fPET graph metrics were significantly associated with more cognitive measures than the fMRI graph metrics (Table 2). The regression analyses were significant for the discrimination index in the HVLT (p = .021), correct number of responses in the digit substitution task (p < .001), reaction time in the category switch task (p = .044), and probability of reaction in the stop signal task (p = .037). The regression analysis for the backward digit span approached significance (p = .051). Higher fPET local efficiency was associated with better recognition discrimination in the HVLT, more correct responses in the digit substitution task and faster reaction time in the category switch tasks. Higher fPET betweenness centrality was also associated with better performance in the backward digit span task and digit substitution task. Higher global efficiency and betweenness centrality in the fMRI was associated with lower probability of reacting in the stop signal task.

**Table 2.** Regression analyses predicting cognitive performance from fPET and fMRI whole brain graph metrics from the Schaefer Atlas. The ANOVA results for the overall models are shown in the top three rows, and the standardized beta weights for each graph metric’s relationship with cognition in subsequent rows. Significant beta weights are indicated at *p < .05 and **p < .001.

|  | HVLT:<br>Delayed<br>Recall | HVLT:<br>Discrim.<br>Index | Digit<br>Span:<br>Forward | Digit<br>Span:<br>Back | Category<br>Switch:<br>% Trials | Category<br>Switch:<br>RT | Dig Sub:<br>Number<br>Correct | Dig Sub:<br>Sec Per<br>Correct | Stop<br>Signal:<br>Pr<br>Reacting | Stop<br>Signal:<br>RT |
| --- | --- | --- | --- | --- | --- | --- | --- | --- | --- | --- |
| ANOVA F | 1.7 | 2.7 | 0.1 | 2.2 | 1.8 | 2.3 | 4.5 | 0.5 | 2.4 | 0.9 |
| ANOVA p | 0.132 | 0.021 | 0.995 | 0.051 | 0.109 | 0.044 | <.001 | 0.784 | 0.037 | 0.498 |
| Variance Explained | 12% | 17% | 1% | 14% | 12% | 15% | 26% | 4% | 15% | 6% |
| fPET: Global Efficiency | 0.08 | 0.25 | 0.04 | -0.05 | -0.15 | 0.20 | 0.11 | -0.17 | -0.03 | 0.04 |
| fPET: Local Efficiency | 0.40* | 0.53* | 0.03 | -0.01 | 0.09 | 0.47* | 0.55* | 0.02 | 0.14 | 0.13 |
| fPET: Betweenness Centrality | 0.15 | 0.20 | 0.07 | 0.33* | 0.15 | 0.15 | 0.23* | 0.12 | -0.12 | 0.00 |
| fMRI: Global Efficiency | 0.18 | -0.15 | -0.08 | 0.49 | 0.32 | 0.21 | 0.22 | 0.02 | -0.89* | -0.02 |
| fMRI: Local Efficiency | -0.09 | -0.11 | 0.05 | -0.09 | -0.13 | -0.12 | -0.26 | 0.07 | -0.15 | -0.01 |
| fMRI: Betweenness Centrality | 0.25 | 0.10 | -0.11 | 0.43 | 0.64 | 0.15 | 0.49 | 0.10 | -0.87* | -0.23 |

Discrim = discrimination; Dig Sub = Digit Substitution; Pr Reacting = probability of reacting.

When age group was included in the regression analyses (see Supplementary Table 6), a significant association remained between fPET local efficiency and the HVLT discrimination index and betweenness centrality in the backward digit span task (both p < .05). However, fPET local efficiency was no longer a significant predictor of the correct number of responses in the digit substitution task and category switch reaction time. Higher fMRI global efficiency and betweenness centrality remained significantly associated with probability of reacting in the stop signal task (p < .001).

## Discussion

In this simultaneous fPET/fMRI study, we found that whole-brain metabolic fPET connectomes have greater predictive utility for age and cognition than functional fMRI connectomes. Although fMRI connectomes showed higher maximum connectivity strength than fPET connectomes, fPET connectomes more strongly predicted age, revealed more age group differences in graph metrics and predicted cognition across more domains than fMRI connectomes. Together, these results build on our previous conclusion that fPET and fMRI provide complementary insights into brain function^24, 29^ by suggesting that metabolic connectivity has greater utility for understanding metabolic network changes in ageing and as a potential biomarker for functional and cognitive declines in ageing and neurodegeneration.^49^

As we hypothesised, the whole-brain fPET connectome was moderately similar in topology to the connectome calculated from fMRI data. These results were expected given that glucose metabolism contributes to the BOLD signal.^48^ We also found noteworthy differences between the modalities; some differences that supported and others that contrasted our hypotheses. Contrary to our hypothesis, age group differences in connectivity were more widespread in the fPET than the fMRI connectomes. As we expected, younger adults had higher fPET within-network connectivity in several associative networks, namely, bilaterally in the salience ventral attention, limbic, control and default networks. In fMRI, older adults also showed lower within-network connectivity, albeit in a smaller number of networks. Older adults had higher fMRI within-network connections than younger adults, mostly in primary processing and attentional networks. Older adults also had higher fMRI local efficiency in a number of regions in the dorsal attention, salience ventral attention and default networks. When between-network connectivity and global efficiency were assessed, the topology and direction of the age differences between the imaging modalities was also noteworthy. Younger adults had significantly higher fPET between-network connectivity and higher global efficiency than older adults, again primarily in the associative networks. A smaller number of associative networks showed age group differences in fMRI. Moreover, older adults had higher fMRI between-network connectivity than younger adults between the visual networks and the dorsal attention, control and default networks. Our fMRI results are consistent with previous research in terms of the primary processing and attentional networks. However, our results also diverge from previous fMRI research which has also largely shown higher connectivity among older adults between associative networks.^32^

A likely reason for the modality differences relates to the fact that fMRI and fPET are indexing different aspects of neuronal activity. The BOLD signal stems from a regional increase in cerebral blood flow, metabolic rate of oxygen, lactate, and a decrease in oxygen extraction fraction.^9^ In contrast, FDG-PET is less likely to be affected by blood flow or haemodynamic differences and provides a more direct measure of glucose uptake at the excitatory post-synaptic neuron.^50, 51^ As such, the differences we found between metabolic fPET and functional fMRI connectivity likely reflect at least in part different effects of ageing on the factors that contribute to the BOLD response and glucose metabolism. A second and related possible reason for the modality differences relates to the underlying rates of CMR_GLC_, cerebral blood flow and neurovascular coupling. Previous research has shown that blood flow and glucose metabolism can be uncoupled^35, 52^ or vary in their association across brain regions in ageing,^53, 54^ and even change in opposite directions from rests to task conditions.^55^ Similarly, there is regional variability between the BOLD signal and local field potentials in EEG,^56^ suggesting that neuronal activity and the BOLD signal are not always well coupled.

As we hypothesised, fPET and fMRI graph metrics were differentially associated with cognition across domains.^24^ However, we also found that differences in graph metrics predicted cognition across more domains in fPET than fMRI. Higher fPET whole-brain local efficiency was associated with better episodic and working memory, visuospatial performance, and speed of response inhibition. Higher global efficiency and betweenness centrality in the fMRI was associated with response inhibition. Similarly, the multivariate canonical relationship between the graph metrics and the cognitive domains was significant for fPET but not fMRI, although the level of the correlations was similar between modalities. The discriminant function analysis was also significant for fPET but not fMRI. Our results build on previous studies suggesting that fPET and fMRI are differentially associated with cognition^24, 29^ (also see^57, 58^). They indicate that there is a reconfiguration of the dynamic metabolic signals in ageing that is different to vascular and haemodynamic alterations and is more strongly linked to cognitive performance.

We assessed metabolic networks in the PET data at a temporal scale of 16 seconds. Additional advances in PET reconstruction and filtering techniques ^23, 59^ may further reduce the temporal resolution of fPET to that of fMRI. On the other hand, some evidence from animal models suggests that changes in metabolic flux emerge in less than half a second after neuronal activity^60^ but persist for tens of seconds.^55^ High-temporal resolution fPET studies in humans show that the FDG signal does not decrease to the baseline level immediately after the task stimulation.^22, 28^ For example, the FDG signal continued to increase during a 32 second working memory task with a return to baseline at approximately 10 seconds after task completion.^61^ Together these studies suggest that a timescale of around 10 seconds may set a limit on the ability of fPET to capture higher-frequency metabolic components of neural signals and their coherence across large-scale brain networks at rest. Additional research at different temporal scales is needed to further test this possibility.

A limitation of this study is the cross-sectional design. It is possible that the age differences we identified reflect underlying cohort or unmeasured differences between the two groups. It is also possible that alterations in metabolic connectivity are non-linear across the lifespan,^32^ something we cannot test due to the absence of middle age adults in our sample. Additional research, including longitudinal studies, is needed to assess fPET changes across the adult lifespan and to characterise the timeframe of their associations with normative cognitive ageing.

In conclusion, functional brain networks can be reliably derived from fPET data, reflecting the close link between neural activity and dynamic glucose metabolism across the brain network. fPET has greater predictive utility for age and cognition than fMRI. There is a reconfiguration of metabolic networks in ageing that is different to haemodynamic alterations and is more strongly linked to cognitive performance. These results highlight the important role that dynamic glucose metabolism plays in neuronal communication across the brain. They suggest that interventions to help maintain efficient brain metabolism in early and mid-adulthood may help to conserve metabolic connectivity and cognitive health in ageing. They also underscore the utility of metabolic connectivity to understand brain network changes in health and disease and as a potential biomarker for functional and cognitive declines in ageing and neurodegeneration.

## Data Availability Statement

The datasets used and/or analysed during the current study available from the corresponding author on reasonable request.

## Acknowledgements

We thank Rob Di Paolo, Gerard Murray, Navyaan Siddiqui, Richard McIntyre, Lauren Hudswell and the staff at Monash Biomedical Imaging for their contributions to data acquisition and image reconstruction.

## Participant Consent

Participants provided informed consent to participate in the study.

