## Supplementary Information for "Metabolic connectivity has greater predictive utility for age and cognition than functional connectivity"

### 1. Supplementary Methods

#### 1.1 Study Ethics

The study protocol was reviewed and approved by the Monash University Human Research Ethics Committee, in accordance with Australian Code for the Responsible Conduct of Research (2007) and the Australian National Statement on Ethical Conduct in Human Research (2007). Participants provided informed consent to participate in the study. Administration of ionizing radiation was approved by the Monash Health Principal Medical Physicist, following the Australian Radiation Protection and Nuclear Safety Agency Code of Practice (2005). For participants older than 18 years, the annual radiation exposure limit of 5 mSv applies; the effective dose in this study was 4.9 mSv.

#### 1.2 Cognitive Battery

Prior to the scan, participants completed an online demographic and lifestyle questionnaire including age, sex, education, height and weight, history of smoking, alcohol and recreational drug use. Participants also completed a cognitive test battery consisting of measures of general intelligence, working memory, cognitive flexibility, inhibitory control and verbal learning.

**Hopkins Verbal Learning Test (HVLT).** A three-trial list learning and free recall task comprising 12 words, four words from each of three semantic categories [1]. Approximately 20–25 minutes later, a delayed recall trial and a recognition trial was completed. The delayed recall required free recall of any words remembered. The recognition trial comprised 24 words, including the 12 target words and 12 false-positives, six semantically related, and six semantically unrelated. Delayed recall (total words recalled) and a recognition discrimination index (number of correct minus number of false positives in the recognition task) were calculated.

**Digit Span.** A measure of verbal short term and working memory used in two formats: Forward and backward digit span [2]. Participants were presented with a series of digits, and are asked to repeat them in either the order presented (forward span) or in reverse order (backwards span). After two consecutive failures of the same length, the test was stopped. Scores were derived as the length of the longest correct series for both forward and backward recall.

**Task Switching.** A computer-based test in which participants were given a word and had to perform one of two simple categorisation tasks, depending on the cue that appeared with the word: 1) 'living' task. If the cue was a heart, participants were asked to categorise the word via a key press based on whether it represents a LIVING versus a NON-LIVING object; and 2) 'size' task. If the cue was an arrow-cross, participants were asked to categorise the word via a key press based on whether it represents an object that is BIGGER or SMALLER than a basketball. The cue selection for each new trial was randomised. Half the test trials were switch trials; half non-switch trials. Half the switch and non-switch trials were congruent in the key presses for either task, half were incongruent. The measures used included the mean latency of correctly responding to a switch trial and switch cost. Switch cost is the difference between mean correct latency of switch trials and nonswitch trials with positive value indicating participants were slower on switch trials, that is, there was a latency cost to switching [3].

**Stop Signal.** A computer-based test in which participants were presented an arrow that pointed either right or left [4]. The task was to press the left response key if the arrow pointed to the left and press the right response key if the arrow pointed to the right, unless a signal beep is played after the presentation of the arrow. In this case the response should be stopped before execution. The delay between presentation of arrow and signal beep (starting at 250ms) was adjusted up or down (by 50ms) depending on performance. The delay got longer if the previous signal stop was successful (up to 1150ms) and smaller if the previous signal stop was not successful (down to 50ms). The stimulus onset asynchrony between the start of each trial (onset of fixation circles) was 2000ms. Variables were the mean reaction time in stop signal trials and stop signal reaction time. Stop signal reaction time is an estimate of inhibition ability, that is, the time required to stop the initiated go-process. The slower the stop signal reaction time, the more difficult to stop the go-process.

**Digit Symbol Substitution.** A computer-based task in which participant were presented with an 18 columns x 16 rows matrix [5]. The task was to translate symbols shown above the matrix (key) into digits in the matrix within a two minute period. Total count of correct responses and seconds per correct response were recorded.

#### 1.3 MR-PET Data Acquisition

Participants underwent a 90-minute simultaneous MR-PET scan in a Siemens (Erlangen) Biograph 3-Tesla molecular MR scanner. Participants were directed to consume a high-protein/low-sugar diet for the 24 hours prior to the scan. They were also instructed to fast for six hours and to drink 2–6 glasses of water. Prior to FDG infusion, participants were cannulated in the vein in each forearm and a 10ml baseline blood sample taken. At the beginning of the scan, half of the 260 MBq FDG tracer was administered via the left forearm as a bolus, providing a strong PET signal from the beginning of the scan. The remaining 130 MBq of the FDG tracer dose was infused at a rate of 36ml/hour over 50 minutes, minimising the amount of signal decay over the course of the data acquisition. We have previously demonstrated that this protocol provides a good balance between a fast increase in signal-to-noise ratio at the start of the scan, and maintenance of signal-to-noise ratio over the duration of the scan [6].

Participants were positioned supine in the scanner bore with their head in a 32-channel radiofrequency head coil and were instructed to lie as still as possible. The scan sequence was as follows. Non-functional MRI scans were acquired during the first 12 minutes, including a T1 3DMPRAGE (TA = 3.49 min, TR = 1640ms, TE = 234ms, flip angle = 8°, field of view = 256 × 256 mm<sup>2</sup>, voxel size = 1.0 × 1.0 × 1.0 mm<sup>3</sup>, 176 slices, sagittal acquisition) and T2 FLAIR (TA = 5.52 min, TR = 5,000ms, TE = 396ms, field of view = 250 × 250 mm<sup>2</sup>, voxel size = .5 × .5 × 1 mm<sup>3</sup>, 160 slices) to image the anatomical grey and white matter structures, respectively. Thirteen minutes into the scan, list-mode PET (voxel size = 1.39 × 1.39 × 5.0mm<sup>3</sup>) and T2\* EPI BOLD-fMRI (TA = 40 minutes; TR = 1000ms, TE = 39ms, FOV = 210 mm<sup>2</sup>, 2.4 × 2.4 × 2.4 mm<sup>3</sup> voxels, 64 slices, ascending axial acquisition) sequences were initiated. A 40-minute resting-state scan was undertaken in naturalistic viewing conditions watching a movie of a drone flying over the Hawaii Islands. At 53 minutes, pseudo-continuous arterial spin labelling (pc-ASL) began, and at 58 minutes, diffusion-weighted imaging (DWI) was acquired with 71 directions to index white matter connectivity. pcASL, DWI and fMRI results are not reported here.

Plasma radioactivity levels were measured throughout the duration of the scan. Beginning at 10-minutes post infusion onset, 5ml blood samples were taken from the right forearm using a vacutainer at 10-minute intervals for a total of nine samples. The blood sample were immediately placed in a Heraeus Megafuge 16 centrifuge (ThermoFisher Scientific, Osterode, Germany) and spun at 2,000 rpm (RCF ~ 515g) for 5 minutes. 1,000-μL plasma was pipetted, transferred to a counting tube, and placed in a well counter for four minutes. The count start time, total number of counts, and counts per minute were recorded for each sample.

##### 1.4 Correction for Partial Volume Effects

PET images were corrected for partial volume effects using the modified Müller-Gartner method implemented in PetSurf (https://surfer.nmr.mgh.harvard.edu/fswiki/PetSurfer) [7, 8]. The method corrects for white matter spill in and grey matter spill out of the PET signal. The equation subtracts from the grey matter voxel signal the white matter signal (convoluted by the point spread function) and divides by the grey matter signal. This division can introduce over-correction at the grey matter boundary, and hence a grey matter binary mask is recommended, with the threshold level needing to be chosen. A grey matter threshold of 20-30% is recommended in ageing because atrophy can influence results [7]. For our analyses, we chose a 25% grey matter threshold and surface-based spatial smoothing [8]. We used a Gaussian kernel with a full width at half maximum of 8mm to increase the signal-to-noise ratio. Subcortical structures were partial volume corrected and spatially smoothed in volume space and merged with the cortical data.

##### 1.5 Graph Theory Metrics

The following graph theory metrics were calculated:

**Global Efficiency** at each node defined as the average of the shortest inverse-distances between the node and all other nodes in the graph. Across the entire graph, global efficiency represents a measure of global integration.

**Local Efficiency** at each node defined as the average of shortest inverse-distances between the nodes within the neighbouring sub-graph (all nodes neighbouring that node and all existing edges among them). Network local efficiency is a measure of local integration of a network.

**Betweenness Centrality** defined as the proportion of times that a node is part of a shortest-path between any two nodes within a graph. It represents a measure of node centrality within a graph.

**Degree** at each node defined as the number of edges from and to that node. Degree characterises the local connectedness of each region within the network.

### 2. Supplementary Data from the Main Manuscript

Supplementary Table 1. Descriptive statistics and z-tests of differences in within- and between-network connectivity in the connectomes of younger and older adults in fPET and fMRI. The connectomes are shown in Figure 2 of the main manuscript.

|  | Within Network Connectivity |  |  |  | Between Network Connectivity |  |  |  |
| --- | --- | --- | --- | --- | --- | --- | --- | --- |
| fPET |  |  |  |  |  |  |  |  |
|  | Min | Max | Mean | SD | Min | Max | Mean | SD |
| Younger | 0.09 | 0.22 | 0.14 | 0.04 | 0.04 | 0.22 | 0.10 | 0.03 |
| Older | 0.06 | 0.17 | 0.11 | 0.03 | 0.04 | 0.17 | 0.08 | 0.02 |
| fMRI |  |  |  |  |  |  |  |  |
|  | Min | Max | Mean | SD | Min | Max | Mean | SD |
| Younger | 0.23 | 0.74 | 0.48 | 0.14 | -0.03 | 0.67 | 0.15 | 0.13 |
| Older | 0.24 | 0.78 | 0.49 | 0.18 | -0.03 | 0.83 | 0.18 | 0.14 |
| fPET vs fMRI, p-value (Z-test) |  |  |  |  |  |  |  |  |
|  | Min | Max | Mean | SD | Min | Max | Mean | SD |
| Younger | .534 | .002 | .100 | .664 | .763 | .012 | .827 | .665 |
| Older | .392 | .000 | .048 | .481 | .745 | .000 | .637 | .575 |

Supplementary Table 2. Mean, standard deviation and effect sizes (Cohen's D) of regional global efficiency for younger and older adults in fPET and fMRI from the Schaefer Atlas, and t-tests of age group differences.

|  | fPET |  |  |  |  |  |  |  | fMRI |  |  |  |  |  |  |  |
| --- | --- | --- | --- | --- | --- | --- | --- | --- | --- | --- | --- | --- | --- | --- | --- | --- |
|  | Younger |  | Older |  | Younger vs Older |  |  |  | Younger |  | Older |  | Younger vs Older |  |  |  |
|  | Mean | SD | Mean | SD | Cohen's D | t-value | p |  | Mean | SD | Mean | SD | Cohen's D | t-value | p |  |
| Visual Central: Extra Striate Cortex 1 L | 0.60 | 0.10 | 0.66 | 0.10 | -0.55 | -2.6 | 0.006 | 0.037 | 0.61 | 0.07 | 0.63 | 0.05 | -0.21 | -1.0 | 0.167 | 0.232 |
| Visual Central: Extra Striate Cortex 2 L | 0.63 | 0.10 | 0.69 | 0.10 | -0.64 | -3.0 | 0.002 | 0.020 | 0.58 | 0.08 | 0.60 | 0.07 | -0.32 | -1.5 | 0.073 | 0.133 |
| Visual Central: Striate Cortex 1 L | 0.59 | 0.11 | 0.64 | 0.09 | -0.48 | -2.2 | 0.014 | 0.061 | 0.57 | 0.07 | 0.60 | 0.06 | -0.46 | -2.1 | 0.018 | 0.057 |
| Visual Central: Extra Striate Cortex 3 L | 0.59 | 0.09 | 0.65 | 0.08 | -0.71 | -3.3 | 0.001 | 0.015 | 0.61 | 0.05 | 0.66 | 0.05 | -0.93 | -4.3 | 0.000 | 0.001 |
| Visual Peripheral: Extra Striate Inferior 1 L | 0.54 | 0.09 | 0.57 | 0.12 | -0.28 | -1.3 | 0.098 | 0.177 | 0.63 | 0.07 | 0.66 | 0.06 | -0.36 | -1.7 | 0.049 | 0.107 |
| Visual Peripheral: Striate Cortex Calcarine 1 L | 0.60 | 0.09 | 0.58 | 0.12 | 0.10 | 0.5 | 0.316 | 0.369 | 0.61 | 0.07 | 0.66 | 0.06 | -0.69 | -3.2 | 0.001 | 0.007 |
| Visual Peripheral: Extra Striate Cortex Sup 1 L | 0.58 | 0.10 | 0.59 | 0.08 | -0.09 | -0.4 | 0.339 | 0.383 | 0.63 | 0.07 | 0.67 | 0.05 | -0.75 | -3.5 | 0.000 | 0.005 |
| Somatomotor A: 1 L | 0.62 | 0.14 | 0.64 | 0.09 | -0.17 | -0.8 | 0.221 | 0.283 | 0.65 | 0.08 | 0.69 | 0.05 | -0.55 | -2.6 | 0.006 | 0.026 |
| Somatomotor A: 2 L | 0.64 | 0.15 | 0.65 | 0.09 | -0.12 | -0.6 | 0.285 | 0.347 | 0.65 | 0.07 | 0.70 | 0.04 | -0.80 | -3.7 | 0.000 | 0.004 |
| Somatomotor B: Auditory 1 L | 0.69 | 0.08 | 0.69 | 0.08 | -0.05 | -0.2 | 0.411 | 0.446 | 0.66 | 0.08 | 0.67 | 0.07 | -0.17 | -0.8 | 0.217 | 0.274 |
| Somatomotor B: S2 1 L | 0.47 | 0.17 | 0.57 | 0.11 | -0.70 | -3.3 | 0.001 | 0.015 | 0.67 | 0.07 | 0.66 | 0.08 | 0.25 | 1.2 | 0.122 | 0.183 |
| Somatomotor B: S2 2 L | 0.58 | 0.09 | 0.63 | 0.08 | -0.61 | -2.8 | 0.003 | 0.024 | 0.68 | 0.07 | 0.66 | 0.05 | 0.29 | 1.3 | 0.095 | 0.161 |
| Somatomotor B: Central 1 L | 0.59 | 0.09 | 0.61 | 0.09 | -0.29 | -1.4 | 0.088 | 0.177 | 0.58 | 0.09 | 0.63 | 0.07 | -0.62 | -2.9 | 0.003 | 0.012 |
| Dorsal Attention A: Temporal Occipital 1 L | 0.60 | 0.11 | 0.63 | 0.09 | -0.33 | -1.5 | 0.065 | 0.144 | 0.69 | 0.06 | 0.67 | 0.05 | 0.32 | 1.5 | 0.070 | 0.133 |
| Dorsal Attention A: Parietal Occipital 1 L | 0.55 | 0.10 | 0.59 | 0.11 | -0.41 | -1.9 | 0.032 | 0.085 | 0.69 | 0.06 | 0.68 | 0.08 | 0.26 | 1.2 | 0.112 | 0.179 |
| Dorsal Attention A: Superior Parietal Lobule 1 L | 0.67 | 0.07 | 0.67 | 0.07 | 0.00 | 0.0 | 0.493 | 0.494 | 0.66 | 0.05 | 0.68 | 0.05 | -0.50 | -2.3 | 0.012 | 0.043 |
| Dorsal Attention B: Post Central 1 L | 0.60 | 0.09 | 0.58 | 0.12 | 0.25 | 1.2 | 0.125 | 0.199 | 0.67 | 0.07 | 0.69 | 0.04 | -0.31 | -1.4 | 0.077 | 0.138 |
| Dorsal Attention B: Post Central 2 L | 0.53 | 0.16 | 0.58 | 0.11 | -0.36 | -1.7 | 0.051 | 0.116 | 0.65 | 0.07 | 0.67 | 0.05 | -0.34 | -1.6 | 0.061 | 0.120 |
| Dorsal Attention B: Post Central 3 L | 0.62 | 0.10 | 0.61 | 0.08 | 0.11 | 0.5 | 0.302 | 0.360 | 0.67 | 0.05 | 0.70 | 0.04 | -0.63 | -2.9 | 0.002 | 0.012 |
| Dorsal Attention B: Frontal Eye Fields 1 L | 0.66 | 0.11 | 0.61 | 0.10 | 0.48 | 2.2 | 0.015 | 0.061 | 0.67 | 0.08 | 0.68 | 0.06 | -0.20 | -0.9 | 0.183 | 0.244 |
| Salience Ventral Attention A: Parietal Operculum 1 L | 0.64 | 0.08 | 0.62 | 0.08 | 0.27 | 1.3 | 0.107 | 0.181 | 0.67 | 0.05 | 0.65 | 0.07 | 0.36 | 1.7 | 0.049 | 0.107 |
| Salience Ventral Attention A: Insula: 1 L | 0.43 | 0.18 | 0.51 | 0.15 | -0.45 | -2.1 | 0.020 | 0.069 | 0.63 | 0.08 | 0.59 | 0.12 | 0.38 | 1.8 | 0.040 | 0.098 |
| Salience Ventral Attention A: Insula: 2 L | 0.55 | 0.16 | 0.60 | 0.08 | -0.42 | -1.9 | 0.028 | 0.081 | 0.64 | 0.06 | 0.62 | 0.06 | 0.49 | 2.2 | 0.014 | 0.045 |
| Salience Ventral Attention A: Parietal Medial 1 L | 0.57 | 0.12 | 0.60 | 0.11 | -0.24 | -1.1 | 0.136 | 0.200 | 0.67 | 0.08 | 0.71 | 0.06 | -0.64 | -2.9 | 0.002 | 0.012 |
| Salience Ventral Attention A: Frontal Medial 1 L | 0.61 | 0.11 | 0.61 | 0.12 | 0.03 | 0.1 | 0.445 | 0.459 | 0.62 | 0.08 | 0.64 | 0.06 | -0.20 | -0.9 | 0.182 | 0.244 |
| Salience Ventral Attention B: Lateral Prefrontal Cortex 1 L | 0.73 | 0.07 | 0.69 | 0.06 | 0.60 | 2.8 | 0.003 | 0.025 | 0.57 | 0.08 | 0.61 | 0.08 | -0.42 | -1.9 | 0.029 | 0.079 |
| Salience Ventral Attention B: Medial Posterior Prefrontal 1 L | 0.63 | 0.08 | 0.59 | 0.12 | 0.36 | 1.7 | 0.049 | 0.114 | 0.62 | 0.08 | 0.63 | 0.07 | -0.25 | -1.2 | 0.124 | 0.183 |
| Limbic A Temporal Pole 1 L | 0.67 | 0.12 | 0.67 | 0.08 | 0.04 | 0.2 | 0.425 | 0.449 | 0.59 | 0.08 | 0.50 | 0.14 | 0.84 | 3.9 | 0.000 | 0.003 |
| Limbic A: Temporal Pole 2 L | 0.53 | 0.15 | 0.53 | 0.16 | 0.00 | 0.0 | 0.494 | 0.494 | 0.61 | 0.08 | 0.57 | 0.08 | 0.51 | 2.3 | 0.011 | 0.042 |
| Limbic B: Orbital Frontal Cortex 1 L | 0.66 | 0.10 | 0.61 | 0.13 | 0.41 | 1.9 | 0.031 | 0.085 | 0.63 | 0.07 | 0.58 | 0.07 | 0.74 | 3.4 | 0.000 | 0.005 |
| Control A: Intraparietal Sulcus 1 L | 0.69 | 0.07 | 0.67 | 0.06 | 0.25 | 1.2 | 0.121 | 0.198 | 0.62 | 0.05 | 0.62 | 0.06 | -0.01 | 0.0 | 0.489 | 0.494 |
| Control A: Lateral Prefrontal Cortex 1 L | 0.76 | 0.05 | 0.73 | 0.07 | 0.58 | 2.7 | 0.004 | 0.031 | 0.59 | 0.06 | 0.60 | 0.05 | -0.08 | -0.4 | 0.362 | 0.393 |
| Control A: Lateral Prefrontal Cortex 2 L | 0.64 | 0.08 | 0.62 | 0.07 | 0.25 | 1.2 | 0.123 | 0.198 | 0.64 | 0.06 | 0.61 | 0.06 | 0.46 | 2.1 | 0.017 | 0.056 |
| Control B: Lateral Prefrontal Cortex 1 L | 0.75 | 0.07 | 0.72 | 0.06 | 0.44 | 2.0 | 0.022 | 0.069 | 0.62 | 0.09 | 0.59 | 0.08 | 0.26 | 1.2 | 0.120 | 0.183 |
| Control C: Precuneus 1 L | 0.66 | 0.07 | 0.67 | 0.08 | -0.18 | -0.8 | 0.209 | 0.274 | 0.58 | 0.07 | 0.60 | 0.06 | -0.37 | -1.7 | 0.046 | 0.105 |
| Control C: Precuneus 2 L | 0.61 | 0.08 | 0.62 | 0.12 | -0.13 | -0.6 | 0.271 | 0.335 | 0.61 | 0.07 | 0.66 | 0.06 | -0.85 | -3.9 | 0.000 | 0.003 |
| Control C: Cingulate Posterior 1 L | 0.60 | 0.13 | 0.63 | 0.12 | -0.30 | -1.4 | 0.084 | 0.172 | 0.58 | 0.07 | 0.60 | 0.06 | -0.17 | -0.8 | 0.212 | 0.271 |
| Default A: Dorsal Prefrontal Cortex 1 L | 0.68 | 0.10 | 0.62 | 0.08 | 0.70 | 3.2 | 0.001 | 0.015 | 0.60 | 0.08 | 0.60 | 0.09 | -0.03 | -0.2 | 0.439 | 0.458 |
| Default A: Precuneus Posterior Cingulate Cortex 1 L | 0.69 | 0.09 | 0.71 | 0.12 | -0.19 | -0.9 | 0.196 | 0.265 | 0.62 | 0.06 | 0.62 | 0.06 | 0.13 | 0.6 | 0.269 | 0.332 |
| Default A: Medial Prefrontal Cortex 1 L | 0.67 | 0.07 | 0.62 | 0.07 | 0.61 | 2.8 | 0.003 | 0.024 | 0.63 | 0.07 | 0.61 | 0.08 | 0.27 | 1.2 | 0.111 | 0.179 |
| Default B: Temp 1 L | 0.57 | 0.16 | 0.60 | 0.08 | -0.21 | -1.0 | 0.168 | 0.237 | 0.65 | 0.07 | 0.64 | 0.06 | 0.27 | 1.3 | 0.105 | 0.175 |
| Default B: Temp 2 L | 0.60 | 0.09 | 0.62 | 0.08 | -0.17 | -0.8 | 0.211 | 0.274 | 0.65 | 0.08 | 0.62 | 0.06 | 0.34 | 1.6 | 0.060 | 0.120 |
| Default B: Inferior Parietal Lobule 1 L | 0.71 | 0.07 | 0.69 | 0.07 | 0.28 | 1.3 | 0.103 | 0.177 | 0.66 | 0.06 | 0.62 | 0.06 | 0.69 | 3.2 | 0.001 | 0.007 |
| Default B: Dorsal Prefrontal Cortex 1 L | 0.75 | 0.05 | 0.68 | 0.08 | 1.03 | 4.8 | 0.000 | 0.000 | 0.65 | 0.07 | 0.64 | 0.07 | 0.10 | 0.5 | 0.327 | 0.372 |
| Default B: Lateral Prefrontal Cortex 1 L | 0.68 | 0.08 | 0.66 | 0.07 | 0.28 | 1.3 | 0.103 | 0.177 | 0.62 | 0.08 | 0.62 | 0.06 | 0.00 | 0.0 | 0.494 | 0.494 |
| Default B: Ventral Prefrontal Cortex 1 L | 0.58 | 0.12 | 0.59 | 0.12 | -0.04 | -0.2 | 0.419 | 0.449 | 0.62 | 0.06 | 0.59 | 0.11 | 0.31 | 1.4 | 0.080 | 0.140 |
| Default B: Ventral Prefrontal Cortex 2 L | 0.75 | 0.08 | 0.73 | 0.08 | 0.28 | 1.3 | 0.103 | 0.177 | 0.63 | 0.06 | 0.61 | 0.06 | 0.40 | 1.9 | 0.033 | 0.085 |
| Default C: RetroSuperior Parietal Lobule 1 L | 0.54 | 0.11 | 0.60 | 0.12 | -0.44 | -2.0 | 0.023 | 0.071 | 0.61 | 0.05 | 0.61 | 0.05 | -0.03 | -0.1 | 0.446 | 0.460 |
| Default C: Parahippocampal Cortex 1 L | 0.45 | 0.14 | 0.51 | 0.15 | -0.40 | -1.9 | 0.034 | 0.085 | 0.60 | 0.07 | 0.57 | 0.11 | 0.23 | 1.1 | 0.144 | 0.203 |
| Temporal Parietal 1 L | 0.56 | 0.12 | 0.60 | 0.09 | -0.42 | -1.9 | 0.028 | 0.081 | 0.66 | 0.07 | 0.66 | 0.06 | -0.13 | -0.6 | 0.282 | 0.343 |
| Visual Central: Extra Striate Cortex 1 R | 0.61 | 0.10 | 0.63 | 0.09 | -0.27 | -1.2 | 0.111 | 0.185 | 0.61 | 0.05 | 0.64 | 0.05 | -0.64 | -3.0 | 0.002 | 0.011 |
| Visual Central: Extra Striate Cortex 2 R | 0.66 | 0.10 | 0.71 | 0.11 | -0.47 | -2.2 | 0.017 | 0.063 | 0.57 | 0.07 | 0.59 | 0.06 | -0.24 | -1.1 | 0.135 | 0.193 |
| Visual Central: Extra Striate Cortex 3 R | 0.59 | 0.09 | 0.67 | 0.09 | -0.84 | -3.9 | 0.000 | 0.005 | 0.61 | 0.05 | 0.64 | 0.05 | -0.62 | -2.9 | 0.002 | 0.012 |
| Visual Peripheral: Striate Cortex Calcarine 1 R | 0.60 | 0.12 | 0.62 | 0.09 | -0.23 | -1.1 | 0.142 | 0.203 | 0.58 | 0.06 | 0.63 | 0.06 | -0.77 | -3.5 | 0.000 | 0.005 |
| Visual Peripheral: Extra Striate Inferior 1 R | 0.54 | 0.11 | 0.57 | 0.13 | -0.24 | -1.1 | 0.135 | 0.200 | 0.66 | 0.08 | 0.68 | 0.06 | -0.41 | -1.9 | 0.031 | 0.081 |
| Visual Peripheral: Extra Striate Superior 1 R | 0.58 | 0.10 | 0.60 | 0.08 | -0.28 | -1.3 | 0.097 | 0.177 | 0.62 | 0.05 | 0.66 | 0.06 | -0.72 | -3.3 | 0.001 | 0.006 |
| Somatomotor A: 1 R | 0.49 | 0.15 | 0.57 | 0.09 | -0.67 | -3.1 | 0.001 | 0.018 | 0.66 | 0.06 | 0.69 | 0.04 | -0.49 | -2.3 | 0.012 | 0.043 |
| Somatomotor A: 2 R | 0.52 | 0.16 | 0.60 | 0.09 | -0.66 | -3.0 | 0.002 | 0.020 | 0.65 | 0.07 | 0.66 | 0.05 | -0.11 | -0.5 | 0.304 | 0.362 |
| Somatomotor A: 3 R | 0.51 | 0.17 | 0.56 | 0.11 | -0.37 | -1.7 | 0.044 | 0.108 | 0.67 | 0.06 | 0.69 | 0.04 | -0.25 | -1.2 | 0.123 | 0.183 |
| Somatomotor A: 4 R | 0.58 | 0.17 | 0.61 | 0.13 | -0.20 | -0.9 | 0.174 | 0.241 | 0.67 | 0.05 | 0.68 | 0.04 | -0.32 | -1.5 | 0.072 | 0.133 |
| Somatomotor B: Auditory 1 R | 0.69 | 0.09 | 0.71 | 0.07 | -0.20 | -0.9 | 0.184 | 0.252 | 0.68 | 0.07 | 0.66 | 0.07 | 0.26 | 1.2 | 0.113 | 0.179 |
| Somatomotor B: S2 1 R | 0.47 | 0.17 | 0.54 | 0.14 | -0.46 | -2.1 | 0.019 | 0.067 | 0.66 | 0.06 | 0.67 | 0.10 | 0.4 | 0.328 | 0.372 |  |
| Somatomotor B: S2 2 R | 0.56 | 0.11 | 0.61 | 0.08 | -0.50 | -2.3 | 0.011 | 0.054 | 0.67 | 0.05 | 0.66 | 0.06 | -0.11 | 0.5 | 0.313 | 0.368 |
| Somatomotor B: Central 1 R | 0.55 | 0.13 | 0.60 | 0.07 | -0.47 | -2.2 | 0.017 | 0.063 | 0.60 | 0.09 | 0.60 | 0.08 | -0.08 | -0.4 | 0.358 | 0.393 |
| Dorsal Attention A: Temporal Occipital 1 R | 0.58 | 0.09 | 0.62 | 0.08 | -0.55 | -2.6 | 0.006 | 0.037 | 0.68 | 0.05 | 0.66 | 0.06 | 0.35 | 1.6 | 0.057 | 0.120 |
| Dorsal Attention A: Parietal Occipital 1 R | 0.61 | 0.08 | 0.65 | 0.07 | -0.54 | -2.5 | 0.007 | 0.037 | 0.69 | 0.06 | 0.67 | 0.06 | 0.41 | 1.9 | 0.029 | 0.079 |
| Dorsal Attention A: Superior Parietal Lobule 1 R | 0.61 | 0.13 | 0.65 | 0.07 | -0.47 | -2.2 | 0.016 | 0.063 | 0.64 | 0.06 | 0.66 | 0.05 | -0.38 | -1.8 | 0.042 | 0.100 |
| Dorsal Attention B: Post Central 1 R | 0.52 | 0.15 | 0.56 | 0.12 | -0.32 | -1.5 | 0.073 | 0.156 | 0.68 | 0.05 | 0.67 | 0.06 | 0.20 | 0.9 | 0.181 | 0.244 |
| Dorsal Attention B: Post Central 2 R | 0.58 | 0.13 | 0.61 | 0.07 | -0.31 | -1.4 | 0.081 | 0.168 | 0.67 | 0.05 | 0.71 | 0.05 | -0.77 | -3.5 | 0.000 | 0.005 |
| Dorsal Attention B: Frontal Eye Fields 1 R | 0.63 | 0.14 | 0.62 | 0.09 | 0.10 | 0.5 | 0.321 | 0.369 | 0.68 | 0.05 | 0.69 | 0.06 | -0.04 | -0.2 | 0.432 | 0.458 |
| Salience Ventral Attention A: Parietal Operculum 1 R | 0.60 | 0.10 | 0.61 | 0.07 | -0.12 | -0.6 | 0.288 | 0.347 |  |  |  |  |  |  |  |  |

Supplementary Table 3. Mean, standard deviation and effect sizes (Cohen's D) of regional local efficiency for younger and older adults in fPET and fMRI from the Schaefer Atlas, and t-tests of age group differences.

|  | fPET |  |  |  |  |  |  |  |  | fMRI |  |  |  |  |  |  |  |  |
| --- | --- | --- | --- | --- | --- | --- | --- | --- | --- | --- | --- | --- | --- | --- | --- | --- | --- | --- |
|  | Younger |  | Older |  | Younger vs Older |  |  |  | p-FDR | Younger |  | Older |  | Younger vs Older |  |  |  | p-FDR |
|  | Mean | SD | Mean | SD | Cohen's D | t-value | p | p |  | Mean | SD | Mean | SD | Cohen's D | t-value | p | p |  |
| Visual Central: Extra Striate Cortex 1 L | 0.71 | 0.22 | 0.69 | 0.17 | 0.11 | 0.5 | 0.312 | 0.389 | 0.81 | 0.06 | 0.84 | 0.06 | -0.44 | -2.0 | 0.022 | 0.150 |  |  |
| Visual Central: Extra Striate Cortex 2 L | 0.74 | 0.15 | 0.71 | 0.12 | 0.19 | 0.9 | 0.190 | 0.283 | 0.81 | 0.09 | 0.85 | 0.08 | -0.46 | -2.1 | 0.019 | 0.150 |  |  |
| Visual Central: Striate Cortex 1 L | 0.69 | 0.29 | 0.70 | 0.13 | -0.08 | -0.4 | 0.360 | 0.428 | 0.81 | 0.10 | 0.82 | 0.10 | -0.17 | -0.8 | 0.211 | 0.360 |  |  |
| Visual Central: Extra Striate Cortex 3 L | 0.70 | 0.21 | 0.70 | 0.14 | 0.00 | 0.0 | 0.497 | 0.497 | 0.80 | 0.05 | 0.82 | 0.05 | -0.31 | -1.4 | 0.078 | 0.280 |  |  |
| Visual Peripheral: Extra Striate Inferior 1 L | 0.63 | 0.30 | 0.66 | 0.19 | -0.12 | -0.5 | 0.297 | 0.376 | 0.82 | 0.04 | 0.83 | 0.05 | -0.26 | -1.2 | 0.119 | 0.280 |  |  |
| Visual Peripheral: Striate Cortex Calcarine 1 L | 0.68 | 0.25 | 0.63 | 0.23 | 0.22 | 1.0 | 0.152 | 0.242 | 0.81 | 0.05 | 0.81 | 0.05 | 0.03 | 0.2 | 0.436 | 0.474 |  |  |
| Visual Peripheral: Extra Striate Cortex Sup 1 L | 0.67 | 0.28 | 0.66 | 0.19 | 0.02 | 0.1 | 0.456 | 0.480 | 0.80 | 0.05 | 0.81 | 0.04 | -0.24 | -1.1 | 0.134 | 0.280 |  |  |
| Somatomotor A: 1 L | 0.77 | 0.17 | 0.68 | 0.15 | 0.53 | 2.4 | 0.008 | 0.031 | 0.79 | 0.15 | 0.82 | 0.05 | -0.31 | -1.4 | 0.076 | 0.280 |  |  |
| Somatomotor A: 2 L | 0.72 | 0.20 | 0.69 | 0.15 | 0.17 | 0.8 | 0.221 | 0.315 | 0.82 | 0.08 | 0.82 | 0.05 | 0.08 | 0.4 | 0.362 | 0.426 |  |  |
| Somatomotor B: Auditory 1 L | 0.75 | 0.17 | 0.74 | 0.13 | 0.06 | 0.3 | 0.388 | 0.446 | 0.77 | 0.11 | 0.79 | 0.06 | -0.25 | -1.2 | 0.125 | 0.280 |  |  |
| Somatomotor B: S2 1 L | 0.57 | 0.34 | 0.72 | 0.11 | -0.63 | -2.8 | 0.004 | 0.015 | 0.79 | 0.05 | 0.80 | 0.06 | -0.30 | -1.4 | 0.085 | 0.280 |  |  |
| Somatomotor B: S2 2 L | 0.68 | 0.19 | 0.69 | 0.19 | -0.03 | -0.1 | 0.451 | 0.480 | 0.78 | 0.06 | 0.82 | 0.06 | -0.74 | -3.4 | 0.000 | 0.023 |  |  |
| Somatomotor B: Central 1 L | 0.79 | 0.14 | 0.66 | 0.20 | 0.69 | 3.2 | 0.001 | 0.006 | 0.83 | 0.14 | 0.86 | 0.05 | -0.31 | -1.4 | 0.077 | 0.280 |  |  |
| Dorsal Attention A: Temporal Occipital 1 L | 0.62 | 0.27 | 0.71 | 0.12 | -0.44 | -2.0 | 0.023 | 0.059 | 0.76 | 0.04 | 0.78 | 0.05 | -0.43 | -2.0 | 0.025 | 0.156 |  |  |
| Dorsal Attention A: Parietal Occipital 1 L | 0.62 | 0.29 | 0.69 | 0.18 | -0.26 | -1.2 | 0.122 | 0.207 | 0.79 | 0.06 | 0.79 | 0.06 | -0.09 | -0.4 | 0.335 | 0.420 |  |  |
| Dorsal Attention A: Superior Parietal Lobule 1 L | 0.80 | 0.09 | 0.75 | 0.07 | 0.70 | 3.2 | 0.001 | 0.005 | 0.77 | 0.05 | 0.78 | 0.05 | -0.28 | -1.3 | 0.103 | 0.280 |  |  |
| Dorsal Attention B: Post Central 1 L | 0.76 | 0.21 | 0.67 | 0.21 | 0.41 | 1.9 | 0.032 | 0.070 | 0.77 | 0.14 | 0.81 | 0.05 | -0.47 | -2.2 | 0.016 | 0.150 |  |  |
| Dorsal Attention B: Post Central 2 L | 0.71 | 0.24 | 0.68 | 0.15 | 0.14 | 0.6 | 0.270 | 0.353 | 0.81 | 0.07 | 0.83 | 0.07 | -0.24 | -1.1 | 0.134 | 0.280 |  |  |
| Dorsal Attention B: Post Central 3 L | 0.72 | 0.25 | 0.70 | 0.16 | 0.11 | 0.5 | 0.315 | 0.389 | 0.80 | 0.07 | 0.80 | 0.06 | -0.02 | -0.1 | 0.458 | 0.486 |  |  |
| Dorsal Attention B: Frontal Eye Fields 1 L | 0.79 | 0.11 | 0.66 | 0.21 | 0.76 | 3.5 | 0.000 | 0.004 | 0.75 | 0.07 | 0.77 | 0.05 | -0.31 | -1.4 | 0.078 | 0.280 |  |  |
| Saliency Ventral Attention A: Parietal Operculum 1 L | 0.80 | 0.12 | 0.73 | 0.12 | 0.59 | 2.7 | 0.004 | 0.016 | 0.77 | 0.09 | 0.76 | 0.13 | 0.13 | 0.6 | 0.273 | 0.386 |  |  |
| Saliency Ventral Attention A: Insula: 1 L | 0.56 | 0.26 | 0.63 | 0.19 | -0.29 | -1.2 | 0.114 | 0.197 | 0.79 | 0.10 | 0.80 | 0.10 | -0.17 | -0.8 | 0.218 | 0.360 |  |  |
| Saliency Ventral Attention A: Insula: 2 L | 0.67 | 0.25 | 0.71 | 0.14 | -0.22 | -1.0 | 0.162 | 0.253 | 0.77 | 0.07 | 0.77 | 0.09 | -0.08 | -0.4 | 0.351 | 0.426 |  |  |
| Saliency Ventral Attention A: Parietal Medial 1 L | 0.79 | 0.14 | 0.74 | 0.11 | 0.42 | 1.9 | 0.032 | 0.070 | 0.74 | 0.07 | 0.77 | 0.05 | -0.52 | -2.4 | 0.009 | 0.097 |  |  |
| Saliency Ventral Attention A: Frontal Medial 1 L | 0.78 | 0.18 | 0.70 | 0.17 | 0.47 | 2.1 | 0.018 | 0.050 | 0.78 | 0.08 | 0.78 | 0.07 | 0.01 | 0.1 | 0.473 | 0.488 |  |  |
| Saliency Ventral Attention B: Lateral Prefrontal Cortex 1 L | 0.80 | 0.05 | 0.76 | 0.06 | 0.79 | 3.7 | 0.000 | 0.002 | 0.70 | 0.18 | 0.74 | 0.11 | -0.25 | -1.1 | 0.130 | 0.280 |  |  |
| Saliency Ventral Attention B: Medial Prefrontal Cortex 1 L | 0.81 | 0.11 | 0.70 | 0.17 | 0.83 | 3.8 | 0.000 | 0.002 | 0.71 | 0.15 | 0.74 | 0.06 | -0.30 | -1.4 | 0.082 | 0.280 |  |  |
| Limbic A: Temporal Pole 1 L | 0.74 | 0.10 | 0.74 | 0.12 | 0.04 | 0.2 | 0.434 | 0.480 | 0.78 | 0.10 | 0.77 | 0.16 | 0.12 | 0.6 | 0.289 | 0.391 |  |  |
| Limbic A: Temporal Pole 2 L | 0.70 | 0.18 | 0.62 | 0.20 | 0.41 | 1.8 | 0.038 | 0.077 | 0.80 | 0.08 | 0.75 | 0.19 | 0.35 | 1.6 | 0.053 | 0.250 |  |  |
| Limbic B: Orbital Frontal Cortex 1 L | 0.75 | 0.13 | 0.71 | 0.14 | 0.30 | 1.4 | 0.089 | 0.161 | 0.78 | 0.07 | 0.76 | 0.09 | 0.27 | 1.3 | 0.107 | 0.280 |  |  |
| Control A: Intraparietal Sulcus 1 L | 0.81 | 0.05 | 0.75 | 0.07 | 1.03 | 4.8 | 0.000 | 0.000 | 0.74 | 0.06 | 0.75 | 0.06 | -0.15 | -0.7 | 0.250 | 0.373 |  |  |
| Control A: Lateral Prefrontal Cortex 1 L | 0.79 | 0.04 | 0.75 | 0.06 | 0.71 | 3.3 | 0.001 | 0.005 | 0.73 | 0.09 | 0.75 | 0.07 | -0.16 | -0.7 | 0.235 | 0.362 |  |  |
| Control A: Lateral Prefrontal Cortex 2 L | 0.81 | 0.12 | 0.73 | 0.12 | 0.67 | 3.1 | 0.001 | 0.006 | 0.75 | 0.06 | 0.73 | 0.10 | 0.21 | 1.0 | 0.172 | 0.320 |  |  |
| Control B: Lateral Prefrontal Cortex 1 L | 0.78 | 0.06 | 0.76 | 0.06 | 0.27 | 1.2 | 0.107 | 0.189 | 0.77 | 0.08 | 0.77 | 0.11 | 0.00 | 0.0 | 0.499 | 0.499 |  |  |
| Control C: Precuneus 1 L | 0.80 | 0.14 | 0.75 | 0.13 | 0.34 | 1.6 | 0.057 | 0.111 | 0.75 | 0.10 | 0.78 | 0.06 | -0.41 | -1.9 | 0.031 | 0.173 |  |  |
| Control C: Precuneus 2 L | 0.78 | 0.16 | 0.74 | 0.09 | 0.30 | 1.4 | 0.088 | 0.161 | 0.72 | 0.10 | 0.76 | 0.06 | -0.52 | -2.4 | 0.009 | 0.097 |  |  |
| Control C: Cingulate Posterior 1 L | 0.83 | 0.07 | 0.75 | 0.12 | 0.71 | 3.2 | 0.001 | 0.005 | 0.78 | 0.07 | 0.76 | 0.07 | 0.27 | 1.3 | 0.106 | 0.280 |  |  |
| Default A: Dorsal Prefrontal Cortex 1 L | 0.80 | 0.14 | 0.74 | 0.15 | 0.41 | 1.9 | 0.032 | 0.070 | 0.71 | 0.14 | 0.73 | 0.14 | -0.18 | -0.8 | 0.208 | 0.360 |  |  |
| Default A: Precuneus Posterior Cingulate Cortex 1 L | 0.81 | 0.06 | 0.76 | 0.06 | 0.83 | 3.8 | 0.000 | 0.002 | 0.74 | 0.06 | 0.74 | 0.08 | 0.09 | 0.4 | 0.336 | 0.420 |  |  |
| Default A: Medial Prefrontal Cortex 1 L | 0.81 | 0.09 | 0.70 | 0.18 | 0.80 | 3.7 | 0.000 | 0.002 | 0.74 | 0.11 | 0.76 | 0.09 | -0.13 | -0.6 | 0.274 | 0.386 |  |  |
| Default B: Temp 1 L | 0.68 | 0.23 | 0.68 | 0.19 | 0.00 | 0.0 | 0.492 | 0.497 | 0.77 | 0.06 | 0.75 | 0.06 | 0.20 | 0.9 | 0.179 | 0.320 |  |  |
| Default B: Temp 2 L | 0.78 | 0.11 | 0.73 | 0.11 | 0.50 | 2.3 | 0.012 | 0.040 | 0.78 | 0.06 | 0.76 | 0.08 | 0.23 | 1.1 | 0.143 | 0.285 |  |  |
| Default B: Inferior Parietal Lobule 1 L | 0.79 | 0.06 | 0.73 | 0.10 | 0.63 | 2.9 | 0.002 | 0.010 | 0.75 | 0.04 | 0.74 | 0.04 | 0.36 | 1.7 | 0.049 | 0.247 |  |  |
| Default B: Dorsal Prefrontal Cortex 1 L | 0.78 | 0.05 | 0.73 | 0.09 | 0.69 | 3.2 | 0.001 | 0.006 | 0.76 | 0.06 | 0.74 | 0.09 | 0.24 | 1.1 | 0.133 | 0.280 |  |  |
| Default B: Lateral Prefrontal Cortex 1 L | 0.81 | 0.12 | 0.74 | 0.10 | 0.63 | 2.9 | 0.002 | 0.010 | 0.74 | 0.11 | 0.74 | 0.10 | -0.04 | -0.2 | 0.431 | 0.474 |  |  |
| Default B: Ventral Prefrontal Cortex 2 L | 0.79 | 0.10 | 0.75 | 0.13 | 0.41 | 1.9 | 0.032 | 0.070 | 0.76 | 0.08 | 0.75 | 0.07 | 0.16 | 0.8 | 0.225 | 0.360 |  |  |
| Default B: Ventral Prefrontal Cortex 2 L | 0.78 | 0.07 | 0.74 | 0.06 | 0.52 | 2.4 | 0.009 | 0.032 | 0.75 | 0.06 | 0.75 | 0.05 | -0.16 | -0.7 | 0.228 | 0.360 |  |  |
| Default C: RetroSuperior Parietal Lobule/central 1 L | 0.67 | 0.30 | 0.70 | 0.13 | -0.15 | -0.7 | 0.249 | 0.341 | 0.75 | 0.07 | 0.78 | 0.11 | -0.28 | -1.3 | 0.097 | 0.280 |  |  |
| Default C: Parahippocampal Cortex 1 L | 0.60 | 0.31 | 0.52 | 0.24 | 0.29 | 1.2 | 0.108 | 0.189 | 0.77 | 0.16 | 0.82 | 0.09 | -0.39 | -1.8 | 0.040 | 0.209 |  |  |
| Temporal Parietal 1 L | 0.73 | 0.22 | 0.70 | 0.17 | 0.18 | 0.8 | 0.203 | 0.298 | 0.76 | 0.09 | 0.78 | 0.06 | -0.27 | -1.2 | 0.110 | 0.280 |  |  |
| Visual Central: Extra Striate Cortex 1 R | 0.73 | 0.16 | 0.73 | 0.08 | 0.02 | 0.1 | 0.456 | 0.480 | 0.79 | 0.05 | 0.83 | 0.04 | -0.82 | -3.8 | 0.000 | 0.013 |  |  |
| Visual Central: Extra Striate Cortex 2 R | 0.75 | 0.11 | 0.70 | 0.13 | 0.40 | 1.9 | 0.033 | 0.070 | 0.82 | 0.11 | 0.86 | 0.07 | -0.42 | -1.9 | 0.029 | 0.171 |  |  |
| Visual Central: Extra Striate Cortex 3 R | 0.76 | 0.17 | 0.72 | 0.17 | 0.23 | 1.0 | 0.149 | 0.242 | 0.79 | 0.07 | 0.84 | 0.05 | -0.71 | -3.3 | 0.001 | 0.023 |  |  |
| Visual Peripheral: Striate Cortex Calcarine 1 R | 0.70 | 0.22 | 0.70 | 0.14 | 0.04 | 0.2 | 0.431 | 0.480 | 0.83 | 0.08 | 0.85 | 0.05 | -0.30 | -1.4 | 0.087 | 0.280 |  |  |
| Visual Peripheral: Extra Striate Inferior 1 R | 0.68 | 0.23 | 0.68 | 0.14 | -0.03 | -0.1 | 0.453 | 0.480 | 0.79 | 0.04 | 0.81 | 0.05 | -0.27 | -1.3 | 0.105 | 0.280 |  |  |
| Visual Peripheral: Extra Striate Superior 1 R | 0.69 | 0.23 | 0.69 | 0.18 | 0.02 | 0.1 | 0.454 | 0.480 | 0.81 | 0.06 | 0.83 | 0.05 | -0.26 | -1.2 | 0.115 | 0.280 |  |  |
| Somatomotor A: 1 R | 0.62 | 0.30 | 0.64 | 0.21 | -0.07 | -0.3 | 0.375 | 0.436 | 0.81 | 0.10 | 0.81 | 0.06 | -0.05 | -0.2 | 0.414 | 0.470 |  |  |
| Somatomotor A: 2 R | 0.64 | 0.26 | 0.63 | 0.23 | 0.01 | 0.1 | 0.479 | 0.497 | 0.84 | 0.06 | 0.84 | 0.05 | -0.04 | -0.2 | 0.427 | 0.474 |  |  |
| Somatomotor A: 3 R | 0.74 | 0.20 | 0.65 | 0.22 | 0.45 | 2.0 | 0.025 | 0.061 | 0.81 | 0.05 | 0.82 | 0.05 | -0.14 | -0.6 | 0.263 | 0.382 |  |  |
| Somatomotor A: 4 R | 0.75 | 0.17 | 0.61 | 0.20 | 0.72 | 3.3 | 0.001 | 0.005 | 0.83 | 0.06 | 0.83 | 0.04 | -0.03 | -0.1 | 0.449 | 0.483 |  |  |
| Somatomotor B: Auditory 1 R | 0.78 | 0.11 | 0.76 | 0.07 | 0.13 | 0.6 | 0.272 | 0.353 | 0.80 | 0.06 | 0.80 | 0.06 | -0.04 | -0.2 | 0.431 | 0.474 |  |  |
| Somatomotor B: S2 1 R | 0.64 | 0.27 | 0.71 | 0.16 | -0.33 | -1.4 | 0.080 | 0.152 | 0.80 | 0.07 | 0.82 | 0.06 | -0.28 | -1.3 | 0.101 | 0.280 |  |  |
| Somatomotor B: S2 2 R | 0.73 | 0.24 | 0.76 | 0.12 | -0.14 | -0.7 | 0.257 | 0.342 | 0.81 | 0.07 | 0.82 | 0.07 | -0.25 | -1.2 | 0.124 | 0.280 |  |  |
| Somatomotor B: Central 1 R | 0.73 | 0.24 | 0.70 | 0.17 | 0.17 | 0.8 | 0.225 | 0.316 | 0.89 | 0.06 | 0.88 | 0.06 | 0.14 | 0.6 | 0.260 | 0.382 |  |  |
| Dorsal Attention A: Temporal Occipital 1 R | 0.65 | 0.25 | 0.68 | 0.15 | -0.14 | -0.7 | 0.256 | 0.342 | 0.77 | 0.06 | 0.80 | 0.05 | -0.59 | -2.7 | 0.004 | 0.059 |  |  |
| Dorsal Attention A: Parietal Occipital 1 R | 0.73 | 0.17 | 0.73 | 0.12 | -0.01 | 0.0 | 0.485 | 0.497 | 0.75 | 0.06 | 0.76 | 0.05 | -0.12 | -0.6 | 0.286 | 0.391 |  |  |
| Dorsal Attention A: Superior Parietal Lobule 1 R | 0.75 | 0.18 | 0.75 | 0.09 | 0.01 | 0.0 | 0.490 | 0.497 | 0.75 | 0.09 | 0.79 | 0.05 | -0.61 | -2.8 | 0.003 | 0.059 |  |  |
| Dorsal Attention B: Post Central 1 R | 0.66 | 0.27 | 0.63 | 0.21 | 0.13 | 0.6 | 0.280 | 0.359 | 0.81 | 0.06 | 0.83 | 0.05 | -0.23 | -1.1 | 0.142 | 0.285 |  |  |
| Dorsal Attention B: Post Central 2 R | 0.77 | 0.17 | 0.70 | 0.16 | 0.38 | 1.8 | 0.041 | 0.083 | 0.78 |  |  |  |  |  |  |  |  |  |

Supplementary Table 4 Mean, standard deviation and effect sizes (Cohen's D) of regional betweenness centrality for younger and older adults from the Schaefer Atlas, and t-tests of age group differences.

|  | fPET |  |  |  |  |  |  |  | fMRI |  |  |  |  |  |  |  |
| --- | --- | --- | --- | --- | --- | --- | --- | --- | --- | --- | --- | --- | --- | --- | --- | --- |
|  | Younger |  | Older |  | Cohen's D | Younger vs Older |  |  | Younger |  | Older |  | Cohen's D | Younger vs Older |  |  |
|  | Mean | SD | Mean | SD |  | t-value | p | p-FDR | Mean | SD | Mean | SD |  | t-value | p | p-FDR |
| Visual Central: Extra Striate Cortex 1 L | 0.008 | 0.012 | 0.010 | 0.008 | -0.26 | -1.2 | 0.115 | 0.244 | 0.006 | 0.007 | 0.005 | 0.006 | 0.19 | 0.9 | 0.189 | 0.411 |
| Visual Central: Extra Striate Cortex 2 L | 0.009 | 0.008 | 0.014 | 0.012 | -0.52 | -2.4 | 0.009 | 0.072 | 0.004 | 0.005 | 0.004 | 0.005 | -0.01 | 0.0 | 0.483 | 0.494 |
| Visual Central: Striate Cortex 1 L | 0.005 | 0.005 | 0.009 | 0.010 | -0.51 | -2.4 | 0.010 | 0.072 | 0.006 | 0.009 | 0.006 | 0.006 | 0.00 | 0.0 | 0.494 | 0.495 |
| Visual Central: Extra Striate Cortex 3 L | 0.006 | 0.006 | 0.009 | 0.005 | -0.46 | -2.1 | 0.019 | 0.097 | 0.006 | 0.005 | 0.007 | 0.004 | -0.22 | -1.0 | 0.157 | 0.397 |
| Visual Peripheral: Extra Striate Inferior 1 L | 0.004 | 0.005 | 0.004 | 0.004 | 0.05 | 0.2 | 0.417 | 0.465 | 0.007 | 0.006 | 0.006 | 0.004 | 0.13 | 0.6 | 0.276 | 0.417 |
| Visual Peripheral: Striate Cortex Calcarine 1 L | 0.008 | 0.010 | 0.006 | 0.005 | 0.32 | 1.5 | 0.069 | 0.185 | 0.006 | 0.004 | 0.007 | 0.005 | -0.35 | -1.6 | 0.055 | 0.306 |
| Visual Peripheral: Extra Striate CortexSup 1 L | 0.005 | 0.006 | 0.005 | 0.005 | 0.06 | 0.3 | 0.389 | 0.450 | 0.007 | 0.005 | 0.007 | 0.004 | -0.09 | -0.4 | 0.334 | 0.432 |
| Somatomotor A: 1 L | 0.007 | 0.007 | 0.009 | 0.011 | -0.32 | -1.5 | 0.070 | 0.185 | 0.006 | 0.005 | 0.007 | 0.005 | -0.31 | -1.4 | 0.077 | 0.345 |
| Somatomotor A: 2 L | 0.009 | 0.008 | 0.009 | 0.007 | 0.04 | 0.2 | 0.419 | 0.465 | 0.005 | 0.006 | 0.008 | 0.005 | -0.64 | -3.0 | 0.002 | 0.095 |
| Somatomotor B: Auditory 1 L | 0.013 | 0.009 | 0.012 | 0.007 | 0.16 | 0.7 | 0.231 | 0.335 | 0.008 | 0.005 | 0.009 | 0.006 | -0.17 | -0.8 | 0.212 | 0.411 |
| Somatomotor B: S2 1 L | 0.002 | 0.002 | 0.003 | 0.003 | -0.51 | -2.4 | 0.010 | 0.072 | 0.010 | 0.006 | 0.007 | 0.005 | 0.49 | 2.3 | 0.013 | 0.142 |
| Somatomotor B: S2 2 L | 0.005 | 0.005 | 0.007 | 0.005 | -0.55 | -2.6 | 0.006 | 0.068 | 0.009 | 0.005 | 0.007 | 0.004 | 0.57 | 2.6 | 0.005 | 0.097 |
| Somatomotor B: Central 1 L | 0.004 | 0.006 | 0.006 | 0.005 | -0.26 | -1.2 | 0.114 | 0.244 | 0.003 | 0.004 | 0.003 | 0.003 | -0.15 | -0.7 | 0.251 | 0.411 |
| Dorsal Attention A: Temporal Occipital 1 L | 0.010 | 0.007 | 0.007 | 0.006 | 0.35 | 1.6 | 0.057 | 0.175 | 0.014 | 0.008 | 0.012 | 0.006 | 0.39 | 1.8 | 0.036 | 0.301 |
| Dorsal Attention A: Parietal Occipital 1 L | 0.005 | 0.006 | 0.005 | 0.004 | -0.01 | 0.0 | 0.490 | 0.495 | 0.012 | 0.007 | 0.011 | 0.008 | 0.10 | 0.5 | 0.320 | 0.432 |
| Dorsal Attention A: Superior Parietal Lobule 1 L | 0.009 | 0.008 | 0.010 | 0.006 | -0.10 | -0.5 | 0.315 | 0.396 | 0.010 | 0.006 | 0.011 | 0.006 | -0.29 | -1.3 | 0.090 | 0.348 |
| Dorsal Attention B: Post Central 1 L | 0.005 | 0.006 | 0.005 | 0.005 | 0.02 | 0.1 | 0.470 | 0.495 | 0.009 | 0.007 | 0.007 | 0.004 | 0.35 | 1.6 | 0.053 | 0.306 |
| Dorsal Attention B: Post Central 2 L | 0.003 | 0.004 | 0.005 | 0.008 | -0.39 | -1.8 | 0.037 | 0.131 | 0.005 | 0.004 | 0.006 | 0.004 | -0.18 | -0.8 | 0.205 | 0.411 |
| Dorsal Attention B: Post Central 3 L | 0.005 | 0.005 | 0.005 | 0.005 | -0.07 | -0.3 | 0.376 | 0.447 | 0.008 | 0.005 | 0.011 | 0.007 | -0.38 | -1.8 | 0.040 | 0.306 |
| Dorsal Attention B: Frontal Eye Fields 1 L | 0.008 | 0.008 | 0.005 | 0.005 | 0.45 | 2.1 | 0.019 | 0.097 | 0.012 | 0.008 | 0.014 | 0.011 | -0.13 | -0.6 | 0.268 | 0.417 |
| Saliency Ventral Attention A: Parietal Operculum 1 L | 0.007 | 0.006 | 0.005 | 0.005 | 0.30 | 1.4 | 0.085 | 0.196 | 0.009 | 0.006 | 0.009 | 0.005 | 0.07 | 0.3 | 0.374 | 0.432 |
| Saliency Ventral Attention A: Insula: 1 L | 0.001 | 0.002 | 0.002 | 0.003 | -0.38 | -1.8 | 0.040 | 0.139 | 0.005 | 0.005 | 0.005 | 0.006 | 0.03 | 0.2 | 0.440 | 0.473 |
| Saliency Ventral Attention A: Insula: 2 L | 0.005 | 0.007 | 0.005 | 0.004 | 0.12 | 0.5 | 0.293 | 0.386 | 0.008 | 0.005 | 0.007 | 0.006 | 0.14 | 0.7 | 0.256 | 0.411 |
| Saliency Ventral Attention A: Parietal Medial 1 L | 0.004 | 0.007 | 0.005 | 0.004 | -0.12 | -0.6 | 0.284 | 0.378 | 0.013 | 0.009 | 0.016 | 0.009 | -0.28 | -1.3 | 0.102 | 0.363 |
| Saliency Ventral Attention A: Frontal Medial 1 L | 0.005 | 0.008 | 0.006 | 0.004 | -0.03 | -0.2 | 0.439 | 0.483 | 0.007 | 0.007 | 0.009 | 0.008 | -0.25 | -1.1 | 0.127 | 0.373 |
| Saliency Ventral Attention B: Lateral Prefrontal Cortex 1 L | 0.013 | 0.010 | 0.010 | 0.006 | 0.40 | 1.9 | 0.034 | 0.129 | 0.005 | 0.005 | 0.009 | 0.008 | -0.55 | -2.5 | 0.007 | 0.097 |
| Saliency Ventral Attention B: Medial Posterior Prefrontal 1 L | 0.006 | 0.006 | 0.004 | 0.004 | 0.23 | 1.1 | 0.147 | 0.266 | 0.009 | 0.008 | 0.013 | 0.013 | -0.31 | -1.4 | 0.080 | 0.345 |
| Limbic A: Temporal Pole 1 L | 0.014 | 0.012 | 0.009 | 0.006 | 0.57 | 2.6 | 0.005 | 0.068 | 0.004 | 0.003 | 0.003 | 0.004 | 0.43 | 2.0 | 0.026 | 0.236 |
| Limbic A: Temporal Pole 2 L | 0.004 | 0.005 | 0.003 | 0.003 | 0.24 | 1.1 | 0.138 | 0.263 | 0.006 | 0.005 | 0.005 | 0.005 | 0.14 | 0.7 | 0.252 | 0.411 |
| Limbic B: Orbital Frontal Cortex 1 L | 0.011 | 0.007 | 0.006 | 0.006 | 0.69 | 3.2 | 0.001 | 0.049 | 0.007 | 0.005 | 0.006 | 0.006 | 0.23 | 1.1 | 0.143 | 0.397 |
| Control A: Intraparietal Sulcus 1 L | 0.011 | 0.009 | 0.009 | 0.007 | 0.25 | 1.2 | 0.124 | 0.252 | 0.008 | 0.006 | 0.009 | 0.006 | -0.08 | -0.4 | 0.355 | 0.432 |
| Control A: Lateral Prefrontal Cortex 1 L | 0.017 | 0.008 | 0.014 | 0.009 | 0.34 | 1.6 | 0.058 | 0.175 | 0.005 | 0.004 | 0.007 | 0.006 | -0.27 | -1.2 | 0.109 | 0.363 |
| Control A: Lateral Prefrontal Cortex 2 L | 0.006 | 0.007 | 0.007 | 0.009 | -0.14 | -0.6 | 0.259 | 0.359 | 0.009 | 0.006 | 0.007 | 0.005 | 0.36 | 1.7 | 0.051 | 0.306 |
| Control B: Lateral Prefrontal Cortex 1 L | 0.019 | 0.012 | 0.014 | 0.010 | 0.49 | 2.3 | 0.013 | 0.084 | 0.008 | 0.006 | 0.007 | 0.006 | 0.07 | 0.3 | 0.366 | 0.432 |
| Control C: Precuneus 1 L | 0.008 | 0.006 | 0.010 | 0.007 | -0.22 | -1.0 | 0.159 | 0.270 | 0.005 | 0.004 | 0.006 | 0.005 | -0.08 | -0.4 | 0.358 | 0.432 |
| Control C: Precuneus 2 L | 0.007 | 0.008 | 0.007 | 0.005 | 0.06 | 0.3 | 0.386 | 0.450 | 0.008 | 0.006 | 0.011 | 0.008 | -0.55 | -2.5 | 0.006 | 0.097 |
| Control C: Cingulate Posterior 1 L | 0.005 | 0.004 | 0.006 | 0.004 | -0.34 | -1.6 | 0.061 | 0.176 | 0.005 | 0.005 | 0.006 | 0.005 | -0.19 | -0.9 | 0.195 | 0.411 |
| Default A: Dorsal Prefrontal Cortex 1 L | 0.007 | 0.004 | 0.005 | 0.005 | 0.27 | 1.2 | 0.108 | 0.240 | 0.006 | 0.004 | 0.007 | 0.005 | -0.15 | -0.7 | 0.240 | 0.411 |
| Default A: Precuneus Posterior Cingulate Cortex 1 L | 0.011 | 0.010 | 0.013 | 0.009 | -0.22 | -1.0 | 0.156 | 0.269 | 0.009 | 0.005 | 0.008 | 0.004 | 0.16 | 0.7 | 0.235 | 0.411 |
| Default A: Medial Prefrontal Cortex 1 L | 0.008 | 0.008 | 0.006 | 0.005 | 0.31 | 1.5 | 0.075 | 0.187 | 0.009 | 0.006 | 0.007 | 0.005 | 0.30 | 1.4 | 0.083 | 0.345 |
| Default B: Temp 1 L | 0.006 | 0.007 | 0.005 | 0.004 | 0.26 | 1.2 | 0.117 | 0.244 | 0.012 | 0.007 | 0.011 | 0.008 | 0.07 | 0.3 | 0.380 | 0.432 |
| Default B: Temp 2 L | 0.005 | 0.004 | 0.006 | 0.006 | -0.10 | -0.5 | 0.327 | 0.403 | 0.009 | 0.006 | 0.008 | 0.006 | 0.12 | 0.6 | 0.285 | 0.417 |
| Default B: Inferior Parietal Lobule 1 L | 0.015 | 0.012 | 0.011 | 0.007 | 0.42 | 1.9 | 0.028 | 0.116 | 0.012 | 0.009 | 0.009 | 0.008 | 0.32 | 1.5 | 0.069 | 0.344 |
| Default B: Dorsal Prefrontal Cortex 1 L | 0.020 | 0.011 | 0.011 | 0.008 | 0.94 | 4.3 | 0.000 | 0.002 | 0.010 | 0.007 | 0.011 | 0.007 | -0.12 | -0.6 | 0.288 | 0.417 |
| Default B: Lateral Prefrontal Cortex 1 L | 0.008 | 0.007 | 0.008 | 0.007 | 0.01 | 0.1 | 0.477 | 0.495 | 0.008 | 0.006 | 0.008 | 0.004 | -0.16 | -0.7 | 0.235 | 0.411 |
| Default B: Ventral Prefrontal Cortex 1 L | 0.004 | 0.004 | 0.004 | 0.005 | -0.16 | -0.7 | 0.229 | 0.335 | 0.008 | 0.005 | 0.008 | 0.008 | -0.07 | -0.3 | 0.375 | 0.432 |
| Default B: Ventral Prefrontal Cortex 2 L | 0.018 | 0.011 | 0.016 | 0.010 | 0.17 | 0.8 | 0.211 | 0.324 | 0.009 | 0.006 | 0.007 | 0.005 | 0.30 | 1.4 | 0.081 | 0.345 |
| Default C: RetroSuperior Parietal Lobule 1 L | 0.004 | 0.007 | 0.005 | 0.004 | -0.14 | -0.6 | 0.264 | 0.361 | 0.008 | 0.007 | 0.006 | 0.004 | 0.27 | 1.3 | 0.107 | 0.363 |
| Default C: Parahippocampal Cortex 1 L | 0.001 | 0.002 | 0.003 | 0.004 | -0.56 | -2.6 | 0.006 | 0.068 | 0.006 | 0.006 | 0.006 | 0.007 | 0.00 | 0.0 | 0.495 | 0.495 |
| Temporal Parietal 1 L | 0.004 | 0.005 | 0.006 | 0.006 | -0.48 | -2.2 | 0.015 | 0.093 | 0.009 | 0.007 | 0.010 | 0.007 | -0.10 | -0.5 | 0.326 | 0.432 |
| Visual Central: Extra Striate Cortex 1 R | 0.006 | 0.007 | 0.007 | 0.005 | -0.10 | -0.5 | 0.317 | 0.396 | 0.007 | 0.007 | 0.005 | 0.003 | 0.25 | 1.1 | 0.127 | 0.373 |
| Visual Central: Extra Striate Cortex 2 R | 0.013 | 0.011 | 0.015 | 0.011 | -0.24 | -1.1 | 0.139 | 0.263 | 0.003 | 0.003 | 0.003 | 0.003 | -0.01 | 0.0 | 0.484 | 0.494 |
| Visual Central: Extra Striate Cortex 3 R | 0.006 | 0.010 | 0.009 | 0.007 | -0.40 | -1.8 | 0.035 | 0.130 | 0.005 | 0.004 | 0.005 | 0.003 | 0.16 | 0.7 | 0.232 | 0.411 |
| Visual Peripheral: Striate Cortex Calcarine 1 R | 0.005 | 0.005 | 0.007 | 0.007 | -0.22 | -1.0 | 0.153 | 0.269 | 0.004 | 0.004 | 0.005 | 0.004 | -0.22 | -1.0 | 0.159 | 0.397 |
| Visual Peripheral: Extra Striate Inferior 1 R | 0.004 | 0.006 | 0.005 | 0.004 | -0.02 | -0.1 | 0.472 | 0.495 | 0.009 | 0.007 | 0.009 | 0.007 | -0.06 | -0.3 | 0.398 | 0.435 |
| Visual Peripheral: Extra Striate Superior 1 R | 0.005 | 0.006 | 0.005 | 0.005 | 0.11 | 0.5 | 0.311 | 0.396 | 0.006 | 0.004 | 0.006 | 0.005 | -0.16 | -0.7 | 0.231 | 0.411 |
| Somatomotor A: 1 R | 0.003 | 0.005 | 0.004 | 0.004 | -0.15 | -0.7 | 0.241 | 0.344 | 0.008 | 0.008 | 0.009 | 0.006 | -0.13 | -0.6 | 0.275 | 0.417 |
| Somatomotor A: 2 R | 0.004 | 0.005 | 0.006 | 0.005 | -0.46 | -2.1 | 0.017 | 0.097 | 0.004 | 0.004 | 0.005 | 0.003 | -0.08 | -0.4 | 0.357 | 0.432 |
| Somatomotor A: 3 R | 0.003 | 0.004 | 0.003 | 0.003 | -0.14 | -0. |  |  |  |  |  |  |  |  |  |  |

Supplementary Table 5. Mean, standard deviation and effect sizes (Cohen's D) of regional degree for younger and older adults in fPET and fMRI from the Schaefer Atlas, and t-tests of age group differences.

|  | fPET |  |  |  |  |  |  |  |  | fMRI |  |  |  |  |  |  |  |  |
| --- | --- | --- | --- | --- | --- | --- | --- | --- | --- | --- | --- | --- | --- | --- | --- | --- | --- | --- |
|  | Younger |  | Older |  | Younger vs Older |  |  |  | Cohen's D | Younger |  | Older |  | Younger vs Older |  |  |  | Cohen's D |
|  | Mean | SD | Mean | SD | t-value | p | p-FDR |  |  | Mean | SD | Mean | SD | t-value | p | p-FDR |  |  |
| Visual Central: Extra Striate Cortex 1 L | 25.4 | 17.0 | 34.0 | 16.6 | -0.51 | -2.4 | 0.010 | 0.038 |  | 26.4 | 11.0 | 28.5 | 8.3 | -0.21 | -1.0 | 0.164 | 0.231 |  |
| Visual Central: Extra Striate Cortex 2 L | 30.0 | 18.6 | 40.3 | 18.3 | -0.56 | -2.6 | 0.006 | 0.027 |  | 21.2 | 11.2 | 25.4 | 9.5 | -0.41 | -1.9 | 0.030 | 0.071 |  |
| Visual Central: Striate Cortex 1 L | 24.1 | 18.2 | 29.7 | 16.0 | -0.33 | -1.5 | 0.066 | 0.126 |  | 20.4 | 8.8 | 24.7 | 9.1 | -0.48 | -2.2 | 0.014 | 0.047 |  |
| Visual Central: Extra Striate Cortex 3 L | 23.6 | 16.6 | 32.5 | 14.3 | -0.58 | -2.7 | 0.005 | 0.024 |  | 26.1 | 7.5 | 34.0 | 8.4 | -0.98 | -4.5 | 0.000 | 0.000 |  |
| Visual Peripheral: Extra Striate Inferior 1 L | 15.5 | 11.0 | 19.6 | 11.2 | -0.37 | -1.7 | 0.045 | 0.116 |  | 29.9 | 11.8 | 33.5 | 10.2 | -0.33 | -1.5 | 0.064 | 0.123 |  |
| Visual Peripheral: Striate Cortex Calcarine 1 L | 24.2 | 15.2 | 21.9 | 12.7 | 0.16 | 0.8 | 0.225 | 0.308 |  | 26.4 | 10.7 | 33.7 | 10.8 | -0.68 | -3.1 | 0.001 | 0.007 |  |
| Visual Peripheral: Extra Striate Cortex Sup 1 L | 22.6 | 15.3 | 22.3 | 11.5 | 0.02 | 0.1 | 0.462 | 0.481 |  | 28.8 | 10.6 | 36.2 | 8.8 | -0.76 | -3.5 | 0.000 | 0.004 |  |
| Somatomotor A: 1 L | 31.4 | 17.3 | 31.0 | 16.8 | 0.02 | 0.1 | 0.455 | 0.479 |  | 32.2 | 13.3 | 38.2 | 9.3 | -0.53 | -2.4 | 0.008 | 0.033 |  |
| Somatomotor A: 2 L | 33.5 | 19.4 | 32.2 | 16.4 | 0.07 | 0.3 | 0.366 | 0.419 |  | 32.1 | 11.8 | 40.3 | 8.4 | -0.81 | -3.7 | 0.000 | 0.003 |  |
| Somatomotor B: Auditory 1 L | 40.5 | 15.0 | 40.2 | 14.7 | 0.02 | 0.1 | 0.471 | 0.486 |  | 33.9 | 13.1 | 35.9 | 12.1 | -0.16 | -0.7 | 0.234 | 0.287 |  |
| Somatomotor B: S2 1 L | 10.5 | 9.6 | 18.9 | 10.5 | -0.84 | -3.9 | 0.000 | 0.002 |  | 36.1 | 11.7 | 33.3 | 12.8 | 0.23 | 1.0 | 0.149 | 0.216 |  |
| Somatomotor B: S2 2 L | 21.6 | 15.7 | 29.2 | 13.5 | -0.52 | -2.4 | 0.009 | 0.037 |  | 37.2 | 11.8 | 34.6 | 9.2 | 0.25 | 1.1 | 0.129 | 0.198 |  |
| Somatomotor B: Central 1 L | 22.5 | 16.4 | 25.5 | 15.5 | -0.19 | -0.9 | 0.192 | 0.285 |  | 22.0 | 13.1 | 29.5 | 11.5 | -0.62 | -2.8 | 0.003 | 0.013 |  |
| Dorsal Attention A: Temporal Occipital 1 L | 25.4 | 17.4 | 28.1 | 13.0 | -0.18 | -0.8 | 0.208 | 0.292 |  | 38.2 | 11.3 | 35.2 | 8.5 | 0.30 | 1.4 | 0.085 | 0.149 |  |
| Dorsal Attention A: Parietal Occipital 1 L | 17.1 | 13.5 | 22.2 | 11.8 | -0.41 | -1.9 | 0.032 | 0.083 |  | 39.7 | 11.0 | 37.0 | 13.6 | 0.21 | 1.0 | 0.164 | 0.231 |  |
| Dorsal Attention A: Superior Parietal Lobule 1 L | 37.2 | 13.3 | 36.0 | 13.5 | 0.09 | 0.4 | 0.343 | 0.408 |  | 32.6 | 9.0 | 37.5 | 9.0 | -0.54 | -2.5 | 0.007 | 0.027 |  |
| Dorsal Attention B: Post Central 1 L | 24.7 | 14.9 | 21.2 | 13.1 | 0.25 | 1.1 | 0.129 | 0.212 |  | 35.3 | 12.1 | 38.2 | 7.1 | -0.30 | -1.4 | 0.084 | 0.149 |  |
| Dorsal Attention B: Post Central 2 L | 17.7 | 13.7 | 20.4 | 12.4 | -0.21 | -1.0 | 0.165 | 0.253 |  | 31.1 | 13.1 | 34.8 | 8.4 | -0.34 | -1.6 | 0.061 | 0.120 |  |
| Dorsal Attention B: Post Central 3 L | 27.8 | 15.8 | 24.2 | 12.8 | 0.25 | 1.2 | 0.124 | 0.207 |  | 35.2 | 9.6 | 41.1 | 8.5 | -0.66 | -3.0 | 0.002 | 0.009 |  |
| Dorsal Attention B: Frontal Eye Fields 1 L | 36.4 | 17.3 | 26.2 | 15.7 | 0.62 | 2.9 | 0.002 | 0.015 |  | 35.5 | 12.9 | 37.5 | 11.4 | -0.17 | -0.8 | 0.222 | 0.284 |  |
| Saliency Ventral Attention A: Parietal Operculum 1 L | 30.8 | 14.2 | 26.3 | 13.1 | 0.33 | 1.5 | 0.065 | 0.126 |  | 35.6 | 10.0 | 32.2 | 10.5 | 0.33 | 1.5 | 0.068 | 0.128 |  |
| Saliency Ventral Attention A: Insula 1 L | 6.8 | 6.1 | 12.6 | 9.1 | -0.74 | -3.4 | 0.000 | 0.005 |  | 28.1 | 12.2 | 23.7 | 11.3 | 0.37 | 1.7 | 0.043 | 0.090 |  |
| Saliency Ventral Attention A: Insula 2 L | 19.7 | 15.4 | 22.3 | 12.1 | -0.19 | -0.9 | 0.194 | 0.285 |  | 30.6 | 10.4 | 25.8 | 9.4 | 0.48 | 2.2 | 0.014 | 0.047 |  |
| Saliency Ventral Attention A: Parietal Medial 1 L | 21.4 | 13.5 | 23.5 | 10.1 | -0.18 | -0.8 | 0.203 | 0.290 |  | 34.6 | 13.7 | 42.7 | 11.6 | -0.64 | -3.0 | 0.002 | 0.010 |  |
| Saliency Ventral Attention A: Frontal Medial 1 L | 27.5 | 16.4 | 25.5 | 13.1 | 0.13 | 0.6 | 0.273 | 0.354 |  | 27.8 | 13.8 | 29.8 | 11.2 | -0.16 | -0.7 | 0.231 | 0.287 |  |
| Saliency Ventral Attention B: Lateral Prefrontal Cortex 1 L | 48.0 | 12.6 | 39.5 | 12.9 | 0.66 | 3.1 | 0.001 | 0.010 |  | 19.9 | 10.1 | 24.8 | 12.1 | -0.44 | -2.0 | 0.023 | 0.063 |  |
| Saliency Ventral Attention B: Medial Posterior Prefrontal 1 L | 29.1 | 14.2 | 22.8 | 13.4 | 0.46 | 2.1 | 0.019 | 0.059 |  | 25.8 | 13.2 | 28.8 | 11.8 | -0.24 | -1.1 | 0.138 | 0.206 |  |
| Limbic A Temporal Pole 1 L | 38.0 | 21.4 | 35.2 | 14.4 | 0.15 | 0.7 | 0.240 | 0.320 |  | 23.0 | 11.5 | 14.0 | 11.2 | 0.79 | 3.7 | 0.000 | 0.003 |  |
| Limbic A: Temporal Pole 2 L | 16.4 | 13.4 | 15.9 | 11.8 | 0.04 | 0.2 | 0.422 | 0.464 |  | 24.5 | 12.6 | 18.6 | 10.9 | 0.51 | 2.3 | 0.011 | 0.040 |  |
| Limbic B: Orbital Frontal Cortex 1 L | 35.7 | 16.5 | 27.3 | 14.4 | 0.54 | 2.5 | 0.007 | 0.033 |  | 27.8 | 11.8 | 19.6 | 10.3 | 0.74 | 3.4 | 0.000 | 0.004 |  |
| Control A: Intraparietal Sulcus 1 L | 40.8 | 12.8 | 36.3 | 11.6 | 0.37 | 1.7 | 0.047 | 0.117 |  | 26.1 | 9.2 | 26.6 | 10.2 | -0.05 | -0.3 | 0.400 | 0.417 |  |
| Control A: Lateral Prefrontal Cortex 1 L | 53.7 | 8.5 | 46.2 | 13.4 | 0.66 | 3.0 | 0.002 | 0.010 |  | 21.8 | 8.8 | 22.2 | 8.8 | -0.05 | -0.2 | 0.414 | 0.421 |  |
| Control A: Lateral Prefrontal Cortex 2 L | 30.7 | 15.6 | 25.9 | 13.4 | 0.33 | 1.5 | 0.064 | 0.126 |  | 29.7 | 11.2 | 24.9 | 9.8 | 0.46 | 2.1 | 0.019 | 0.056 |  |
| Control B: Lateral Prefrontal Cortex 1 L | 52.5 | 13.9 | 46.0 | 12.4 | 0.50 | 2.3 | 0.012 | 0.043 |  | 27.3 | 13.9 | 23.5 | 11.4 | 0.30 | 1.4 | 0.088 | 0.149 |  |
| Control C: Precuneus 1 L | 34.7 | 11.8 | 36.1 | 12.4 | -0.11 | -0.5 | 0.300 | 0.375 |  | 19.7 | 9.0 | 23.6 | 9.9 | -0.41 | -1.9 | 0.030 | 0.071 |  |
| Control C: Precuneus 2 L | 26.0 | 12.5 | 28.4 | 13.5 | -0.18 | -0.8 | 0.202 | 0.290 |  | 24.6 | 11.3 | 34.3 | 11.4 | -0.86 | -4.0 | 0.000 | 0.002 |  |
| Control C: Cingulate Posterior 1 L | 25.5 | 12.6 | 30.0 | 12.4 | -0.36 | -1.7 | 0.050 | 0.120 |  | 21.1 | 11.0 | 22.2 | 8.9 | -0.11 | -0.5 | 0.308 | 0.363 |  |
| Default A: Dorsal Prefrontal Cortex 1 L | 40.2 | 15.8 | 26.6 | 12.9 | 0.95 | 4.4 | 0.000 | 0.001 |  | 23.4 | 10.5 | 23.8 | 12.5 | -0.04 | -0.2 | 0.429 | 0.429 |  |
| Default A: Precuneus Posterior Cingulate Cortex 1 L | 39.9 | 13.9 | 43.9 | 12.2 | -0.31 | -1.4 | 0.079 | 0.146 |  | 27.1 | 10.3 | 25.3 | 8.7 | 0.18 | 0.9 | 0.198 | 0.261 |  |
| Default A: Medial Prefrontal Cortex 1 L | 36.2 | 13.1 | 26.8 | 12.6 | 0.74 | 3.4 | 0.001 | 0.005 |  | 28.0 | 10.7 | 25.4 | 10.0 | 0.25 | 1.2 | 0.125 | 0.196 |  |
| Default B: Temp 1 L | 24.1 | 18.2 | 23.2 | 12.9 | 0.06 | 0.3 | 0.397 | 0.441 |  | 32.6 | 11.4 | 28.9 | 10.2 | 0.34 | 1.6 | 0.060 | 0.120 |  |
| Default B: Temp 2 L | 25.6 | 14.3 | 25.7 | 13.0 | -0.01 | 0.0 | 0.485 | 0.489 |  | 31.1 | 11.8 | 26.4 | 9.9 | 0.43 | 2.0 | 0.024 | 0.065 |  |
| Default B: Inferior Parietal Lobule 1 L | 44.9 | 14.9 | 40.0 | 13.7 | 0.34 | 1.6 | 0.061 | 0.126 |  | 33.0 | 11.1 | 25.3 | 10.2 | 0.73 | 3.4 | 0.001 | 0.005 |  |
| Default B: Dorsal Prefrontal Cortex 1 L | 52.5 | 10.9 | 38.3 | 16.1 | 1.03 | 4.8 | 0.000 | 0.000 |  | 31.1 | 11.4 | 30.2 | 11.4 | 0.07 | 0.3 | 0.368 | 0.402 |  |
| Default B: Lateral Prefrontal Cortex 1 L | 38.3 | 14.7 | 33.3 | 14.3 | 0.34 | 1.6 | 0.059 | 0.125 |  | 27.1 | 12.4 | 26.5 | 10.2 | 0.05 | 0.2 | 0.407 | 0.419 |  |
| Default B: Ventral Prefrontal Cortex 1 L | 23.1 | 13.2 | 22.7 | 11.7 | 0.03 | 0.1 | 0.444 | 0.479 |  | 26.8 | 9.8 | 24.0 | 10.7 | 0.27 | 1.3 | 0.107 | 0.175 |  |
| Default B: Ventral Prefrontal Cortex 2 L | 51.5 | 15.5 | 46.1 | 15.1 | 0.35 | 1.6 | 0.054 | 0.122 |  | 28.0 | 10.7 | 24.1 | 8.8 | 0.40 | 1.8 | 0.035 | 0.075 |  |
| Default C: Retro Superior Parietal Lobule 1 L | 17.8 | 14.4 | 23.7 | 11.9 | -0.45 | -2.1 | 0.020 | 0.060 |  | 24.2 | 9.3 | 25.2 | 8.1 | -0.11 | -0.5 | 0.307 | 0.363 |  |
| Default C: Parahippocampal Cortex 1 L | 6.7 | 6.3 | 13.1 | 9.7 | -0.78 | -3.6 | 0.000 | 0.005 |  | 22.8 | 11.5 | 21.8 | 11.1 | 0.09 | 0.4 | 0.341 | 0.388 |  |
| Temporal Parietal 1 L | 19.3 | 14.0 | 24.0 | 12.2 | -0.36 | -1.7 | 0.051 | 0.120 |  | 32.1 | 12.1 | 33.8 | 11.7 | -0.15 | -0.7 | 0.250 | 0.301 |  |
| Visual Central: Extra Striate Cortex 1 R | 26.2 | 15.7 | 28.4 | 13.5 | -0.15 | -0.7 | 0.246 | 0.324 |  | 25.3 | 7.8 | 31.5 | 8.8 | -0.75 | -3.5 | 0.000 | 0.004 |  |
| Visual Central: Extra Striate Cortex 2 R | 35.2 | 18.6 | 43.3 | 19.3 | -0.43 | -2.0 | 0.026 | 0.071 |  | 19.9 | 9.8 | 22.7 | 9.0 | -0.30 | -1.4 | 0.086 | 0.149 |  |
| Visual Central: Extra Striate Cortex 3 R | 23.4 | 14.9 | 35.8 | 14.3 | -0.85 | -3.9 | 0.000 | 0.002 |  | 24.3 | 8.0 | 30.4 | 7.7 | -0.77 | -3.6 | 0.000 | 0.004 |  |
| Visual Peripheral: Striate Cortex Calcarine 1 R | 25.7 | 17.7 | 27.2 | 15.4 | -0.09 | -0.4 | 0.334 | 0.403 |  | 21.2 | 8.8 | 28.5 | 8.5 | -0.85 | -3.9 | 0.000 | 0.002 |  |
| Visual Peripheral: Extra Striate Inferior 1 R | 16.6 | 10.8 | 20.6 | 12.1 | -0.35 | -1.6 | 0.055 | 0.122 |  | 33.3 | 13.0 | 38.3 | 10.4 | -0.42 | -2.0 | 0.027 | 0.070 |  |
| Visual Peripheral: Extra Striate Superior 1 R | 21.4 | 15.7 | 23.5 | 12.9 | -0.15 | -0.7 | 0.240 | 0.320 |  | 26.5 | 8.3 | 34.0 | 11.3 | -0.75 | -3.5 | 0.000 | 0.004 |  |
| Somatomotor A: 1 R | 12.2 | 9.2 | 19.2 | 11.6 | -0.67 | -3.1 | 0.001 | 0.010 |  | 33.8 | 11.5 | 38.3 | 7.8 | -0.46 | -2.1 | 0.018 | 0.053 |  |
| Somatomotor A: 2 R | 16.4 | 14.7 | 23.5 | 14.2 | -0.49 | -2.3 | 0.012 | 0.043 |  | 32.2 | 12.2 | 33.1 | 9.0 | -0.09 | -0.4 | 0.338 | 0.388 |  |
| Somatomotor A: 3 R | 15.3 | 11.6 | 17.8 | 11.0 | -0.22 | -1.0 | 0.160 | 0.252 |  | 36.3 | 10.9 | 38.4 | 8.1 | -0.23 | -1.1 | 0.147 | 0.216 |  |
| Somatomotor A: 4 R | 25.2 | 17.9 | 26.4 | 16.2 | -0.07 | -0.3 | 0.369 | 0.419 |  | 34.5 | 9.7 | 37.3 | 8.6 | -0.30 | -1.4 | 0.081 | 0.148 |  |
| Somatomotor B: Auditory 1 R | 41.0 | 15.6 | 42.3 | 11.8 | -0.10 | -0.5 | 0.324 | 0.400 |  | 37.8 | 12.2 | 34.8 | 12.1 | 0.25 | 1.2 | 0.125 | 0.196 |  |
| Somatomotor B: S2 1 R | 10.5 | 8.5 | 17.0 | 10.5 | -0.68 | -3.1 | 0.001 | 0.010 |  | 34.9 | 10.9 | 34.0 | 11.6 | 0.08 | 0.4 | 0.360 | 0.400 |  |
| Somatomotor B: S2 2 R | 18.9 | 11.1 | 24.7 | 11.5 | -0.52 | -2.4 | 0.009 | 0.038 |  | 34.8 | 9.8 | 34.3 | 9.7 | 0.05 | 0.2 | 0.417 | 0.421 |  |
| Somatomotor B: Central 1 R | 19.8 | 13.8 | 23.2 | 12.1 | -0.26 | -1.2 | 0.114 | 0.197 |  | 24.5 | 14.5 | 25.5 | 11.9 | -0.08 | -0.4 | 0.354 | 0.397 |  |
| Dorsal Attention A: Temporal Occipital 1 R | 20.9 | 13.7 | 26.9 | 14.4 | -0.43 | -2.0 | 0.025 | 0.071 |  | 37.3 | 9.4 | 34.8 | 10.1 | 0.25 | 1.2 | 0.123 | 0.196 |  |
| Dorsal Attention A: Parietal Occipital 1 R | 26.3 | 14.1 | 32.0 | 12.7 | -0.43 | -2.0 | 0.025 | 0.071 |  | 39.2 | 10.8 | 34.9 | 11.6 | 0.39 | 1.8 | 0.039 | 0.082 |  |
| Dorsal Attention A: Superior Parietal Lobule 1 R | 27.5 | 15.3 | 32.4 | 13.4 | -0.34 | -1.6 | 0.059 | 0.125 |  | 30.3 | 9.4 | 34.5 | 8.4 | -0.47 | -2.2 | 0.016 | 0.049 |  |
| Dorsal Attention B: Post Central 1 R | 15.7 | 12.5 | 18.1 | 11.9 | -0.20 | -0.9 |  |  |  |  |  |  |  |  |  |  |  |  |

Supplementary Table 6. Regression analyses predicting cognitive performance from fPET and fMRI whole brain graph metrics from the Schaefer Atlas and age group. The ANOVA results for the overall models are shown in the top three rows, and the standardized beta weights for age and each graph metric's relationship with cognition in the subsequent rows. Significant beta weights are indicated at \*p < .05 and \*\*p < .001.

|  | HVLT:<br>Delayed<br>Recall | HVLT:<br>Discrim.<br>Index | Digit<br>Span:<br>Forward | Digit<br>Span:<br>Back | Category<br>Switch:<br>% Trials | Category<br>Switch:<br>RT | Dig Sub:<br>Number<br>Correct | Dig Sub:<br>Sec Per<br>Correct | Stop<br>Signal:<br>Pro Reac | Stop<br>Signal:<br>RT |
| --- | --- | --- | --- | --- | --- | --- | --- | --- | --- | --- |
| ANOVA F | 2.3 | 3.5 | 0.2 | 2.0 | 1.9 | 5.5 | 18.3 | 0.7 | 2.5 | 1.7 |
| ANOVA p | 0.028 | 0.004 | 0.743 | 0.077 | 0.043 | 0.001 | 0.001 | 0.865 | 0.134 | 0.193 |
| Variance Explained | 17% | 24% | 1% | 16% | 15% | 33% | 63% | 6% | 18% | 13% |
| fPET: Global Efficiency | 0.04 | 0.21 | 0.05 | -0.07 | -0.17 | 0.14 | 0.02 | -0.17 | -0.06 | 0.00 |
| fPET: Local Efficiency | 0.25 | 0.38* | 0.07 | -0.07 | -0.01 | 0.21 | 0.19 | -0.06 | 0.04 | -0.02 |
| fPET: Betweenness Centrality | 0.08 | 0.12 | 0.10 | 0.30* | 0.10 | 0.01 | 0.04 | 0.06 | -0.17 | -0.09 |
| fMRI: Global Efficiency | 0.11 | -0.23 | -0.06 | 0.46 | 0.27 | 0.09 | 0.05 | -0.03 | -0.94 | -0.09 |
| fMRI: Local Efficiency | -0.03 | -0.05 | 0.03 | -0.06 | -0.09 | -0.02 | -0.11 | 0.10 | -0.11 | 0.05 |
| fMRI: Betweenness Centrality | 0.14 | -0.01 | -0.07 | 0.39 | 0.57 | -0.04 | 0.25 | 0.05 | -0.94* | -0.34 |
| Age Group | -0.28* | -0.30* | 0.09 | -0.12 | -0.18 | -0.49** | -0.69** | -0.15 | -0.19 | -0.29 |

Discrim = discrimination; Dig Sub = Digit Substitution; Pro React = probability of reacting.

#### 3. Anatomical Parcellation: Harvard Oxford Atlas

There is currently no consensus on the optimal atlas to parcellate metabolic data. In fMRI, it is widely accepted that a 'functional' atlas derived from resting-state fMRI data provides a robust estimates and high interpretability of functional connectivity results (e.g., Schaefer atlas [9]). Brain parcellations derived from a single modality (e.g., resting-state fMRI) show varying levels of transferability to other modalities [10] and it is currently untested how transferable fMRI-derived atlases are to FDG-PET data. Previous results from metabolic covariance analyses [11, 12] indicate that fMRI-derived atlases are may not be directly transferable to FDG-PET data. Therefore, we chose to use a fMRI-derived functional atlas for results in the main paper to compare the fPET results to fMRI. However, results for the Harvard Oxford atlas are provided here for comparison. In addition, because PET SNR scales directly with spatial scale and the size of the ROI [13], we chose parcellations with relatively coarse granularity.

We found some differences between the parcellations, such as slightly lower maximum connectivity strength in the functional parcellation but slightly higher topological similarities to fMRI in the anatomical parcellation. The anatomical parcellation appears to be particularly effective at highlighting a reduction in connectivity strength in the frontal regions of the brain in ageing that parallel similar BOLD changes and changes to underlying cerebral metabolic rates of glucose metabolism in ageing [15]. The anatomical parcellation also predicted performance in different aspects of cognition, such as the percentage of correct response inhibition trials and reaction time in the stop signal task. On the basis of these and our previous results [16] at this stage we conclude that fMRI-derived parcellations of fPET data appear to be a valid approach for studying the coherence of dynamic glucose metabolism signals. We encourage the comparison of different parcellation approaches in fPET studies of other populations to further characterise similarities and differences.

Supplementary Table 7. Mean, standard deviation and effect sizes (Cohen's D) of regional global efficiency for younger and older adults in fPET and fMRI from the Harvard Oxford Atlas, and t-tests of age group differences.

|  | fPET |  |  |  |  |  |  |  |  | fMRI |  |  |  |  |  |  |  |  |
| --- | --- | --- | --- | --- | --- | --- | --- | --- | --- | --- | --- | --- | --- | --- | --- | --- | --- | --- |
|  | Younger |  |  | Older |  |  | Younger vs Older |  |  | Younger |  |  | Older |  |  | Younger vs Older |  |  |
|  | Mean | SD |  | Mean | SD | Cohen's D | t-value | p | p-FDR | Mean | SD |  | Mean | SD | Cohen's D | t-value | p | p-FDR |
| Frontal Orbital Cortex L | 0.71 | 0.06 | 0.67 | 0.06 | 0.55 | 2.6 | 0.006 | 0.038 |  | 0.68 | 0.07 | 0.64 | 0.09 | 0.48 | 2.2 | 0.015 | 0.054 |  |
| Frontal Pole L | 0.82 | 0.04 | 0.79 | 0.05 | 0.72 | 3.3 | 0.001 | 0.010 |  | 0.65 | 0.08 | 0.64 | 0.08 | 0.42 | 0.7 | 0.232 | 0.345 |  |
| Frontal Operculum Cortex L | 0.62 | 0.06 | 0.63 | 0.06 | -0.12 | -0.5 | 0.295 | 0.406 |  | 0.63 | 0.07 | 0.62 | 0.07 | 0.16 | 0.6 | 0.288 | 0.368 |  |
| Superior Frontal Gyrus L | 0.73 | 0.08 | 0.68 | 0.08 | 0.66 | 3.1 | 0.002 | 0.018 |  | 0.63 | 0.08 | 0.64 | 0.06 | 0.12 | -0.3 | 0.381 | 0.444 |  |
| Middle Frontal Gyrus L | 0.76 | 0.04 | 0.72 | 0.06 | 0.72 | 3.3 | 0.001 | 0.010 |  | 0.61 | 0.07 | 0.62 | 0.07 | 0.12 | -0.7 | 0.241 | 0.345 |  |
| Inferior Frontal Gyrus; Pars Triangularis L | 0.70 | 0.06 | 0.68 | 0.05 | 0.34 | 1.6 | 0.058 | 0.143 |  | 0.62 | 0.07 | 0.61 | 0.06 | -0.07 | 0.7 | 0.237 | 0.345 |  |
| Inferior Frontal Gyrus; pars opercularis L | 0.67 | 0.07 | 0.64 | 0.06 | 0.46 | 2.1 | 0.019 | 0.074 |  | 0.63 | 0.06 | 0.62 | 0.07 | -0.15 | 0.7 | 0.257 | 0.349 |  |
| Paracingulate Gyrus L | 0.71 | 0.05 | 0.65 | 0.06 | 1.02 | 4.7 | 0.000 | 0.000 |  | 0.61 | 0.08 | 0.63 | 0.07 | -0.15 | -1.0 | 0.169 | 0.280 |  |
| Insular Cortex L | 0.57 | 0.11 | 0.58 | 0.10 | -0.09 | -0.4 | 0.342 | 0.409 |  | 0.66 | 0.07 | 0.66 | 0.05 | 0.15 | -0.4 | 0.334 | 0.408 |  |
| Amygdala l | 0.54 | 0.09 | 0.56 | 0.06 | -0.31 | -1.4 | 0.076 | 0.168 |  | 0.60 | 0.08 | 0.52 | 0.17 | 0.14 | 2.8 | 0.003 | 0.017 |  |
| Juxtapositional Lobule Cortex - L | 0.62 | 0.08 | 0.61 | 0.12 | 0.08 | 0.4 | 0.350 | 0.409 |  | 0.63 | 0.07 | 0.66 | 0.05 | 0.14 | -2.6 | 0.005 | 0.028 |  |
| Precentral Gyrus L | 0.72 | 0.08 | 0.69 | 0.07 | 0.34 | 1.6 | 0.061 | 0.147 |  | 0.67 | 0.09 | 0.71 | 0.05 | -0.21 | -2.4 | 0.009 | 0.040 |  |
| PostCG l Postcentral Gyrus L | 0.70 | 0.08 | 0.70 | 0.08 | -0.05 | -0.2 | 0.418 | 0.443 |  | 0.66 | 0.08 | 0.70 | 0.04 | -0.09 | -3.4 | 0.001 | 0.006 |  |
| Central Opercular Cortex L | 0.59 | 0.07 | 0.63 | 0.07 | -0.55 | -2.6 | 0.006 | 0.038 |  | 0.67 | 0.07 | 0.67 | 0.06 | -0.11 | 0.1 | 0.468 | 0.491 |  |
| Superior Parietal Lobule L | 0.67 | 0.09 | 0.66 | 0.06 | 0.21 | 1.0 | 0.164 | 0.285 |  | 0.64 | 0.07 | 0.69 | 0.05 | 0.61 | -3.9 | 0.000 | 0.003 |  |
| Supramarginal Gyrus; Anterior Division L | 0.66 | 0.07 | 0.63 | 0.06 | 0.53 | 2.4 | 0.008 | 0.044 |  | 0.63 | 0.06 | 0.64 | 0.05 | -0.57 | -0.9 | 0.173 | 0.282 |  |
| Supramarginal Gyrus; Posterior Division L | 0.69 | 0.06 | 0.67 | 0.05 | 0.33 | 1.5 | 0.064 | 0.148 |  | 0.65 | 0.07 | 0.66 | 0.07 | -0.69 | -0.4 | 0.327 | 0.408 |  |
| Angular Gyrus L | 0.65 | 0.07 | 0.66 | 0.05 | -0.06 | -0.3 | 0.400 | 0.435 |  | 0.66 | 0.07 | 0.61 | 0.07 | -0.51 | 3.5 | 0.000 | 0.004 |  |
| Parietal Operculum Cortex L | 0.62 | 0.08 | 0.63 | 0.06 | -0.11 | -0.5 | 0.308 | 0.408 |  | 0.68 | 0.06 | 0.68 | 0.07 | -0.74 | 0.0 | 0.500 | 0.500 |  |
| Parahippocampal Gyrus; Anterior Division L | 0.54 | 0.08 | 0.57 | 0.10 | -0.31 | -1.4 | 0.080 | 0.172 |  | 0.61 | 0.09 | 0.60 | 0.11 | -1.11 | 0.4 | 0.343 | 0.409 |  |
| Parahippocampal Gyrus; Posterior Division L | 0.52 | 0.06 | 0.56 | 0.11 | -0.37 | -1.7 | 0.045 | 0.118 |  | 0.59 | 0.09 | 0.59 | 0.09 | 0.02 | 0.0 | 0.488 | 0.498 |  |
| Hippocampus l | 0.56 | 0.09 | 0.56 | 0.11 | -0.04 | -0.2 | 0.422 | 0.443 |  | 0.64 | 0.07 | 0.59 | 0.12 | -0.84 | 2.2 | 0.015 | 0.054 |  |
| Thalamus L | 0.55 | 0.06 | 0.59 | 0.11 | -0.52 | -2.4 | 0.009 | 0.046 |  | 0.60 | 0.10 | 0.59 | 0.10 | -1.10 | 0.6 | 0.269 | 0.356 |  |
| Caudate L | 0.58 | 0.07 | 0.63 | 0.11 | -0.55 | -2.5 | 0.007 | 0.038 |  | 0.56 | 0.13 | 0.57 | 0.09 | -0.20 | -0.4 | 0.342 | 0.409 |  |
| Putamen L | 0.63 | 0.08 | 0.65 | 0.12 | -0.19 | -0.9 | 0.190 | 0.315 |  | 0.57 | 0.13 | 0.57 | 0.09 | -0.10 | 0.0 | 0.493 | 0.498 |  |
| Palidum L | 0.53 | 0.07 | 0.58 | 0.10 | -0.62 | -2.8 | 0.003 | 0.027 |  | 0.56 | 0.16 | 0.53 | 0.17 | -0.10 | 0.6 | 0.276 | 0.356 |  |
| Accumbens L | 0.53 | 0.07 | 0.58 | 0.10 | -0.57 | -2.7 | 0.005 | 0.036 |  | 0.54 | 0.14 | 0.47 | 0.21 | 0.76 | 1.8 | 0.040 | 0.112 |  |
| Temporal Pole L | 0.60 | 0.07 | 0.59 | 0.11 | 0.19 | 0.9 | 0.193 | 0.315 |  | 0.69 | 0.05 | 0.67 | 0.06 | 0.00 | 1.5 | 0.068 | 0.155 |  |
| Planum Polare L | 0.53 | 0.07 | 0.57 | 0.10 | -0.41 | -1.9 | 0.029 | 0.092 |  | 0.64 | 0.07 | 0.62 | 0.09 | 0.00 | 1.1 | 0.130 | 0.242 |  |
| Superior Temporal Gyrus; Anterior Division L | 0.55 | 0.07 | 0.58 | 0.06 | -0.47 | -2.2 | 0.017 | 0.072 |  | 0.65 | 0.07 | 0.65 | 0.09 | 0.09 | 0.2 | 0.418 | 0.466 |  |
| Superior Temporal Gyrus; Posterior Division L | 0.60 | 0.07 | 0.63 | 0.05 | -0.50 | -2.3 | 0.011 | 0.055 |  | 0.67 | 0.07 | 0.67 | 0.07 | -0.01 | -0.1 | 0.447 | 0.483 |  |
| Middle Temporal Gyrus; Anterior Division L | 0.59 | 0.07 | 0.59 | 0.06 | 0.06 | 0.3 | 0.398 | 0.435 |  | 0.63 | 0.09 | 0.63 | 0.07 | -0.01 | 0.0 | 0.490 | 0.498 |  |
| Middle Temporal Gyrus; Posterior Division L | 0.72 | 0.07 | 0.67 | 0.07 | 0.76 | 3.5 | 0.000 | 0.009 |  | 0.66 | 0.08 | 0.64 | 0.06 | 0.47 | 1.3 | 0.094 | 0.185 |  |
| Inferior Temporal Gyrus; Anterior Division L | 0.58 | 0.07 | 0.57 | 0.10 | 0.13 | 0.6 | 0.279 | 0.395 |  | 0.63 | 0.06 | 0.59 | 0.08 | 0.13 | 2.2 | 0.015 | 0.054 |  |
| Inferior Temporal Gyrus; Posterior Division L | 0.67 | 0.06 | 0.62 | 0.07 | 0.65 | 3.0 | 0.002 | 0.018 |  | 0.67 | 0.07 | 0.60 | 0.07 | 0.13 | 4.0 | 0.000 | 0.003 |  |
| Planum Temporale L | 0.70 | 0.06 | 0.68 | 0.05 | 0.44 | 2.0 | 0.022 | 0.080 |  | 0.68 | 0.07 | 0.70 | 0.07 | -0.09 | -1.4 | 0.079 | 0.170 |  |
| Heschl's Gyrus L | 0.64 | 0.07 | 0.66 | 0.06 | -0.28 | -1.3 | 0.101 | 0.194 |  | 0.64 | 0.07 | 0.65 | 0.08 | 0.00 | -0.8 | 0.220 | 0.338 |  |
| Temporal Fusiform Cortex; Anterior Division L | 0.54 | 0.06 | 0.57 | 0.11 | -0.28 | -1.3 | 0.101 | 0.194 |  | 0.57 | 0.13 | 0.58 | 0.12 | 0.00 | -0.3 | 0.379 | 0.444 |  |
| Temporal Fusiform Cortex; Posterior Division L | 0.58 | 0.06 | 0.59 | 0.10 | -0.13 | -0.6 | 0.280 | 0.395 |  | 0.67 | 0.08 | 0.66 | 0.05 | 0.13 | 0.8 | 0.214 | 0.334 |  |
| Temporal Occipital Fusiform Cortex L | 0.58 | 0.06 | 0.61 | 0.06 | -0.60 | -2.8 | 0.003 | 0.031 |  | 0.66 | 0.06 | 0.65 | 0.06 | 0.38 | 0.7 | 0.250 | 0.349 |  |
| Middle Temporal Gyrus; Temporoccipital Part L | 0.63 | 0.08 | 0.64 | 0.06 | -0.07 | -0.3 | 0.372 | 0.424 |  | 0.67 | 0.07 | 0.64 | 0.05 | 0.33 | 2.0 | 0.023 | 0.073 |  |
| Inferior Temporal Gyrus; Temporoccipital Part L | 0.60 | 0.07 | 0.61 | 0.05 | -0.20 | -0.9 | 0.175 | 0.298 |  | 0.66 | 0.06 | 0.63 | 0.06 | 0.32 | 2.4 | 0.009 | 0.040 |  |
| Occipital Fusiform Gyrus L | 0.64 | 0.09 | 0.66 | 0.07 | -0.30 | -1.4 | 0.081 | 0.172 |  | 0.62 | 0.07 | 0.65 | 0.06 | 0.25 | -1.8 | 0.039 | 0.111 |  |
| Supracalcarine Cortex L | 0.61 | 0.05 | 0.63 | 0.06 | -0.25 | -1.2 | 0.122 | 0.227 |  | 0.59 | 0.08 | 0.64 | 0.05 | 0.22 | -4.0 | 0.000 | 0.003 |  |
| Cuneal Cortex L | 0.61 | 0.07 | 0.62 | 0.06 | -0.16 | -0.7 | 0.233 | 0.363 |  | 0.61 | 0.08 | 0.67 | 0.06 | 0.04 | -3.8 | 0.000 | 0.003 |  |
| Lingual Gyrus L | 0.61 | 0.12 | 0.63 | 0.12 | -0.11 | -0.5 | 0.300 | 0.406 |  | 0.61 | 0.07 | 0.65 | 0.05 | -0.03 | -2.6 | 0.005 | 0.028 |  |
| Intracalcarine Cortex L | 0.64 | 0.08 | 0.64 | 0.07 | 0.02 | 0.1 | 0.458 | 0.467 |  | 0.58 | 0.08 | 0.64 | 0.06 | -0.03 | -3.7 | 0.000 | 0.003 |  |
| Lateral Occipital Cortex; Superior Division L | 0.77 | 0.05 | 0.75 | 0.06 | 0.39 | 1.8 | 0.039 | 0.112 |  | 0.67 | 0.06 | 0.68 | 0.05 | 0.01 | -0.7 | 0.254 | 0.349 |  |
| Lateral Occipital Cortex; Inferior Division L | 0.65 | 0.09 | 0.68 | 0.07 | -0.34 | -1.6 | 0.062 | 0.147 |  | 0.67 | 0.06 | 0.67 | 0.06 | 0.29 | 0.0 | 0.485 | 0.498 |  |
| Occipital Pole L | 0.69 | 0.08 | 0.70 | 0.08 | -0.15 | -0.7 | 0.238 | 0.366 |  | 0.59 | 0.07 | 0.63 | 0.06 | 0.34 | -3.3 | 0.001 | 0.007 |  |
| Frontal Medial Cortex | 0.71 | 0.08 | 0.68 | 0.06 | 0.35 | 1.6 | 0.056 | 0.140 |  | 0.62 | 0.08 | 0.62 | 0.08 | 0.47 | 0.5 | 0.302 | 0.381 |  |
| Cingulate Gyrus; Anterior Division | 0.70 | 0.08 | 0.65 | 0.13 | 0.46 | 2.1 | 0.018 | 0.074 |  | 0.62 | 0.08 | 0.65 | 0.06 | 0.86 | -2.5 | 0.007 | 0.036 |  |
| Subcallosal Cortex | 0.62 | 0.09 | 0.60 | 0.11 | 0.16 | 0.8 | 0.226 | 0.358 |  | 0.60 | 0.12 | 0.57 | 0.16 | 0.83 | 0.9 | 0.189 | 0.303 |  |
| Cingulate Gyrus; Posterior Division | 0.76 | 0.07 | 0.76 | 0.07 | 0.00 | 0.0 | 0.496 | 0.496 |  | 0.60 | 0.09 | 0.63 | 0.06 | -0.30 | -1.6 | 0.062 | 0.145 |  |
| Precuneus Cortex | 0.80 | 0.06 | 0.80 | 0.05 | -0.09 | -0.4 | 0.341 | 0.409 |  | 0.60 | 0.08 | 0.66 | 0.06 | -0.17 | -3.6 | 0.000 | 0.004 |  |
| Brain-Stem | 0.69 | 0.09 | 0.64 | 0.08 | 0.56 | 2.6 | 0.005 | 0.038 |  | 0.49 | 0.21 | 0.55 | 0.13 | -0.16 | -1.7 | 0.051 | 0.133 |  |
| Frontal Orbital Cortex R | 0.72 | 0.07 | 0.69 | 0.06 | 0.37 | 1.7 | 0.044 | 0.118 |  | 0.66 | 0.06 | 0.64 | 0.07 | -0.07 | 1.6 | 0.053 | 0.133 |  |
| Frontal Pole R | 0.84 | 0.03 | 0.81 | 0.05 | 0.77 | 3.6 | 0.000 | 0.009 |  | 0.66 | 0.08 | 0.63 | 0.07 | 0.17 | 1.7 | 0.044 | 0.120 |  |
| Frontal Operculum Cortex R | 0.64 | 0.07 | 0.66 | 0.06 | -0.29 | -1.4 | 0.089 | 0.183 |  | 0.64 | 0.07 | 0.62 | 0.07 | 0.22 | 1.4 | 0.080 | 0.170 |  |
| Superior Frontal Gyrus R | 0.72 | 0.08 | 0.69 | 0.07 | 0.48 | 2.2 | 0.015 | 0.064 |  | 0.65 | 0.07 | 0.65 | 0.06 | 0.14 | -0.2 | 0.427 | 0.470 |  |
| Middle Frontal Gyrus R | 0.78 | 0.07 | 0.75 | 0.07 | 0.55 | 2.5 | 0.007 | 0.038 |  | 0.62 | 0.07 | 0.61 | 0.06 | 0.44 | 0.2 | 0.430 | 0.470 |  |
| Inferior Frontal Gyrus; Pars Triangularis R | 0.70 | 0.05 | 0.69 | 0.05 | 0.13 | 0.6 | 0.277 | 0.395 |  | 0.65 | 0.07 | 0.63 | 0.05 | 0.51 | 1.4 | 0.077 | 0.170 |  |
| Inferior Frontal Gyrus; Pars Opercularis R | 0.73 | 0.05 | 0.69 | 0.05 | 0.88 | 4.1 | 0.000 | 0.003 |  | 0.65 | 0.07 | 0.61 | 0.06 | 0.52 | 2.3 | 0.013 | 0.051 |  |
| Paracingulate Gyrus R | 0.72 | 0.06 | 0.68 | 0.06 | 0.75 | 3.5 | 0.000 | 0.009 |  | 0.60 | 0.08 | 0.63 | 0.06 | -0.39 | -2.0 | 0.027 | 0.084 |  |
| Insular Cortex R | 0.60 | 0.07 | 0.60 | 0.10 | 0.10 | 0.4 | 0.329 | 0.409 |  | 0.68 | 0.05 | 0.68 | 0.05 | -0.44 | -0.1 | 0.454 | 0.483 |  |
| Amygdala r | 0.56 | 0.12 | 0.58 | 0.07 | -0.23 | -1.1 | 0.144 | 0.261 |  | 0.59 | 0.08 | 0.55 | 0.13 | -0.85 | 1.7 | 0.047 | 0.125 |  |
| Juxtapositional Lobule Cortex R | 0.60 | 0.13 | 0.60 | 0.11 | -0.05 | -0.2 | 0.408 | 0.437 |  | 0.63 | 0.07 | 0.67 | 0.05 | -0.82 | -3.0 | 0.002 | 0.014 |  |
| Precentral Gyrus R | 0.73 | 0.06 | 0.70 | 0.08 | 0.41 | 1.9 | 0.031 | 0.093 |  | 0.69 | 0.07 | 0.70 | 0.04 | -0.96 | -1.0 | 0.158 | 0.269 |  |
| Postcentral Gyrus R | 0.66 | 0.07 | 0.66 | 0.08 | -0.05 | -0.2 | 0.402 | 0.435 |  | 0.6 |  |  |  |  |  |  |  |  |

Supplementary Table 8. Mean, standard deviation and effect sizes (Cohen's D) of regional local efficiency for younger and older adults in fPET and fMRI from the Harvard Oxford Atlas, and t-tests of age group differences.

|  | fPET |  |  |  |  |  |  |  |  | fMRI |  |  |  |  |  |  |  |  |
| --- | --- | --- | --- | --- | --- | --- | --- | --- | --- | --- | --- | --- | --- | --- | --- | --- | --- | --- |
|  | Younger |  | Older |  | Younger vs Older |  |  |  | Cohen's D | Younger |  | Older |  | Younger vs Older |  |  |  | Cohen's D |
|  | Mean | SD | Mean | SD | t-value | p | p-FDR | p |  | Mean | SD | Mean | SD | t-value | p | p-FDR | p |  |
| Frontal Orbital Cortex L | 0.78 | 0.06 | 0.75 | 0.07 | 0.42 | 1.9 | 0.027 | 0.061 | 0.75 | 0.08 | 0.75 | 0.05 | 0.05 | -0.44 | 0.3 | 0.391 | 0.491 |  |
| Frontal Pole L | 0.75 | 0.03 | 0.74 | 0.04 | 0.41 | 1.9 | 0.031 | 0.067 | 0.78 | 0.06 | 0.74 | 0.07 | 0.23 | 2.7 | 0.004 | 0.081 | 0.004 | 0.081 |
| Frontal Operculum Cortex L | 0.73 | 0.14 | 0.72 | 0.11 | 0.13 | 0.6 | 0.273 | 0.340 | 0.78 | 0.06 | 0.76 | 0.06 | 0.06 | 0.6 | 1.3 | 0.096 | 0.356 |  |
| Superior Frontal Gyrus L | 0.78 | 0.03 | 0.74 | 0.10 | 0.58 | 2.7 | 0.004 | 0.019 | 0.75 | 0.09 | 0.76 | 0.05 | 0.07 | -0.1 | 0.471 | 0.495 | 0.471 | 0.495 |
| Middle Frontal Gyrus L | 0.78 | 0.03 | 0.75 | 0.05 | 0.82 | 3.8 | 0.000 | 0.001 | 0.77 | 0.07 | 0.76 | 0.05 | 0.26 | 0.7 | 0.245 | 0.433 | 0.245 | 0.433 |
| Inferior Frontal Gyrus; Pars Triangularis L | 0.80 | 0.07 | 0.75 | 0.08 | 0.72 | 3.3 | 0.001 | 0.005 | 0.77 | 0.08 | 0.77 | 0.07 | -0.32 | 0.0 | 0.500 | 0.500 | 0.500 | 0.500 |
| Inferior Frontal Gyrus; Pars Opercularis L | 0.79 | 0.08 | 0.75 | 0.08 | 0.55 | 2.6 | 0.006 | 0.020 | 0.78 | 0.05 | 0.75 | 0.09 | -0.34 | 1.4 | 0.082 | 0.347 | 0.082 | 0.347 |
| Paracingulate Gyrus L | 0.81 | 0.04 | 0.74 | 0.09 | 1.01 | 4.7 | 0.000 | 0.000 | 0.76 | 0.08 | 0.73 | 0.11 | -0.09 | 1.4 | 0.082 | 0.347 | 0.082 | 0.347 |
| Insular Cortex L | 0.67 | 0.19 | 0.65 | 0.11 | 0.08 | 0.4 | 0.354 | 0.395 | 0.77 | 0.06 | 0.77 | 0.05 | 0.21 | -0.4 | 0.346 | 0.480 | 0.346 | 0.480 |
| Amygdala l | 0.52 | 0.26 | 0.53 | 0.20 | -0.08 | -0.4 | 0.359 | 0.397 | 0.78 | 0.11 | 0.77 | 0.21 | 0.19 | 0.3 | 0.400 | 0.491 | 0.400 | 0.491 |
| Juxtastriatal Lobule Cortex - L | 0.77 | 0.18 | 0.73 | 0.09 | 0.26 | 1.2 | 0.117 | 0.189 | 0.79 | 0.09 | 0.79 | 0.06 | -0.13 | -0.4 | 0.356 | 0.480 | 0.356 | 0.480 |
| Precentral Gyrus L | 0.78 | 0.07 | 0.73 | 0.08 | 0.73 | 3.4 | 0.001 | 0.005 | 0.72 | 0.17 | 0.78 | 0.05 | 0.02 | -2.3 | 0.013 | 0.175 | 0.013 | 0.175 |
| PostCG I Postcentral Gyrus L | 0.78 | 0.09 | 0.72 | 0.12 | 0.57 | 2.6 | 0.005 | 0.020 | 0.75 | 0.17 | 0.78 | 0.04 | 0.03 | -1.3 | 0.097 | 0.356 | 0.097 | 0.356 |
| Central Opercular Cortex L | 0.67 | 0.18 | 0.72 | 0.09 | -0.31 | -1.4 | 0.079 | 0.141 | 0.78 | 0.08 | 0.79 | 0.06 | -0.25 | -1.1 | 0.132 | 0.368 | 0.132 | 0.368 |
| Superior Parietal Lobule L | 0.79 | 0.07 | 0.75 | 0.09 | 0.57 | 2.6 | 0.005 | 0.020 | 0.75 | 0.11 | 0.77 | 0.05 | -0.16 | -0.9 | 0.177 | 0.373 | 0.177 | 0.373 |
| Supramarginal Gyrus; Anterior Division L | 0.81 | 0.08 | 0.73 | 0.09 | 0.97 | 4.5 | 0.000 | 0.000 | 0.80 | 0.07 | 0.80 | 0.07 | -0.19 | -0.4 | 0.353 | 0.480 | 0.353 | 0.480 |
| Supramarginal Gyrus; Posterior Division L | 0.80 | 0.05 | 0.74 | 0.07 | 0.99 | 4.6 | 0.000 | 0.000 | 0.76 | 0.05 | 0.76 | 0.05 | 0.15 | -0.2 | 0.436 | 0.491 | 0.436 | 0.491 |
| Angular Gyrus L | 0.80 | 0.08 | 0.74 | 0.09 | 0.64 | 3.0 | 0.002 | 0.011 | 0.76 | 0.05 | 0.76 | 0.06 | 0.06 | -0.2 | 0.418 | 0.491 | 0.418 | 0.491 |
| Parietal Operculum Cortex L | 0.73 | 0.18 | 0.72 | 0.11 | 0.03 | 0.1 | 0.452 | 0.480 | 0.80 | 0.04 | 0.80 | 0.05 | 0.06 | 0.1 | 0.452 | 0.491 | 0.452 | 0.491 |
| Parahippocampal Gyrus; Anterior Division L | 0.51 | 0.25 | 0.57 | 0.17 | -0.26 | -1.2 | 0.116 | 0.188 | 0.79 | 0.08 | 0.79 | 0.08 | 0.45 | 0.2 | 0.439 | 0.491 | 0.439 | 0.491 |
| Parahippocampal Gyrus; Posterior Division L | 0.53 | 0.28 | 0.57 | 0.19 | -0.16 | -0.7 | 0.229 | 0.307 | 0.78 | 0.16 | 0.79 | 0.15 | 0.30 | -0.4 | 0.358 | 0.480 | 0.358 | 0.480 |
| Hippocampus l | 0.55 | 0.28 | 0.55 | 0.19 | -0.01 | 0.0 | 0.488 | 0.494 | 0.76 | 0.08 | 0.76 | 0.13 | 0.27 | -0.2 | 0.406 | 0.480 | 0.406 | 0.480 |
| Thalamus L | 0.60 | 0.29 | 0.70 | 0.15 | -0.43 | -2.0 | 0.025 | 0.058 | 0.74 | 0.16 | 0.75 | 0.14 | 0.24 | -0.2 | 0.413 | 0.491 | 0.413 | 0.491 |
| Caudate L | 0.64 | 0.22 | 0.71 | 0.07 | -0.42 | -2.0 | 0.027 | 0.061 | 0.75 | 0.12 | 0.80 | 0.09 | 0.52 | -1.9 | 0.030 | 0.250 | 0.030 | 0.250 |
| Putamen L | 0.70 | 0.18 | 0.72 | 0.11 | -0.17 | -0.8 | 0.212 | 0.292 | 0.79 | 0.11 | 0.77 | 0.14 | 0.51 | 0.7 | 0.228 | 0.417 | 0.228 | 0.417 |
| Pallidum L | 0.56 | 0.25 | 0.62 | 0.15 | -0.33 | -1.5 | 0.068 | 0.126 | 0.83 | 0.10 | 0.75 | 0.20 | 0.62 | 2.3 | 0.012 | 0.175 | 0.012 | 0.175 |
| Accumbens L | 0.57 | 0.27 | 0.68 | 0.15 | -0.51 | -2.3 | 0.012 | 0.033 | 0.78 | 0.14 | 0.82 | 0.13 | 0.69 | -1.1 | 0.144 | 0.368 | 0.144 | 0.368 |
| Temporal Pole L | 0.66 | 0.17 | 0.60 | 0.20 | 0.32 | 1.5 | 0.073 | 0.134 | 0.79 | 0.06 | 0.75 | 0.05 | 0.72 | 3.3 | 0.001 | 0.043 | 0.001 | 0.043 |
| Planum Polare L | 0.56 | 0.29 | 0.65 | 0.15 | -0.40 | -1.8 | 0.034 | 0.072 | 0.79 | 0.05 | 0.76 | 0.18 | 0.30 | 1.0 | 0.158 | 0.368 | 0.158 | 0.368 |
| Superior Temporal Gyrus; Anterior Division L | 0.56 | 0.27 | 0.64 | 0.14 | -0.37 | -1.7 | 0.046 | 0.094 | 0.78 | 0.05 | 0.80 | 0.08 | 0.23 | -1.0 | 0.171 | 0.373 | 0.171 | 0.373 |
| Superior Temporal Gyrus; Posterior Division L | 0.73 | 0.17 | 0.73 | 0.09 | 0.02 | 0.1 | 0.465 | 0.488 | 0.77 | 0.05 | 0.77 | 0.05 | 0.22 | 0.2 | 0.433 | 0.491 | 0.433 | 0.491 |
| Middle Temporal Gyrus; Anterior Division L | 0.61 | 0.25 | 0.65 | 0.11 | -0.21 | -1.0 | 0.164 | 0.241 | 0.74 | 0.19 | 0.78 | 0.07 | -0.39 | -1.3 | 0.104 | 0.357 | 0.104 | 0.357 |
| Middle Temporal Gyrus; Posterior Division L | 0.76 | 0.07 | 0.71 | 0.09 | 0.57 | 2.6 | 0.005 | 0.020 | 0.78 | 0.05 | 0.76 | 0.07 | 0.06 | 1.5 | 0.064 | 0.322 | 0.064 | 0.322 |
| Inferior Temporal Gyrus; Anterior Division L | 0.67 | 0.21 | 0.58 | 0.20 | 0.43 | 2.0 | 0.024 | 0.058 | 0.81 | 0.06 | 0.79 | 0.07 | 0.06 | 1.6 | 0.055 | 0.292 | 0.055 | 0.292 |
| Inferior Temporal Gyrus; Posterior Division L | 0.76 | 0.09 | 0.68 | 0.17 | 0.63 | 2.9 | 0.002 | 0.011 | 0.78 | 0.06 | 0.76 | 0.09 | -0.18 | 0.9 | 0.176 | 0.373 | 0.176 | 0.373 |
| Planum Temporale L | 0.79 | 0.06 | 0.75 | 0.09 | 0.52 | 2.4 | 0.010 | 0.029 | 0.77 | 0.06 | 0.78 | 0.05 | 0.50 | -0.6 | 0.275 | 0.463 | 0.275 | 0.463 |
| Heschl's Gyrus L | 0.80 | 0.09 | 0.74 | 0.09 | 0.66 | 3.1 | 0.001 | 0.009 | 0.78 | 0.07 | 0.76 | 0.09 | 0.47 | 1.2 | 0.125 | 0.368 | 0.125 | 0.368 |
| Temporal Fusiform Cortex; Anterior Division L | 0.56 | 0.25 | 0.59 | 0.16 | -0.12 | -0.5 | 0.295 | 0.347 | 0.78 | 0.18 | 0.79 | 0.14 | -0.34 | -0.2 | 0.417 | 0.491 | 0.417 | 0.491 |
| Temporal Fusiform Cortex; Posterior Division L | 0.69 | 0.14 | 0.64 | 0.13 | 0.40 | 1.8 | 0.035 | 0.073 | 0.78 | 0.06 | 0.77 | 0.06 | 0.35 | 0.8 | 0.212 | 0.400 | 0.212 | 0.400 |
| Temporal Occipital Fusiform Cortex L | 0.68 | 0.19 | 0.70 | 0.11 | -0.14 | -0.6 | 0.264 | 0.340 | 0.79 | 0.06 | 0.80 | 0.06 | 0.35 | -1.0 | 0.161 | 0.368 | 0.161 | 0.368 |
| Middle Temporal Gyrus; Temporoccipital Part L | 0.75 | 0.12 | 0.74 | 0.09 | 0.12 | 0.5 | 0.296 | 0.347 | 0.78 | 0.05 | 0.77 | 0.09 | 0.57 | 0.6 | 0.287 | 0.465 | 0.287 | 0.465 |
| Inferior Temporal Gyrus; Temporoccipital Part L | 0.69 | 0.19 | 0.70 | 0.10 | -0.10 | -0.5 | 0.319 | 0.367 | 0.77 | 0.06 | 0.78 | 0.05 | -0.71 | -0.4 | 0.328 | 0.480 | 0.328 | 0.480 |
| Occipital Fusiform Gyrus L | 0.74 | 0.12 | 0.72 | 0.12 | 0.13 | 0.6 | 0.269 | 0.340 | 0.82 | 0.07 | 0.83 | 0.06 | -0.73 | -0.3 | 0.386 | 0.491 | 0.386 | 0.491 |
| Supracalcarine Cortex L | 0.77 | 0.11 | 0.71 | 0.12 | 0.56 | 2.6 | 0.005 | 0.020 | 0.83 | 0.07 | 0.84 | 0.05 | -0.93 | -1.0 | 0.153 | 0.368 | 0.153 | 0.368 |
| Cuneal Cortex L | 0.74 | 0.15 | 0.71 | 0.13 | 0.21 | 1.0 | 0.171 | 0.248 | 0.82 | 0.07 | 0.82 | 0.05 | -0.73 | -0.4 | 0.352 | 0.480 | 0.352 | 0.480 |
| Lingual Gyrus L | 0.73 | 0.12 | 0.70 | 0.10 | 0.25 | 1.2 | 0.125 | 0.194 | 0.82 | 0.06 | 0.84 | 0.05 | -0.90 | -1.7 | 0.042 | 0.262 | 0.042 | 0.262 |
| Intracalcarine Cortex L | 0.74 | 0.15 | 0.70 | 0.13 | 0.27 | 1.2 | 0.108 | 0.182 | 0.83 | 0.08 | 0.85 | 0.06 | -0.65 | -0.8 | 0.202 | 0.393 | 0.202 | 0.393 |
| Lateral Occipital Cortex; Superior Division L | 0.77 | 0.03 | 0.74 | 0.04 | 1.00 | 4.6 | 0.000 | 0.000 | 0.74 | 0.06 | 0.76 | 0.06 | -0.53 | -1.1 | 0.137 | 0.368 | 0.137 | 0.368 |
| Lateral Occipital Cortex; Inferior Division L | 0.71 | 0.12 | 0.72 | 0.08 | -0.13 | -0.6 | 0.272 | 0.340 | 0.78 | 0.04 | 0.79 | 0.04 | -0.61 | -0.6 | 0.264 | 0.451 | 0.264 | 0.451 |
| Occipital Pole L | 0.76 | 0.08 | 0.72 | 0.09 | 0.49 | 2.2 | 0.014 | 0.038 | 0.83 | 0.08 | 0.84 | 0.07 | -0.29 | -1.1 | 0.139 | 0.368 | 0.139 | 0.368 |
| Frontal Medial Cortex | 0.76 | 0.08 | 0.74 | 0.08 | 0.26 | 1.2 | 0.112 | 0.186 | 0.77 | 0.11 | 0.77 | 0.07 | -0.13 | -0.1 | 0.459 | 0.491 | 0.459 | 0.491 |
| Cingulate Gyrus; Anterior Division | 0.80 | 0.09 | 0.72 | 0.13 | 0.73 | 3.3 | 0.001 | 0.005 | 0.76 | 0.08 | 0.76 | 0.05 | -0.13 | -0.1 | 0.461 | 0.491 | 0.461 | 0.491 |
| Subcallosal Cortex | 0.70 | 0.21 | 0.71 | 0.12 | -0.06 | -0.3 | 0.399 | 0.436 | 0.76 | 0.15 | 0.75 | 0.14 | -0.39 | 0.2 | 0.420 | 0.491 | 0.420 | 0.491 |
| Cingulate Gyrus; Posterior Division | 0.77 | 0.13 | 0.75 | 0.06 | 0.13 | 0.6 | 0.277 | 0.342 | 0.70 | 0.13 | 0.74 | 0.10 | 0.06 | -1.6 | 0.053 | 0.292 | 0.053 | 0.292 |
| Precuneus Cortex | 0.76 | 0.03 | 0.73 | 0.04 | 0.88 | 4.1 | 0.000 | 0.001 | 0.70 | 0.15 | 0.77 | 0.06 | 0.07 | -2.8 | 0.003 | 0.081 | 0.003 | 0.081 |
| Brain-Stem | 0.81 | 0.07 | 0.71 | 0.16 | 0.73 | 3.3 | 0.001 | 0.005 | 0.80 | 0.11 | 0.77 | 0.15 | 0.59 | 1.3 | 0.101 | 0.356 | 0.101 | 0.356 |
| Frontal Orbital Cortex R | 0.78 | 0.07 | 0.75 | 0.08 | 0.48 | 2.2 | 0.015 | 0.041 | 0.77 | 0.06 | 0.75 | 0.07 | 0.28 | 1.9 | 0.031 | 0.250 | 0.031 | 0.250 |
| Frontal Pole R | 0.74 | 0.02 | 0.73 | 0.03 | 0.29 | 1.4 | 0.089 | 0.155 | 0.77 | 0.06 | 0.74 | 0.08 | 0.28 | 1.7 | 0.042 | 0.262 | 0.042 | 0.262 |
| Frontal Operculum Cortex R | 0.78 | 0.15 | 0.75 | 0.09 | 0.24 | 1.1 | 0.140 | 0.212 | 0.78 | 0.06 | 0.79 | 0.08 | -0.02 | -0.2 | 0.439 | 0.491 | 0.439 | 0.491 |
| Superior Frontal Gyrus R | 0.77 | 0.13 | 0.74 | 0.10 | 0.34 | 1.6 | 0.059 | 0.111 | 0.77 | 0.07 | 0.74 | 0.08 | 0.15 | 2.2 | 0.015 | 0.177 | 0.015 | 0.177 |
| Middle Frontal Gyrus R | 0.77 | 0.05 | 0.74 | 0.05 | 0.70 | 3.2 | 0.001 | 0.006 | 0.77 | 0.06 | 0.76 | 0.07 | 0.17 | 0.7 | 0.240 | 0.432 | 0.240 | 0.432 |
| Inferior Frontal Gyrus; Pars Triangularis R | 0.82 | 0.07 | 0.76 | 0.06 | 0.89 | 4.1 | 0.000 | 0.001 | 0.77 | 0.06 | 0.76 | 0.05 | 0.00 | 0.8 | 0.227 | 0.417 | 0.227 | 0.417 |
| Inferior Frontal Gyrus; Pars Opercularis R | 0.81 | 0.05 | 0.76 | 0.07 | 0.83 | 3.8 | 0.000 | 0.001 | 0.76 | 0.09 | 0.74 | 0.11 | 0.30 | 1.0 | 0.149 | 0.368 | 0.149 | 0.368 |
| Paracingulate Gyrus R | 0.80 | 0.05 | 0.77 | 0.06 | 0.51 | 2.3 | 0.011 | 0.032 | 0.75 | 0.11 | 0.75 | 0.06 | 0.26 | -0.1 | 0.463 | 0.491 | 0.463 | 0.491 |

Supplementary Table 9. Mean, SD and effect sizes (Cohen's D) of regional betweenness centrality for younger and old adults in fPET and fMRI from the Harvard Oxford Atlas, and t-tests of age group differences.

|  | fPET |  |  |  |  |  |  |  |  | fMRI |  |  |  |  |  |  |  |  |
| --- | --- | --- | --- | --- | --- | --- | --- | --- | --- | --- | --- | --- | --- | --- | --- | --- | --- | --- |
|  | Younger |  |  | Older |  |  | Younger vs Older |  |  | Younger |  |  | Older |  |  | Younger vs Older |  |  |
|  | Mean | SD |  | Mean | SD | Cohen's D | t-value | p | p-FDR | Mean | SD |  | Mean | SD | Cohen's D | t-value | p | p-FDR |
| Frontal Orbital Cortex L | 0.011 | 0.007 | 0.008 | 0.005 | 0.006 | 0.60 | 2.8 | 0.003 | 0.022 | 0.012 | 0.009 | 0.011 | 0.008 | 0.011 | -0.13 | 0.5 | 0.315 | 0.433 |
| Frontal Pole L | 0.026 | 0.013 | 0.021 | 0.009 | 0.009 | 0.53 | 2.4 | 0.008 | 0.043 | 0.009 | 0.006 | 0.010 | 0.007 | 0.007 | 0.25 | -1.0 | 0.156 | 0.373 |
| Frontal Operculum Cortex L | 0.005 | 0.004 | 0.005 | 0.004 | 0.007 | 0.03 | 0.379 | 0.709 | 0.423 | 0.007 | 0.006 | 0.007 | 0.005 | 0.005 | 0.23 | -0.1 | 0.480 | 0.496 |
| Superior Frontal Gyrus L | 0.013 | 0.008 | 0.008 | 0.006 | 0.006 | 0.63 | 2.9 | 0.002 | 0.022 | 0.008 | 0.006 | 0.009 | 0.007 | 0.007 | -0.05 | -0.7 | 0.229 | 0.389 |
| Middle Frontal Gyrus L | 0.015 | 0.006 | 0.012 | 0.007 | 0.007 | 0.39 | 1.8 | 0.039 | 0.115 | 0.005 | 0.004 | 0.006 | 0.004 | 0.004 | 0.17 | -1.5 | 0.071 | 0.354 |
| Inferior Frontal Gyrus; Pars Triangularis L | 0.008 | 0.005 | 0.008 | 0.004 | 0.004 | 0.01 | 0.0 | 0.482 | 0.490 | 0.006 | 0.005 | 0.006 | 0.004 | 0.004 | 0.17 | 0.6 | 0.291 | 0.417 |
| Inferior Frontal Gyrus; pars opercularis L | 0.007 | 0.005 | 0.006 | 0.005 | 0.005 | 0.15 | 0.7 | 0.243 | 0.362 | 0.007 | 0.006 | 0.006 | 0.005 | 0.005 | -0.21 | 0.7 | 0.238 | 0.389 |
| Paracalcarine Gyrus L | 0.010 | 0.007 | 0.006 | 0.004 | 0.004 | 0.60 | 2.8 | 0.003 | 0.022 | 0.007 | 0.005 | 0.008 | 0.005 | 0.005 | 0.12 | -1.3 | 0.103 | 0.360 |
| Insular Cortex L | 0.004 | 0.003 | 0.004 | 0.003 | 0.005 | 0.02 | 0.410 | 0.439 | 0.022 | 0.010 | 0.006 | 0.010 | 0.007 | 0.010 | -0.2 | 0.414 | 0.462 |  |
| Amygdala l | 0.004 | 0.004 | 0.003 | 0.002 | 0.002 | 0.31 | 1.4 | 0.077 | 0.190 | 0.005 | 0.006 | 0.003 | 0.005 | 0.005 | -0.10 | 1.9 | 0.030 | 0.255 |
| Juxtastriatal Lobule Cortex - L | 0.005 | 0.006 | 0.005 | 0.003 | 0.003 | 0.13 | 0.6 | 0.274 | 0.392 | 0.005 | 0.004 | 0.007 | 0.004 | 0.004 | -0.06 | -1.9 | 0.031 | 0.255 |
| Precentral Gyrus L | 0.011 | 0.006 | 0.010 | 0.008 | 0.006 | 0.06 | 0.3 | 0.398 | 0.435 | 0.010 | 0.007 | 0.012 | 0.007 | 0.007 | -0.07 | -1.5 | 0.069 | 0.354 |
| PostCG I Postcentral Gyrus L | 0.009 | 0.006 | 0.012 | 0.008 | 0.008 | -0.39 | -1.8 | 0.037 | 0.115 | 0.009 | 0.008 | 0.010 | 0.005 | 0.005 | -0.22 | -0.6 | 0.260 | 0.394 |
| Central Opercular Cortex L | 0.003 | 0.003 | 0.006 | 0.005 | 0.005 | -0.63 | -2.9 | 0.002 | 0.022 | 0.008 | 0.005 | 0.007 | 0.004 | 0.004 | -0.08 | 1.1 | 0.136 | 0.360 |
| Superior Parietal Lobule L | 0.007 | 0.005 | 0.006 | 0.004 | 0.004 | 0.25 | 1.1 | 0.129 | 0.261 | 0.007 | 0.005 | 0.010 | 0.005 | 0.010 | -0.10 | -2.7 | 0.004 | 0.136 |
| Supramarginal Gyrus; Anterior Division L | 0.005 | 0.004 | 0.005 | 0.004 | 0.010 | 0.5 | 0.318 | 0.409 | 0.005 | 0.005 | 0.005 | 0.004 | 0.004 | 0.004 | -0.37 | -0.5 | 0.301 | 0.419 |
| Supramarginal Gyrus; Posterior Division L | 0.008 | 0.004 | 0.008 | 0.004 | 0.007 | -0.3 | 0.368 | 0.422 | 0.009 | 0.007 | 0.008 | 0.004 | 0.004 | 0.004 | -0.18 | 0.3 | 0.394 | 0.460 |
| Angular Gyrus L | 0.005 | 0.004 | 0.006 | 0.004 | 0.004 | -0.24 | -1.1 | 0.138 | 0.263 | 0.011 | 0.009 | 0.005 | 0.004 | 0.004 | -0.20 | 3.8 | 0.000 | 0.015 |
| Parietal Operculum Cortex L | 0.005 | 0.004 | 0.006 | 0.005 | 0.005 | -0.29 | -1.3 | 0.091 | 0.201 | 0.009 | 0.006 | 0.009 | 0.006 | 0.006 | -0.24 | 0.0 | 0.498 | 0.498 |
| Parahippocampal Gyrus; Anterior Division L | 0.003 | 0.003 | 0.005 | 0.003 | 0.003 | -0.46 | -2.1 | 0.019 | 0.076 | 0.007 | 0.010 | 0.005 | 0.005 | 0.005 | -0.14 | 0.9 | 0.198 | 0.373 |
| Parahippocampal Gyrus; Posterior Division L | 0.002 | 0.002 | 0.003 | 0.003 | 0.003 | -0.67 | -3.1 | 0.001 | 0.022 | 0.005 | 0.004 | 0.005 | 0.005 | 0.005 | -0.14 | 0.1 | 0.477 | 0.496 |
| Hippocampus l | 0.005 | 0.005 | 0.004 | 0.003 | 0.004 | 0.24 | 1.1 | 0.131 | 0.261 | 0.008 | 0.005 | 0.006 | 0.006 | 0.006 | -0.23 | 1.1 | 0.130 | 0.360 |
| Thalamus L | 0.002 | 0.002 | 0.004 | 0.003 | 0.003 | -0.62 | -2.9 | 0.002 | 0.022 | 0.007 | 0.006 | 0.006 | 0.006 | 0.006 | -0.02 | 1.1 | 0.131 | 0.360 |
| Caudate L | 0.004 | 0.004 | 0.007 | 0.004 | 0.004 | -0.61 | -2.8 | 0.003 | 0.022 | 0.005 | 0.004 | 0.006 | 0.008 | 0.008 | -0.03 | -0.7 | 0.240 | 0.389 |
| Putamen L | 0.009 | 0.009 | 0.008 | 0.006 | 0.006 | 0.15 | 0.7 | 0.249 | 0.366 | 0.005 | 0.004 | 0.006 | 0.009 | 0.009 | -0.02 | -0.3 | 0.365 | 0.457 |
| Pallidum L | 0.003 | 0.002 | 0.005 | 0.003 | 0.003 | -0.79 | -3.6 | 0.000 | 0.008 | 0.004 | 0.004 | 0.005 | 0.006 | 0.004 | -0.13 | 1.0 | 0.100 | 0.360 |
| Accumbens L | 0.002 | 0.002 | 0.003 | 0.003 | 0.003 | -0.39 | -1.8 | 0.039 | 0.115 | 0.005 | 0.007 | 0.004 | 0.006 | 0.005 | 1.0 | 0.168 | 0.373 |  |
| Temporal Pole L | 0.006 | 0.005 | 0.005 | 0.003 | 0.003 | 0.38 | 1.8 | 0.041 | 0.116 | 0.011 | 0.007 | 0.013 | 0.010 | 0.010 | -0.35 | -1.3 | 0.095 | 0.360 |
| Planum Polare L | 0.002 | 0.002 | 0.003 | 0.002 | 0.002 | -0.63 | -2.9 | 0.002 | 0.022 | 0.006 | 0.004 | 0.005 | 0.004 | 0.004 | -0.60 | 0.8 | 0.213 | 0.382 |
| Superior Temporal Gyrus; Anterior Division L | 0.002 | 0.002 | 0.003 | 0.002 | 0.002 | -0.43 | -2.0 | 0.026 | 0.095 | 0.009 | 0.008 | 0.008 | 0.008 | 0.008 | -1.20 | 0.2 | 0.437 | 0.468 |
| Superior Temporal Gyrus; Posterior Division L | 0.004 | 0.004 | 0.005 | 0.004 | 0.004 | -0.42 | -1.9 | 0.028 | 0.098 | 0.009 | 0.006 | 0.009 | 0.006 | 0.006 | 0.29 | 0.0 | 0.484 | 0.496 |
| Middle Temporal Gyrus; Anterior Division L | 0.006 | 0.006 | 0.004 | 0.003 | 0.003 | 0.40 | 1.9 | 0.033 | 0.105 | 0.007 | 0.005 | 0.006 | 0.005 | 0.005 | 0.41 | 0.6 | 0.271 | 0.405 |
| Middle Temporal Gyrus; Posterior Division L | 0.014 | 0.010 | 0.007 | 0.004 | 0.004 | 0.89 | 4.1 | 0.000 | 0.004 | 0.008 | 0.006 | 0.008 | 0.005 | 0.005 | 0.39 | -0.1 | 0.442 | 0.468 |
| Inferior Temporal Gyrus; Anterior Division L | 0.004 | 0.003 | 0.003 | 0.002 | 0.010 | 0.5 | 0.324 | 0.409 | 0.005 | 0.004 | 0.005 | 0.004 | 0.004 | 0.004 | 0.37 | 0.2 | 0.437 | 0.468 |
| Inferior Temporal Gyrus; Posterior Division L | 0.008 | 0.006 | 0.006 | 0.004 | 0.004 | 0.51 | 2.4 | 0.011 | 0.051 | 0.010 | 0.010 | 0.005 | 0.004 | 0.004 | -0.03 | 3.1 | 0.001 | 0.074 |
| Planum Temporale L | 0.010 | 0.007 | 0.007 | 0.004 | 0.004 | 0.44 | 2.0 | 0.023 | 0.086 | 0.009 | 0.006 | 0.011 | 0.006 | 0.006 | -0.03 | -1.8 | 0.041 | 0.262 |
| Heschl's Gyrus L | 0.006 | 0.005 | 0.007 | 0.004 | 0.004 | -0.23 | -1.1 | 0.141 | 0.263 | 0.008 | 0.007 | 0.009 | 0.006 | 0.006 | 0.47 | -0.9 | 0.191 | 0.373 |
| Temporal Fusiform Cortex; Anterior Division L | 0.003 | 0.003 | 0.003 | 0.003 | 0.003 | -0.24 | -1.1 | 0.139 | 0.263 | 0.005 | 0.006 | 0.005 | 0.005 | 0.005 | 0.15 | 0.3 | 0.367 | 0.457 |
| Temporal Fusiform Cortex; Posterior Division L | 0.004 | 0.003 | 0.005 | 0.004 | 0.004 | -0.27 | -1.3 | 0.106 | 0.230 | 0.012 | 0.010 | 0.011 | 0.007 | 0.007 | 0.14 | 0.2 | 0.402 | 0.460 |
| Temporal Occipital Fusiform Cortex L | 0.003 | 0.003 | 0.005 | 0.004 | 0.004 | -0.63 | -2.9 | 0.002 | 0.022 | 0.011 | 0.009 | 0.008 | 0.006 | 0.006 | 0.16 | 2.1 | 0.019 | 0.220 |
| Middle Temporal Gyrus; Temporoccipital Part L | 0.006 | 0.004 | 0.006 | 0.004 | 0.004 | 0.18 | 0.8 | 0.202 | 0.330 | 0.009 | 0.006 | 0.007 | 0.004 | 0.004 | 0.23 | 2.4 | 0.009 | 0.136 |
| Inferior Temporal Gyrus; Temporoccipital Part L | 0.004 | 0.003 | 0.005 | 0.004 | 0.004 | -0.26 | -1.2 | 0.113 | 0.240 | 0.009 | 0.006 | 0.007 | 0.004 | 0.004 | 0.21 | 2.5 | 0.008 | 0.136 |
| Occipital Fusiform Gyrus L | 0.006 | 0.005 | 0.007 | 0.005 | 0.005 | -0.14 | -0.7 | 0.253 | 0.367 | 0.008 | 0.010 | 0.006 | 0.005 | 0.005 | -0.02 | 1.2 | 0.121 | 0.360 |
| Supracalcarine Cortex L | 0.004 | 0.003 | 0.005 | 0.003 | 0.003 | -0.46 | -2.1 | 0.019 | 0.076 | 0.004 | 0.004 | 0.004 | 0.003 | 0.003 | 0.20 | 0.4 | 0.343 | 0.449 |
| Cuneal Cortex L | 0.004 | 0.004 | 0.005 | 0.003 | 0.003 | -0.08 | -0.4 | 0.350 | 0.422 | 0.006 | 0.005 | 0.007 | 0.005 | 0.005 | 0.22 | -0.4 | 0.330 | 0.448 |
| Lingual Gyrus L | 0.006 | 0.005 | 0.007 | 0.005 | 0.005 | -0.11 | -0.5 | 0.301 | 0.404 | 0.006 | 0.007 | 0.005 | 0.005 | 0.005 | -0.25 | 1.0 | 0.172 | 0.373 |
| Intracalcarine Cortex L | 0.006 | 0.006 | 0.006 | 0.004 | 0.004 | 0.13 | 0.6 | 0.278 | 0.392 | 0.004 | 0.004 | 0.004 | 0.004 | 0.004 | 0.32 | -0.3 | 0.383 | 0.460 |
| Lateral Occipital Cortex; Superior Division L | 0.017 | 0.009 | 0.015 | 0.008 | 0.008 | 0.29 | 1.3 | 0.090 | 0.201 | 0.014 | 0.008 | 0.012 | 0.008 | 0.008 | 0.33 | 0.9 | 0.176 | 0.373 |
| Lateral Occipital Cortex; Inferior Division L | 0.008 | 0.006 | 0.009 | 0.006 | 0.006 | -0.10 | -0.5 | 0.317 | 0.409 | 0.011 | 0.007 | 0.009 | 0.006 | 0.006 | 0.24 | 1.3 | 0.104 | 0.360 |
| Occipital Pole L | 0.011 | 0.009 | 0.012 | 0.007 | 0.007 | -0.09 | -0.4 | 0.346 | 0.421 | 0.004 | 0.004 | 0.005 | 0.005 | 0.005 | 0.09 | -0.2 | 0.403 | 0.460 |
| Frontal Medial Cortex | 0.012 | 0.008 | 0.010 | 0.006 | 0.006 | 0.31 | 1.4 | 0.080 | 0.192 | 0.007 | 0.005 | 0.007 | 0.007 | 0.007 | 0.10 | -0.6 | 0.282 | 0.410 |
| Cingulate Gyrus; Anterior Division | 0.009 | 0.006 | 0.008 | 0.006 | 0.006 | 0.23 | 1.1 | 0.144 | 0.264 | 0.008 | 0.006 | 0.010 | 0.006 | 0.006 | 0.12 | -1.8 | 0.038 | 0.262 |
| Subcallosal Cortex | 0.005 | 0.005 | 0.004 | 0.004 | 0.004 | 0.32 | 1.5 | 0.073 | 0.185 | 0.007 | 0.006 | 0.006 | 0.007 | 0.007 | 0.13 | 0.8 | 0.225 | 0.389 |
| Cingulate Gyrus; Posterior Division | 0.016 | 0.010 | 0.015 | 0.007 | 0.007 | 0.12 | 0.6 | 0.292 | 0.400 | 0.008 | 0.009 | 0.009 | 0.008 | 0.008 | 0.13 | -0.3 | 0.397 | 0.460 |
| Precuneus Cortex | 0.021 | 0.010 | 0.022 | 0.009 | 0.009 | -0.12 | -0.6 | 0.291 | 0.400 | 0.010 | 0.008 | 0.008 | 0.005 | 0.005 | -0.14 | 0.9 | 0.190 | 0.373 |
| Brain-Stem | 0.010 | 0.008 | 0.006 | 0.005 | 0.005 | 0.67 | 3.1 | 0.001 | 0.022 | 0.003 | 0.005 | 0.004 | 0.006 | 0.006 | 0.00 | -0.8 | 0.216 | 0.382 |
| Frontal Orbital Cortex R | 0.011 | 0.006 | 0.009 | 0.005 | 0.005 | 0.37 | 1.7 | 0.047 | 0.129 | 0.009 | 0.008 | 0.011 | 0.010 | 0.010 | 0.00 | -1.1 | 0.147 | 0.370 |
| Frontal Pole R | 0.030 | 0.012 | 0.022 | 0.007 | 0.007 | 0.86 | 4.0 | 0.000 | 0.004 | 0.008 | 0.004 | 0.009 | 0.005 | 0.005 | 0.18 | -0.4 | 0.334 | 0.448 |
| Frontal Operculum Cortex R | 0.006 | 0.005 | 0.007 | 0.005 | 0.005 | -0.29 | -1.3 | 0.091 | 0.201 | 0.008 | 0.006 | 0.007 | 0.006 | 0.006 | -0.03 | 0.7 | 0.249 | 0.389 |
| Superior Frontal Gyrus R | 0.011 | 0.006 | 0.008 | 0.005 | 0.005 | 0.47 | 2.2 | 0.016 | 0.073 | 0.008 | 0.005 | 0.009 | 0.004 | 0.004 | -0.03 | -0.9 | 0.184 | 0.373 |
| Middle Frontal Gyrus R | 0.018 | 0.007 | 0.014 | 0.007 | 0.007 | 0.55 | 2.6 | 0.006 | 0.037 | 0.005 | 0.004 | 0.006 | 0.006 | 0.006 | 0.11 | -0.8 | 0.199 | 0.373 |
| Inferior Frontal Gyrus; Pars Triangularis R | 0.007 | 0.005 | 0.009 | 0.005 | 0.005 | -0.37 |  |  |  |  |  |  |  |  |  |  |  |  |

Supplementary Table 10. Mean, SD and effect sizes (Cohen's D) of regional betweenness centrality for younger and older adults in fPET and fMRI from the Harvard Oxford Atlas, and t-tests of age group differences.

|  | fPET |  |  |  |  |  |  |  |  | fMRI |  |  |  |  |  |  |  |  |
| --- | --- | --- | --- | --- | --- | --- | --- | --- | --- | --- | --- | --- | --- | --- | --- | --- | --- | --- |
|  | Younger |  |  | Older |  |  | Younger vs Older |  |  | Younger |  |  | Older |  |  | Younger vs Older |  |  |
|  | Mean | SD |  | Mean | SD | Cohen's D | t-value | p | p-FDR | Mean | SD |  | Mean | SD | Cohen's D | t-value | p | p-FDR |
| Frontal Orbital Cortex L | 44.28 | 13.57 | 37.43 | 13.00 | 0.52 | 2.4 | 0.010 | 0.032 |  | 39.00 | 12.37 | 32.35 | 15.24 | 0.10 | 2.2 | 0.015 | 0.057 |  |
| Frontal Pole L | 68.63 | 8.77 | 61.76 | 10.48 | 0.71 | 3.3 | 0.001 | 0.006 |  | 35.20 | 13.42 | 32.89 | 12.98 | 0.11 | 0.8 | 0.210 | 0.329 |  |
| Frontal Operculum Cortex L | 27.05 | 12.22 | 28.41 | 12.01 | -0.11 | -0.5 | 0.302 | 0.410 |  | 30.63 | 11.84 | 28.87 | 11.23 | -0.22 | 0.7 | 0.241 | 0.347 |  |
| Superior Frontal Gyrus L | 50.20 | 15.53 | 39.35 | 16.10 | 0.69 | 3.2 | 0.001 | 0.006 |  | 31.80 | 13.46 | 31.54 | 12.12 | -0.01 | 0.1 | 0.463 | 0.487 |  |
| Middle Frontal Gyrus L | 55.45 | 8.63 | 47.98 | 12.22 | 0.70 | 3.2 | 0.001 | 0.006 |  | 26.05 | 12.63 | 27.83 | 12.30 | -0.01 | -0.7 | 0.256 | 0.348 |  |
| Inferior Frontal Gyrus; Pars Triangularis L | 43.10 | 13.10 | 39.24 | 11.12 | 0.32 | 1.5 | 0.072 | 0.147 |  | 28.35 | 12.86 | 26.41 | 9.98 | -0.16 | 0.8 | 0.217 | 0.333 |  |
| Inferior Frontal Gyrus; pars opercularis L | 37.88 | 13.99 | 31.52 | 12.82 | 0.48 | 2.2 | 0.015 | 0.049 |  | 29.50 | 10.70 | 27.67 | 11.47 | -0.32 | 0.8 | 0.225 | 0.333 |  |
| Paracingulate Gyrus L | 45.48 | 10.48 | 33.43 | 12.43 | 1.04 | 4.8 | 0.000 | 0.000 |  | 28.00 | 13.22 | 30.11 | 12.34 | -0.30 | -0.8 | 0.223 | 0.333 |  |
| Insular Cortex L | 20.28 | 10.54 | 21.15 | 9.65 | -0.09 | -0.4 | 0.344 | 0.414 |  | 36.00 | 12.63 | 36.54 | 10.11 | 0.12 | -0.2 | 0.413 | 0.460 |  |
| Amygdala l | 15.00 | 11.79 | 16.72 | 8.76 | -0.17 | -0.8 | 0.221 | 0.349 |  | 25.60 | 12.81 | 17.50 | 12.78 | 0.15 | 2.9 | 0.002 | 0.014 |  |
| Juxtastriatal Lobule Cortex - L | 26.85 | 13.62 | 26.93 | 12.59 | -0.01 | 0.0 | 0.488 | 0.493 |  | 30.30 | 12.68 | 36.20 | 10.10 | 0.17 | -2.4 | 0.009 | 0.043 |  |
| Precentral Gyrus L | 46.28 | 16.53 | 40.91 | 15.80 | 0.33 | 1.5 | 0.064 | 0.142 |  | 38.83 | 16.48 | 45.52 | 9.99 | -0.27 | -2.3 | 0.012 | 0.049 |  |
| PostCG l Postcentral Gyrus L | 43.43 | 15.64 | 44.35 | 15.80 | -0.06 | -0.3 | 0.393 | 0.439 |  | 35.50 | 14.58 | 44.15 | 9.04 | -0.05 | -3.4 | 0.001 | 0.005 |  |
| Central Opercular Cortex L | 20.93 | 12.94 | 28.76 | 12.83 | -0.61 | -2.8 | 0.003 | 0.014 |  | 38.03 | 12.49 | 37.33 | 11.97 | -0.04 | 0.3 | 0.396 | 0.451 |  |
| Superior Parietal Lobule L | 37.78 | 16.33 | 34.17 | 12.87 | 0.25 | 1.1 | 0.128 | 0.219 |  | 31.98 | 12.63 | 41.28 | 9.63 | 0.41 | -3.9 | 0.000 | 0.000 |  |
| Supramarginal Gyrus; Anterior Division L | 34.88 | 13.80 | 28.15 | 12.07 | 0.52 | 2.4 | 0.009 | 0.031 |  | 29.85 | 11.58 | 31.80 | 9.80 | -0.41 | -0.8 | 0.200 | 0.320 |  |
| Supramarginal Gyrus; Posterior Division L | 41.25 | 11.81 | 37.57 | 11.35 | 0.32 | 1.5 | 0.072 | 0.147 |  | 34.30 | 13.30 | 35.72 | 12.72 | -0.39 | -0.5 | 0.308 | 0.388 |  |
| Angular Gyrus L | 33.68 | 12.86 | 34.15 | 11.49 | -0.04 | -0.2 | 0.428 | 0.468 |  | 35.60 | 12.65 | 25.76 | 12.75 | -0.32 | 3.6 | 0.000 | 0.004 |  |
| Parietal Operculum Cortex L | 28.33 | 14.59 | 29.09 | 10.75 | -0.06 | -0.3 | 0.391 | 0.439 |  | 40.60 | 10.82 | 40.80 | 13.14 | -0.14 | -0.1 | 0.469 | 0.487 |  |
| Parahippocampal Gyrus; Anterior Division L | 14.45 | 9.49 | 18.57 | 8.42 | -0.46 | -2.1 | 0.018 | 0.054 |  | 27.13 | 14.33 | 25.93 | 12.94 | -0.19 | 0.4 | 0.343 | 0.409 |  |
| Parahippocampal Gyrus; Posterior Division L | 11.18 | 6.50 | 17.00 | 9.66 | -0.70 | -3.2 | 0.001 | 0.006 |  | 23.63 | 12.72 | 23.89 | 13.77 | 0.24 | -0.1 | 0.463 | 0.487 |  |
| Hippocampus l | 17.23 | 12.68 | 17.91 | 10.84 | -0.06 | -0.3 | 0.393 | 0.439 |  | 31.38 | 13.28 | 24.59 | 15.22 | -0.58 | 2.2 | 0.016 | 0.057 |  |
| Thalamus L | 14.80 | 9.45 | 23.02 | 11.04 | -0.80 | -3.7 | 0.000 | 0.003 |  | 27.03 | 15.29 | 24.35 | 15.43 | -0.57 | 0.8 | 0.211 | 0.329 |  |
| Caudate L | 19.73 | 11.53 | 29.72 | 13.14 | -0.80 | -3.7 | 0.000 | 0.003 |  | 21.63 | 12.89 | 21.76 | 12.51 | -0.11 | 0.0 | 0.480 | 0.488 |  |
| Putamen L | 29.43 | 14.48 | 34.15 | 14.50 | -0.33 | -1.5 | 0.068 | 0.146 |  | 23.80 | 13.47 | 21.22 | 12.77 | 0.06 | 0.9 | 0.182 | 0.307 |  |
| Pallidum L | 12.43 | 7.45 | 21.11 | 10.15 | -0.97 | -4.5 | 0.000 | 0.000 |  | 22.03 | 14.52 | 20.20 | 14.79 | 0.07 | 0.6 | 0.283 | 0.366 |  |
| Accumbens L | 13.20 | 9.02 | 21.65 | 9.86 | -0.89 | -4.1 | 0.000 | 0.001 |  | 19.25 | 15.16 | 15.39 | 14.51 | 0.81 | 1.2 | 0.116 | 0.223 |  |
| Temporal Pole L | 24.13 | 13.39 | 21.85 | 10.39 | 0.19 | 0.9 | 0.189 | 0.303 |  | 40.80 | 10.65 | 36.78 | 11.73 | 0.00 | 1.7 | 0.051 | 0.132 |  |
| Planum Polare L | 12.65 | 8.08 | 19.00 | 9.02 | -0.74 | -3.4 | 0.000 | 0.005 |  | 32.50 | 12.01 | 29.70 | 14.23 | 0.00 | 1.0 | 0.165 | 0.282 |  |
| Superior Temporal Gyrus; Anterior Division L | 15.23 | 10.33 | 18.52 | 8.33 | -0.35 | -1.6 | 0.053 | 0.122 |  | 34.90 | 13.34 | 34.50 | 15.01 | 0.18 | 0.1 | 0.449 | 0.485 |  |
| Superior Temporal Gyrus; Posterior Division L | 23.38 | 13.93 | 28.72 | 10.72 | -0.43 | -2.0 | 0.024 | 0.067 |  | 36.83 | 13.21 | 37.22 | 12.49 | 0.01 | -0.1 | 0.444 | 0.485 |  |
| Middle Temporal Gyrus; Anterior Division L | 22.30 | 12.83 | 20.93 | 9.05 | 0.12 | 0.6 | 0.283 | 0.395 |  | 30.75 | 13.69 | 29.41 | 12.49 | 0.01 | 0.5 | 0.319 | 0.397 |  |
| Middle Temporal Gyrus; Posterior Division L | 46.13 | 13.93 | 35.96 | 14.04 | 0.73 | 3.4 | 0.001 | 0.005 |  | 35.33 | 14.16 | 30.78 | 10.71 | 0.24 | 1.7 | 0.047 | 0.125 |  |
| Inferior Temporal Gyrus; Anterior Division L | 19.73 | 11.35 | 18.76 | 8.54 | 0.10 | 0.4 | 0.327 | 0.414 |  | 29.90 | 11.71 | 23.85 | 13.06 | 0.24 | 2.2 | 0.014 | 0.053 |  |
| Inferior Temporal Gyrus; Posterior Division L | 36.58 | 12.81 | 27.65 | 12.88 | 0.69 | 3.2 | 0.001 | 0.006 |  | 36.53 | 13.57 | 25.35 | 12.49 | 0.25 | 4.0 | 0.000 | 0.003 |  |
| Planum Temporale L | 43.05 | 12.38 | 37.89 | 10.63 | 0.45 | 2.1 | 0.020 | 0.058 |  | 39.13 | 12.98 | 42.93 | 12.78 | -0.15 | -1.4 | 0.087 | 0.181 |  |
| Heschl's Gyrus L | 30.60 | 12.90 | 33.87 | 11.17 | -0.27 | -1.3 | 0.106 | 0.189 |  | 32.70 | 12.43 | 34.72 | 14.33 | -0.08 | -0.7 | 0.245 | 0.347 |  |
| Temporal Fusiform Cortex; Anterior Division L | 13.83 | 9.57 | 18.54 | 10.62 | -0.47 | -2.2 | 0.017 | 0.054 |  | 23.75 | 16.07 | 23.78 | 12.75 | -0.06 | 0.0 | 0.496 | 0.496 |  |
| Temporal Fusiform Cortex; Posterior Division L | 19.85 | 9.37 | 22.67 | 9.95 | -0.29 | -1.3 | 0.091 | 0.178 |  | 38.35 | 15.11 | 35.98 | 9.92 | -0.28 | 0.9 | 0.193 | 0.320 |  |
| Temporal Occipital Fusiform Cortex L | 19.50 | 10.56 | 25.41 | 9.03 | -0.60 | -2.8 | 0.003 | 0.014 |  | 36.25 | 12.40 | 34.61 | 10.76 | 0.21 | 0.7 | 0.256 | 0.348 |  |
| Middle Temporal Gyrus; Temporoccipital Part L | 30.68 | 12.73 | 30.72 | 11.99 | 0.00 | 0.0 | 0.494 | 0.494 |  | 38.03 | 13.53 | 32.63 | 10.15 | 0.23 | 2.1 | 0.019 | 0.062 |  |
| Inferior Temporal Gyrus; Temporoccipital Part L | 24.18 | 12.04 | 25.67 | 9.34 | -0.14 | -0.6 | 0.259 | 0.376 |  | 36.08 | 12.23 | 30.17 | 10.60 | -0.28 | 2.4 | 0.009 | 0.043 |  |
| Occipital Fusiform Gyrus L | 30.65 | 16.60 | 34.70 | 13.31 | -0.27 | -1.3 | 0.107 | 0.189 |  | 29.58 | 11.94 | 33.74 | 10.15 | 0.17 | -1.7 | 0.042 | 0.122 |  |
| Supracalcarine Cortex L | 25.45 | 9.50 | 27.87 | 10.66 | -0.24 | -1.1 | 0.136 | 0.230 |  | 24.60 | 11.43 | 33.28 | 9.68 | 0.18 | -3.8 | 0.000 | 0.003 |  |
| Cuneal Cortex L | 25.18 | 13.49 | 26.83 | 10.96 | -0.14 | -0.6 | 0.267 | 0.377 |  | 27.80 | 11.57 | 37.28 | 11.33 | 0.03 | -3.8 | 0.000 | 0.003 |  |
| Lingual Gyrus L | 27.95 | 12.90 | 29.93 | 13.72 | -0.15 | -0.7 | 0.247 | 0.368 |  | 28.68 | 11.01 | 34.26 | 9.29 | -0.01 | -2.6 | 0.006 | 0.033 |  |
| Intracalcarine Cortex L | 31.48 | 15.30 | 30.78 | 13.16 | 0.05 | 0.2 | 0.411 | 0.454 |  | 23.80 | 11.74 | 32.54 | 9.79 | -0.01 | -3.8 | 0.000 | 0.003 |  |
| Lateral Occipital Cortex; Superior Division L | 56.88 | 10.17 | 52.76 | 12.34 | 0.36 | 1.7 | 0.049 | 0.121 |  | 37.83 | 11.14 | 39.37 | 10.62 | 0.13 | -0.7 | 0.256 | 0.348 |  |
| Lateral Occipital Cortex; Inferior Division L | 33.18 | 17.22 | 38.28 | 14.40 | -0.32 | -1.5 | 0.069 | 0.146 |  | 38.20 | 11.90 | 38.30 | 11.76 | -0.03 | 0.0 | 0.484 | 0.488 |  |
| Occipital Pole L | 41.60 | 16.29 | 44.07 | 16.33 | -0.15 | -0.7 | 0.243 | 0.368 |  | 23.78 | 9.30 | 30.98 | 9.47 | -0.03 | -3.5 | 0.000 | 0.004 |  |
| Frontal Medial Cortex | 44.60 | 15.99 | 39.63 | 12.70 | 0.35 | 1.6 | 0.056 | 0.126 |  | 29.23 | 12.48 | 27.80 | 13.34 | 0.03 | 0.5 | 0.307 | 0.388 |  |
| Cingulate Gyrus; Anterior Division | 42.80 | 14.40 | 34.30 | 15.65 | 0.56 | 2.6 | 0.005 | 0.021 |  | 28.88 | 12.52 | 34.39 | 11.15 | 0.67 | -2.2 | 0.017 | 0.059 |  |
| Subcallosal Cortex | 27.50 | 15.87 | 24.35 | 11.13 | 0.23 | 1.1 | 0.142 | 0.236 |  | 26.78 | 16.09 | 24.09 | 16.75 | 1.19 | 0.8 | 0.226 | 0.333 |  |
| Cingulate Gyrus; Posterior Division | 54.88 | 12.82 | 54.76 | 10.69 | 0.01 | 0.0 | 0.482 | 0.491 |  | 25.50 | 14.36 | 29.41 | 11.95 | -0.38 | -1.4 | 0.086 | 0.181 |  |
| Precuneous Cortex | 62.65 | 12.27 | 63.89 | 8.90 | -0.12 | -0.5 | 0.295 | 0.406 |  | 25.75 | 12.31 | 34.76 | 11.77 | -0.19 | -3.5 | 0.000 | 0.004 |  |
| Brain-Stem | 41.30 | 15.99 | 32.13 | 15.03 | 0.59 | 2.7 | 0.004 | 0.015 |  | 16.10 | 13.84 | 19.39 | 14.47 | -0.21 | -1.1 | 0.143 | 0.257 |  |
| Frontal Orbital Cortex R | 46.63 | 13.98 | 41.48 | 11.74 | 0.40 | 1.9 | 0.033 | 0.091 |  | 35.90 | 12.33 | 31.59 | 12.79 | 0.07 | 1.6 | 0.058 | 0 |  |

Supplementary Table 11. Regression analyses predicting cognitive performance from fPET and fMRI whole brain graph metrics from the Harvard Oxford Atlas. The ANOVA results for the overall models are shown in the top three rows, and the standardized beta weights for each predictor in subsequent rows. Significant beta weights are indicated at \*p < .05; and \*\*p < .001.

|  | HVLT:<br>Delayed<br>Recall | HVLT:<br>Discrim.<br>Index | Digit<br>Span:<br>Forward | Digit<br>Span:<br>Back | Category<br>Switch:<br>% Trials | Category<br>Switch:<br>RT | Dig Sub:<br>Number<br>Correct | Dig Sub:<br>Sec Per<br>Correct | Stop<br>Signal:<br>Pro Reac | Stop<br>Signal:<br>RT |
| --- | --- | --- | --- | --- | --- | --- | --- | --- | --- | --- |
| ANOVA F | 1.2 | 3.1 | 0.3 | 1.6 | 2.6 | 1.8 | 2.5 | 0.6 | 1.5 | 1.9 |
| ANOVA p | 0.341 | 0.010 | 0.936 | 0.167 | 0.023 | 0.106 | 0.026 | 0.699 | 0.181 | 0.089 |
| Variance Explained | 8% | 19% | 2% | 11% | 17% | 12% | 17% | 5% | 10% | 13% |
| fPET: Global Efficiency | 0.16 | 0.73* | 0.12 | 0.12 | 0.26 | -0.09 | -0.07 | -0.39 | -0.29 | 0.30 |
| fPET: Local Efficiency | 0.35 | 0.83** | 0.00 | -0.08 | 0.41 | 0.02 | 0.19 | -0.29 | 0.00 | 0.47* |
| fPET: Betweenness Centrality | 0.01 | -0.14 | -0.04 | 0.05 | -0.09 | 0.17 | 0.26 | 0.17 | 0.32* | 0.05 |
| fMRI: Global Efficiency | -0.25 | -0.16 | 0.09 | 0.17 | -0.30 | 0.07 | -0.18 | -0.13 | -0.01 | -0.16 |
| fMRI: Local Efficiency | -0.35 | -0.05 | -0.10 | -0.24 | -0.39* | -0.40 | -0.37 | -0.06 | -0.07 | -0.28 |
| fMRI: Betweenness Centrality | 0.11 | 0.03 | 0.12 | 0.31 | 0.33* | 0.35* | 0.34* | 0.07 | 0.04 | -0.12 |

Discrim = discrimination; Dig Sub = Digit Substitution; Pro React = probability of reacting.

Supplementary Table 12. Regression analyses predicting cognitive performance from fPET and fMRI whole brain graph metrics from the Harvard Oxford Atlas and age group. The ANOVA results for the overall models are shown in the top three rows, and the standardized beta weights for each predictor in subsequent rows. Significant beta weights are indicated at \*p < .05; and \*\*p < .001

|  | HVLT:<br>Delayed<br>Recall | HVLT:<br>Discrim.<br>Index | Digit<br>Span:<br>Forward | Digit<br>Span:<br>Back | Category<br>Switch:<br>% Trials | Category<br>Switch:<br>RT | Dig Sub:<br>Number<br>Correct | Dig Sub:<br>Sec Per<br>Correct | Stop<br>Signal:<br>Pro Reac | Stop<br>Signal:<br>RT |
| --- | --- | --- | --- | --- | --- | --- | --- | --- | --- | --- |
| ANOVA F | 2.4 | 4.7 | 0.3 | 1.7 | 2.6 | 6.3 | 17.2 | 0.9 | 1.4 | 2.3 |
| ANOVA p | 0.028 | <.001 | 0.948 | 0.113 | 0.018 | <.001 | <.001 | 0.529 | 0.230 | 0.038 |
| Variance Explained | 18% | 30% | 3% | 14% | 19% | 36% | 61% | 8% | 11% | 17% |
| fPET: Global Efficiency | 0.09 | 0.66* | 0.13 | 0.08 | 0.23 | -0.19 | -0.21 | -0.42 | -0.31 | 0.25 |
| fPET: Local Efficiency | 0.20 | 0.67* | 0.03 | -0.16 | 0.33 | -0.21 | -0.14 | -0.38 | -0.04 | 0.37 |
| fPET: Betweenness Centrality | -0.09 | -0.24 | -0.02 | -0.01 | -0.14 | 0.02 | 0.06 | 0.12 | 0.29* | -0.02 |
| fMRI: Global Efficiency | -0.24 | -0.16 | 0.09 | 0.17 | -0.29 | 0.08 | -0.14 | -0.12 | -0.01 | -0.15 |
| fMRI: Local Efficiency | -0.27 | 0.04 | -0.12 | -0.20 | -0.35 | -0.27 | -0.15 | 0.00 | -0.05 | -0.22 |
| fMRI: Betweenness Centrality | 0.01 | -0.07 | 0.15 | 0.26 | 0.28 | 0.20 | 0.11 | 0.01 | 0.01 | -0.19 |
| Age Group | -0.35* | -0.37* | 0.08 | -0.19 | -0.17 | -0.55* | -0.75** | -0.19 | -0.08 | -0.23 |

Discrim = discrimination; Dig Sub = Digit Substitution; Pro React = probability of reacting.

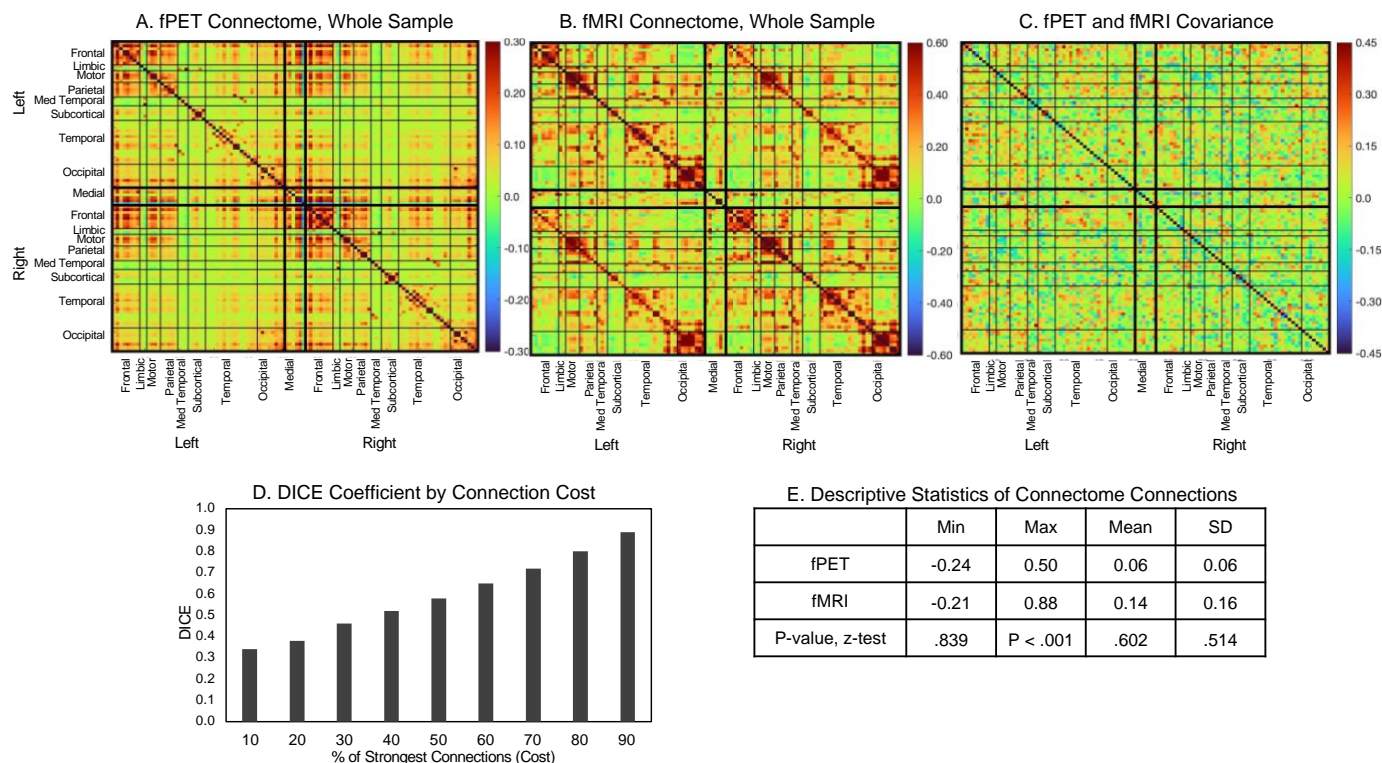

Supplementary Figure 1. fPET and fMRI connectomes and covariance among the whole sample derived from the Harvard Oxford Atlas parcellation. (A) fPET connectome, (B) fMRI connectome, (C) across-subject covariance of fPET and fMRI connectomes, (D) DICE coefficients at connection costs, and (E) descriptive statistics and z-test of modality differences.

It is noteworthy that the maximum connections in the fPET connectomes derived from the Harvard Oxford Atlas (Supplementary Figure 1) were higher than in the Schaefer (max $r = .50$) but were similar for the fMRI connectomes (max $r = .88$). The DICE coefficient was .58 for the top 50% of connections and .34 at top 10% of edges.

The fPET connectome has a relatively homogenous connection strength (SD = .06) within and between networks, mostly in the order of $r = .20$ to $0.40$ (Supplementary Figure 1A). In contrast, although not statistically significant, the fMRI connectome is more heterogenous in strength (SD = .16). The highest correlations in the fMRI are larger than the fPET connections, particularly the within-lobe network connections (Supplementary Figure 1B). The strongest covariance for appears to be in between-lobe connections, particularly for regions between the frontal, motor, parietal and occipital cortices (Supplementary Figure 1C).

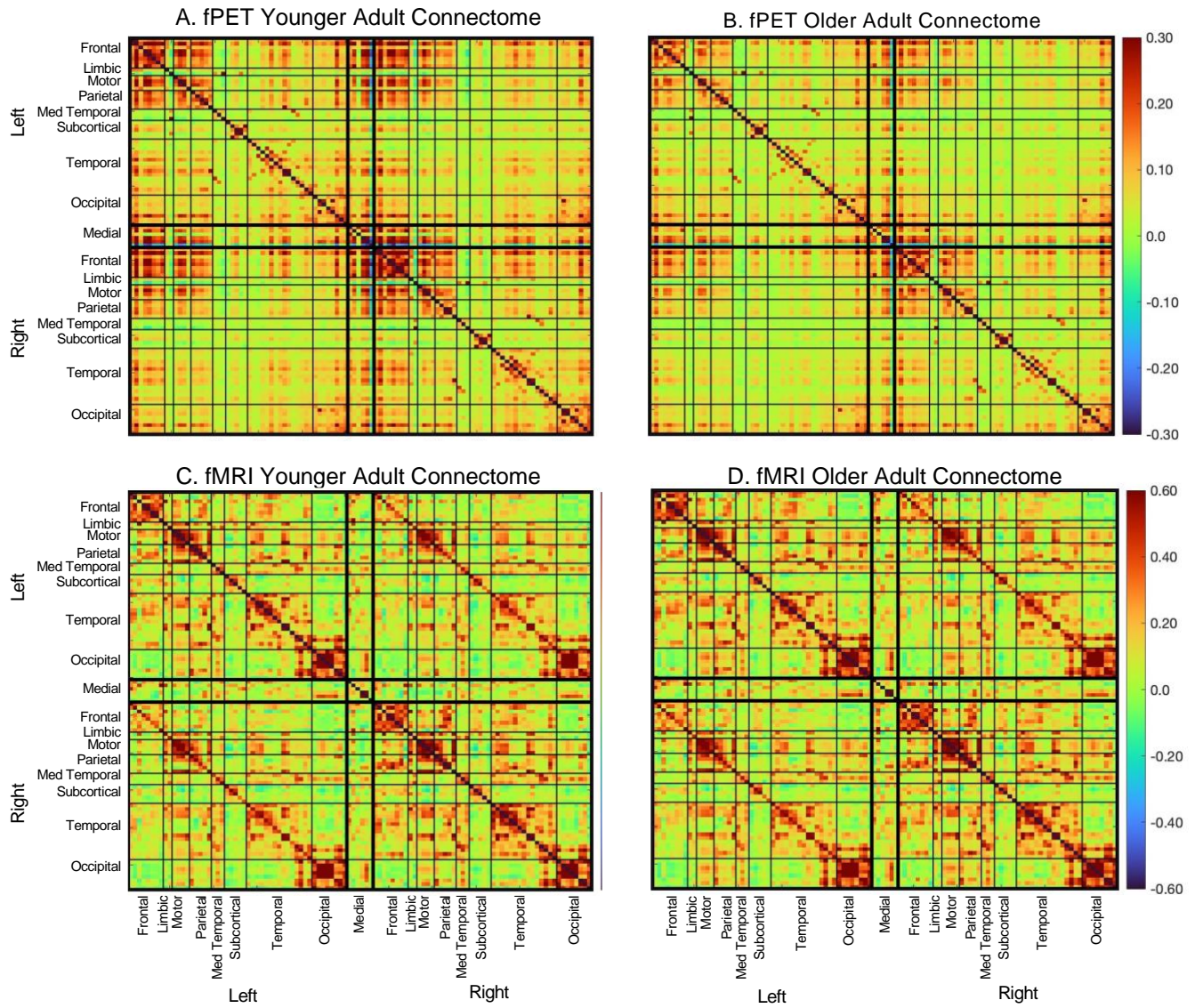

Supplementary Figure 2. fPET and fMRI connectivity for younger and older adults from the Harvard Oxford Atlas parcellation. (A) Younger and (B) older adult fPET metabolic connectomes, and (C) younger and (D) older adult fMRI connectomes.

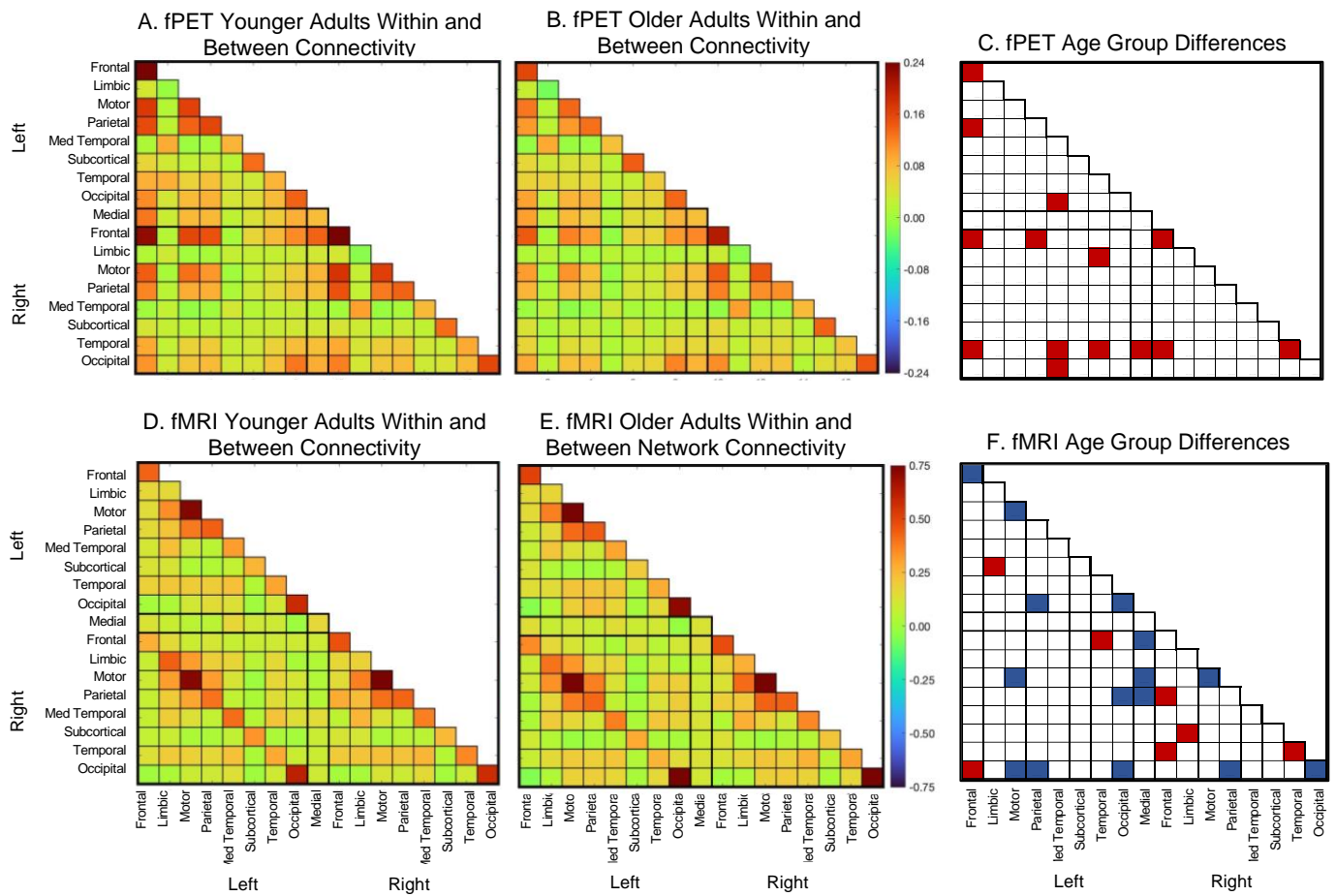

Supplementary Figure 3. Within and between connectivity averaging the nodes of the Harvard Oxford Atlas within the lobes for (A) younger and (B) older adults using fPET, and (D) Younger and (E) older adults in fMRI. Within connectivity is shown in the diagonal cells and between connectivity on the off-diagonal cells of each matrix. (significance test ($t$-test, $df = 84$) of younger vs older adults, with shaded cells indicating a statistically significant age group differences at $p$-FDR<.05 ((C) and (F)). Red shading younger > older; blue shading older > younger.

The younger adult fPET within-network connectivity Younger mean = .12, SD = .07 and between-network mean = .06, SD = .04. Older adult fPET within-network mean = .10, SD = .06 and between-network mean = .04, SD = .04. Younger adult fMRI within-network mean = .40, SD = .18 and between-network mean = .14, SD = .15. Older adult fMRI within-network mean = .43, SD = .19 and between-network mean = .15, SD = .15

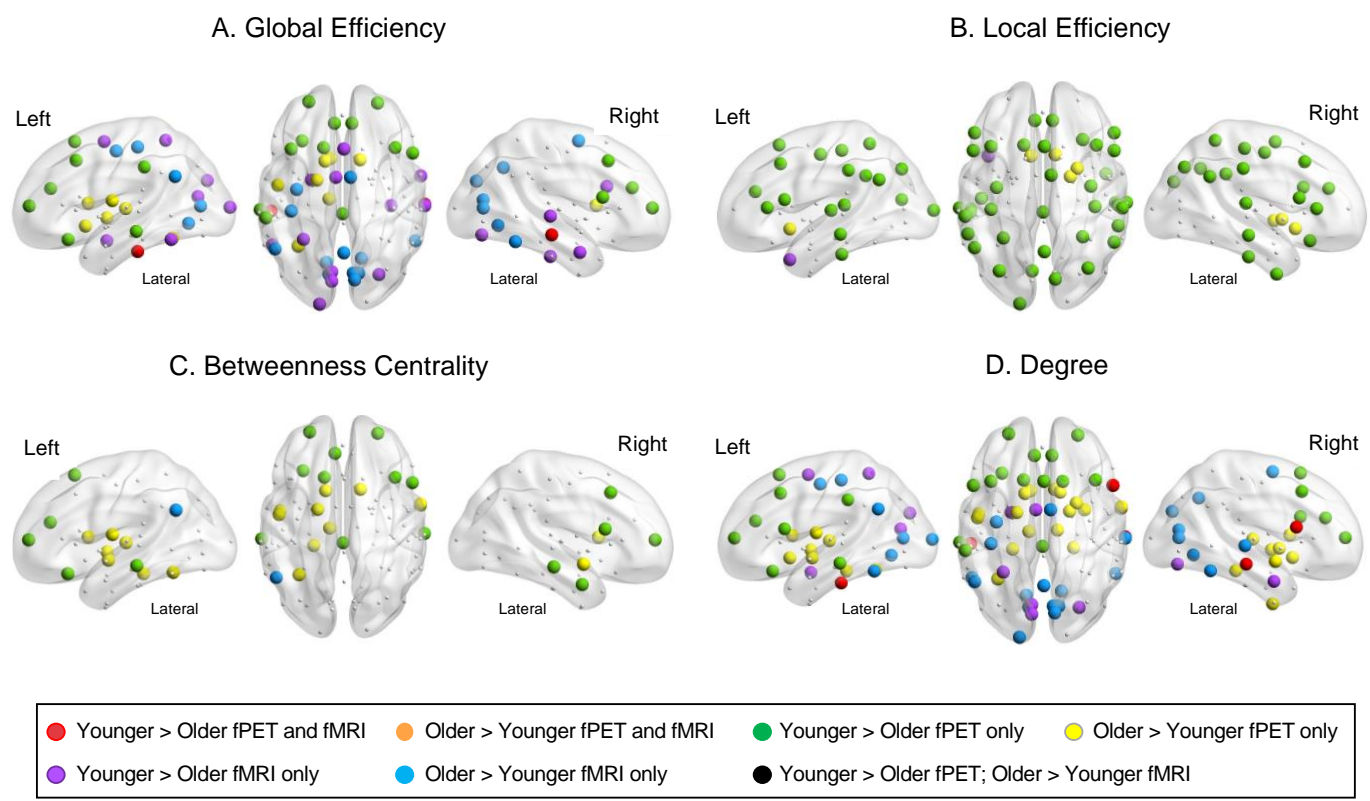

Supplementary Figure 4. fPET and fMRI age group differences in regions of the Harvard Oxford Atlas for (A) global efficiency, (B) local efficiency, (C) betweenness centrality, and (D) degree. Regions are shown as colored dots where there is a statistically significant age group difference at $p\text{-FDR} < .05$ from one-sided $t$-tests. For each region, the mean, standard deviations, effect sizes and $t$-tests are shown in Supplementary Tables 7 to 10. FR = frontal; LIM = limbic; SM = motor; PAR = parietal; MEDTP = medial temporal; SUB = subcortical; TEMP = temporal; OCC = occipital; MED = medial.

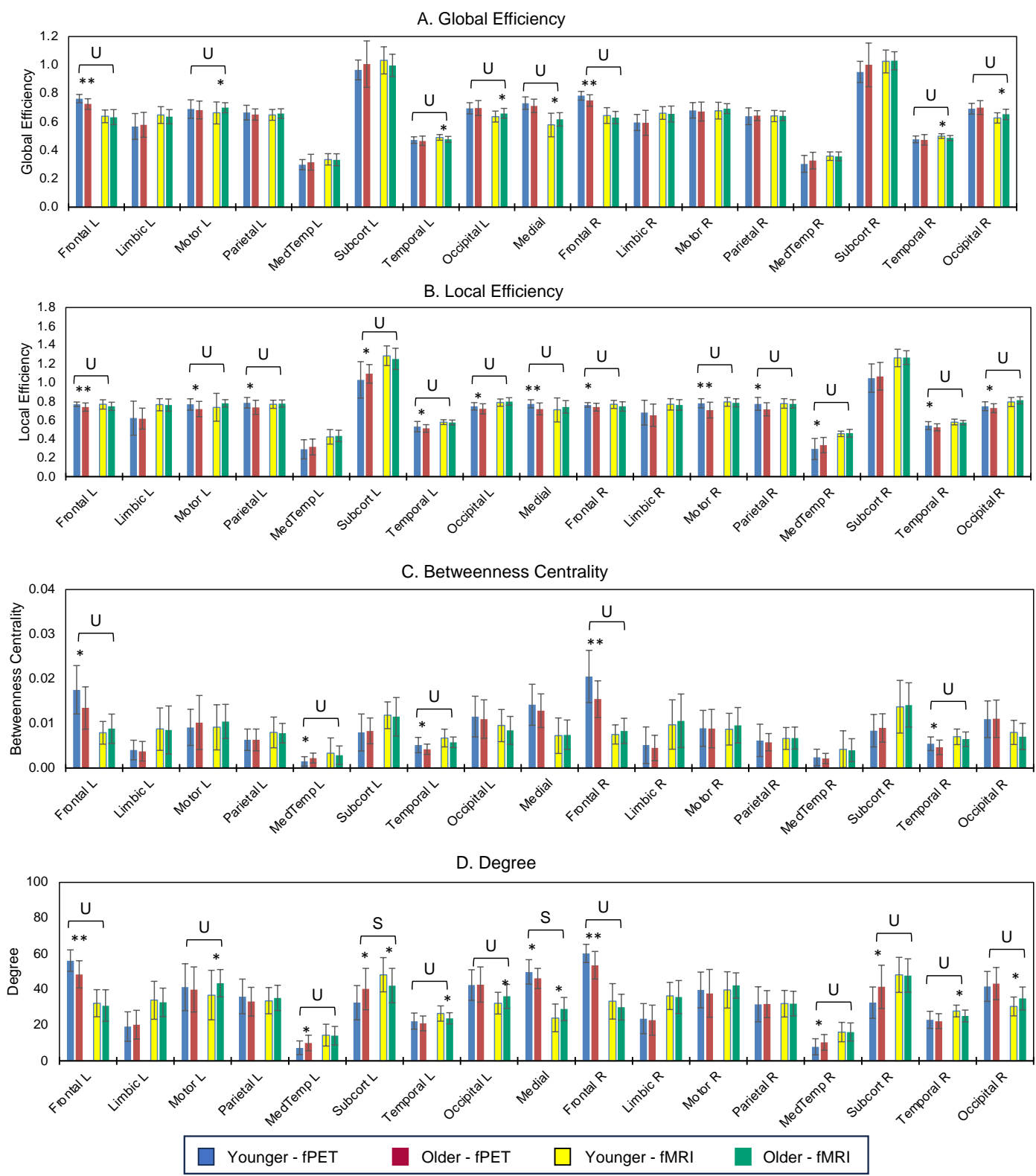

Supplementary Figure 5. Mean and standard deviation (error bars) of fPET and fMRI graph metrics for younger and older adults from the Harvard Oxford Atlas. Age group differences at \*P-FDR < .05 and \*\*p-FDR < .001 from one-sided *t*-tests; S = Age group differences, same direction in both modalities; U = unique age group differences in one modality; D = different age group direction in one modality vs the other.

### Relative Predictive Strength of fPET and fMRI Graph Metrics for Age Group

The discriminant function analysis was significant for the fPET whole brain metrics (Wilk's Lambda = .837, Chi-square = 14.7, $p < .002$) but not the fMRI whole brain graph metrics (Wilk's Lambda = .955, Chi-square = 3.8, $p = .285$). The probability of a correct age group classification was 57% and 59% for younger and older adults in the fPET, respectively; and 52% and 52% for younger and older adults in the fMRI.

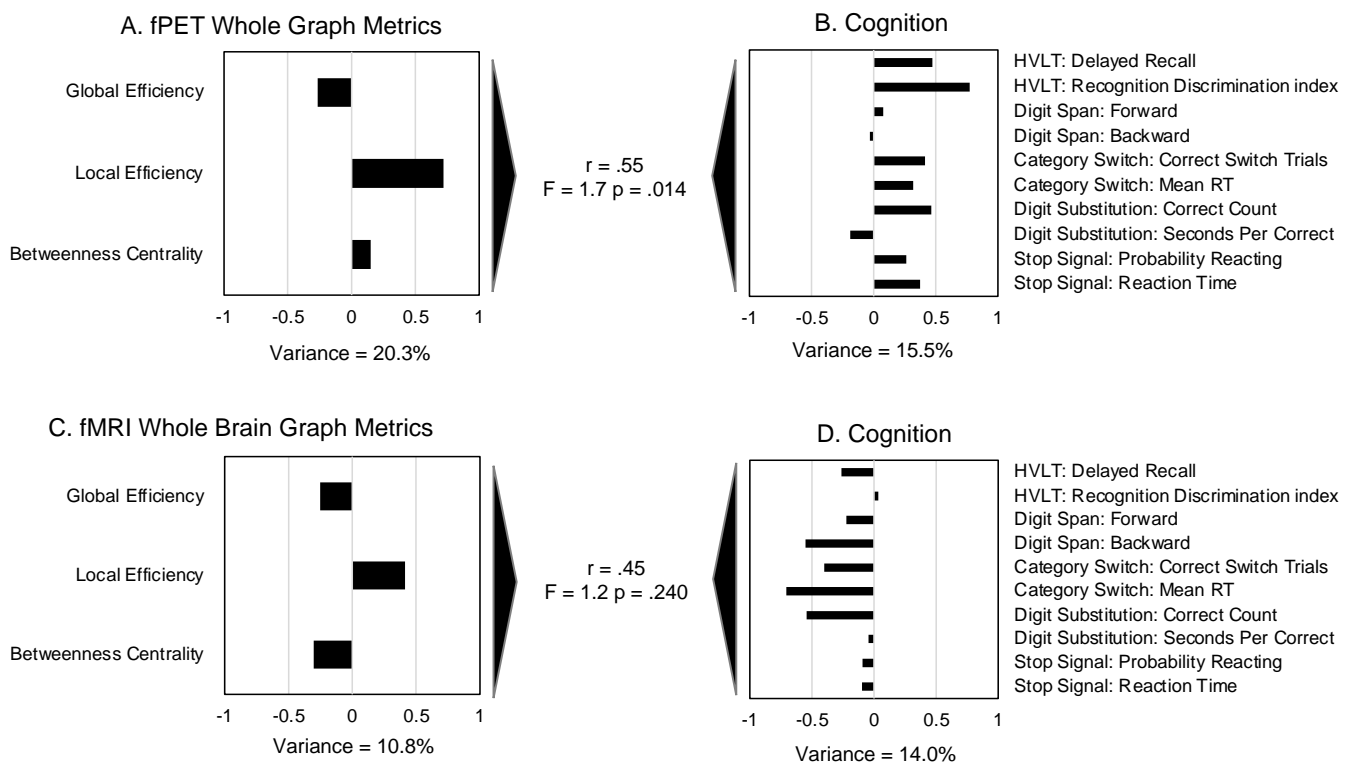

Supplementary Figure 6. Canonical correlations between the whole brain graph metrics from the Harvard Oxford Atlas for fPET (A) and fMRI (C) and cognition ((B) and (D)). $r$-value is the canonical correlation between the linear combinations of the graph metrics and cognition variables that maximally covary across subjects. $F$-statistic is Wilk's test of the null hypothesis that the canonical correlation and all smaller ones are equal to zero and was significant for one canonical variate for fPET but not fMRI. The correlations on each variable set represent the strength of the association between the variable and the canonical variate. Variance explained is the percentage of variance explained by the variables in their variate. Stop signal reaction time, seconds per correct response in the digit substitution and category switch reaction time were multiplied by -1 so that higher scores reflect better performance. The test of difference between two correlations = 0.86; $p = .389$.

1. Shapiro, A.M., et al., *Construct and concurrent validity of the Hopkins Verbal Learning Test-revised*. Clin Neuropsychol, 1999. **13**(3): p. 348-58.
2. Blackburn, H.L. and A.L. Benton, *Revised administration and scoring of the digit span test*. J Consult Psychol, 1957. **21**(2): p. 139-43.
3. Friedman, D., et al., *Age-related changes in executive function: an event-related potential (ERP) investigation of task-switching*. Neuropsychol Dev Cogn B Aging Neuropsychol Cogn, 2008. **15**(1): p. 95-128.
4. Verbruggen, F., G.D. Logan, and M.A. Stevens, *STOP-IT: Windows executable software for the stop-signal paradigm*. Behav Res Methods, 2008. **40**(2): p. 479-83.
5. Thorndike, E.L., *A Standardized group examination of intelligence independent of language*. Journal of Applied Psychology, 1919. **31**: p. 13-32.
6. Jamadar, S.D., et al., *Simultaneous BOLD-fMRI and constant infusion FDG-PET data of the resting human brain*. Sci Data, 2020. **7**(1): p. 363.
7. Greve, D.N., et al., *Different partial volume correction methods lead to different conclusions: An (18)F-FDG-PET study of aging*. Neuroimage, 2016. **132**: p. 334-343.
8. Greve, D.N., et al., *Cortical surface-based analysis reduces bias and variance in kinetic modeling of brain PET data*. Neuroimage, 2014. **92**: p. 225-36.
9. Schaefer, A., et al., *Local-Global Parcellation of the Human Cerebral Cortex from Intrinsic Functional Connectivity MRI*. Cereb Cortex, 2018. **28**(9): p. 3095-3114.
10. Eickhoff, S.B., R.T. Constable, and B.T.T. Yeo, *Topographic organization of the cerebral cortex and brain cartography*. Neuroimage, 2018. **170**: p. 332-347.
11. Di, X., B.B. Biswal, and I. Alzheimer's Disease Neuroimaging, *Metabolic brain covariant networks as revealed by FDG-PET with reference to resting-state fMRI networks*. Brain Connect, 2012. **2**(5): p. 275-83.
12. Savio, A., et al., *Resting-State Networks as Simultaneously Measured with Functional MRI and PET*. J Nucl Med, 2017. **58**(8): p. 1314-1317.
13. Lodge, M.A., A. Rahmim, and R.L. Wahl, *Simultaneous measurement of noise and spatial resolution in PET phantom images*. Phys Med Biol, 2010. **55**(4): p. 1069-81.
14. Yeo, B.T., et al., *The organization of the human cerebral cortex estimated by intrinsic functional connectivity*. J Neurophysiol, 2011. **106**(3): p. 1125-65.
15. Deery, H.A., et al., *Lower brain glucose metabolism in normal ageing is predominantly frontal and temporal: A systematic review and pooled effect size and activation likelihood estimates meta-analyses*. Hum Brain Mapp, 2023. **44**(3): p. 1251-1277.
16. Deery, H.A., Liang, E., Siddiqui, M. N., Murray, G., Voigt, K., Di Paolo, R., Moran, C., Egan, G.F., and Jamadar, S.D., *Metabolic Connectivity in Ageing*, in *BioRxiv*. 2024.
